# Food-Poisoning Bacteria Employ a Citrate Synthase and a Type II NRPS to Synthesize Bolaamphiphilic Lipopeptide Antibiotics

**DOI:** 10.1101/2020.06.26.173229

**Authors:** Benjamin Dose, Claudia Ross, Sarah P. Niehs, Kirstin Scherlach, Johanna P. Bauer, Christian Hertweck

## Abstract

Mining the genome of the food-spoiling bacterium *Burkholderia gladioli* pv. *cocovenenans* revealed five non-ribosomal peptide synthetase (NRPS) gene clusters, including an orphan gene locus *(bol).* Gene inactivation and metabolic profiling linked the *bol* gene cluster to novel bolaamphiphilic lipopeptides with antimycobacterial activity. A combination of chemical analyses and bioinformatics elucidated the structures of bolagladin A and B, lipocyclopeptides featuring an unusual dehydro-β-alanine enamide linker fused to an unprecedented tricarboxylic fatty acid tail. Through a series of targeted gene deletions we proved the involvement of a designated citrate synthase (CS), priming ketosynthases (KS III), a type II NRPS including a novel desaturase for enamide formation, and a multimodular NRPS generating the cyclopeptide. Network analyses revealed the evolutionary origin of the CS and identified cryptic CS/NRPS gene loci in various bacterial genomes.

---

**A**mong the many food-spoiling bacteria, one species stands out, as it is responsible for the governmental ban of a national dish: *Burkholderia gladioli* pv. *cocovenenans* (literally meaning *making coconut poisonous).* These bacteria poison the Indonesian specialty tempeh bongkrek, a coconut press cake produced by fermentation using a harmless mold fungus *(Rhizopus).* Yet, frequent contaminations of the *Rhizopus* cultures with bacterial toxin producers have been responsible for the deaths of thousands that have consumed the delicacy.^[1]^ The causative agent of these intoxications has been identified to be the respiratory toxin bongkrekic acid.^[1]^ To gain insight into its biosynthesis, we have sequenced the genome of *B. gladioli* pv. *cocovenenans*. In addition to the molecular basis of bongkrekic acid biosynthesis, we discovered various other biosynthesis gene clusters.^[2]^ Yet, surprisingly little is known about the biosynthetic potential of the food-poisoning bacteria, in particular with respect to peptidic natural products. Here, we deduce orphan non-ribosomal peptide synthetase (NRPS)^[3]^ gene clusters of *B. gladioli* pv. *cocovenenans*, and report the surprising discovery of a new family of antibiotics, non-canonical lipopeptides equipped with an unprecedented tricarboxylic acid tail that results from the recruitment of a citrate synthase.

Genome mining^[4]^ of *B. gladioli* pv. *cocovenenans* revealed the presence of five orphan NRPS gene clusters. Bioinformatics predictions and BLAST homology search indicated that four of these gene clusters code for the biosynthesis of characterized lipopeptides, namely haereogladins (promoting biofilm formation), burriogladins (facilitating swarming), and icosalides (surfactant and swarming inhibitor) (Figure 1A).^[5]^ Furthermore, one NRPS gene cluster codes for the rare diazeniumdiolate siderophore gladiobactin.^[6]^ We verified the predicted functions by HPLC-HR-MS of culture extracts and MS/MS fragmentation.^[6]^ The fifth cryptic biosynthetic gene cluster (named *bol*) consists of a central 13 kb NRPS gene *(bolA)* that is flanked by a number of putative accessory genes *(bolB–Y)* (Figure 1B). By means of database searches we noted that genomes of numerous other *B. gladioli* isolates from highly diverse niches harbor orthologous, yet cryptic, gene clusters (Supporting Information).

**Figure 1:**
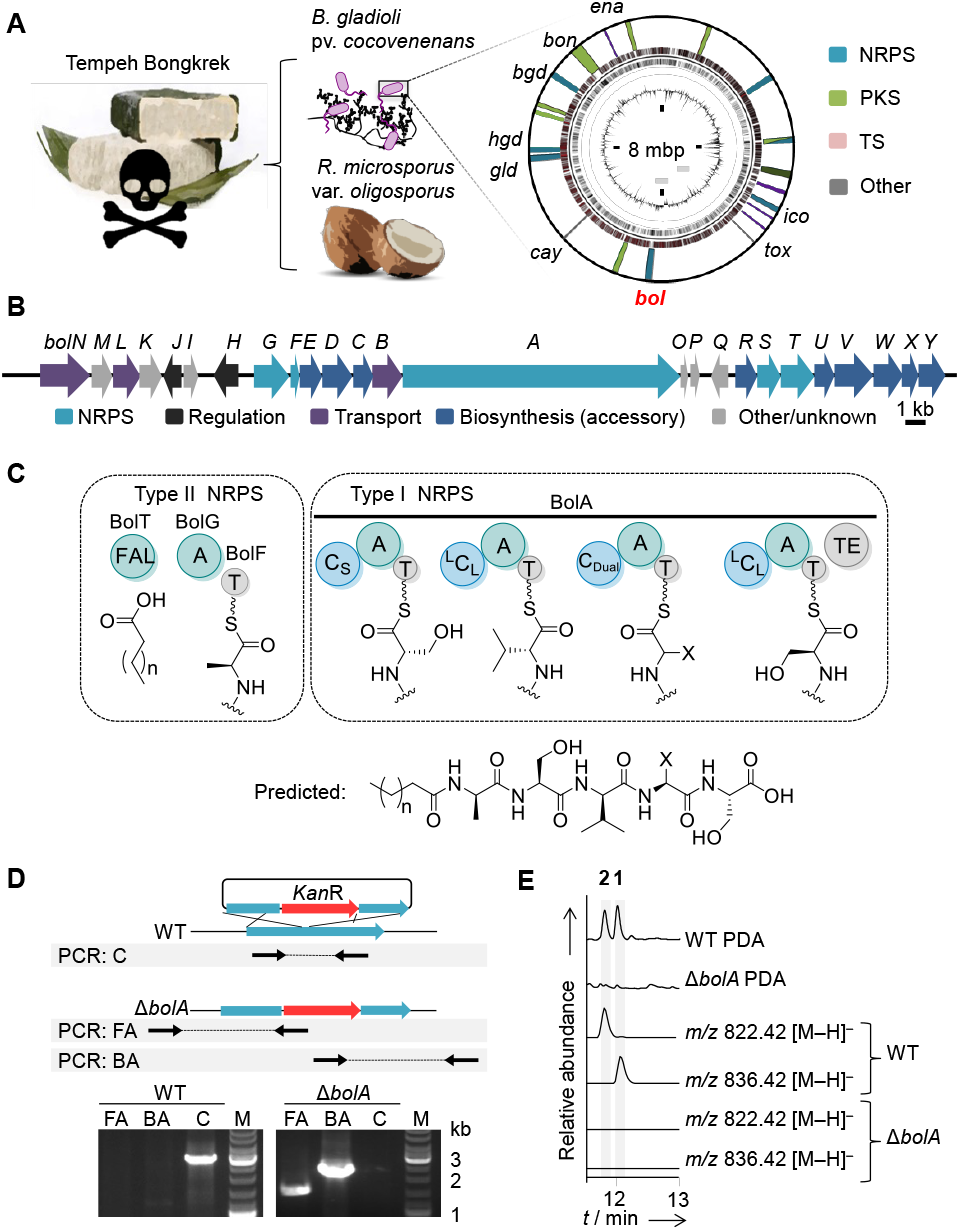
Genome mining of the bacterial contaminant of tempeh bongkrek. A) Biosynthetic potential encoded in the chromosome. Gene loci for known compounds: *ena,* enacyloxin; *bon*, bongkrekic acid; *cay*, caryoynencin; *tox,* toxoflavin; NRPS: *bgd*, burriogladin; *hgd*, haereogladin; *glb*, gladiobactin; *ico*, icosalide*; bol*, cryptic NRPS gene locus, marked in red. B) Organization of the *bol* gene cluster. C) Encoded assembly line and predicted peptide backbone. FAL, fatty acid ligase; A, adenylation domain; T, thiolation domain or peptide carrier protein; C_S_, starter condensation domain; ^L^C_L_, C domain fusing two L-amino acids; C_dual_, C domain also promoting epimerization; TE, thioesterase domain. D) Mutant construction using homologous recombination, and PCR-based verification of *ΔbolA* mutant. E) Production of **1** and **2** in *B. gladioli* pv. *cocovenenans* (WT), and metabolic profile of deletion mutant *ΔbolA*.

To correlate the *bol* gene cluster with metabolites we deduced the core peptide structure from *in silico* analyses and performed comparative metabolic profiling of the wild type and a targeted null mutant. First, we gleaned the architecture of the cryptic peptide from bioinformatics analysis of the *bol* assembly line. In brief, the encoded type I NRPS (BolA) consists of four modules each composed of condensation (C), adenylation (A) and thiolation (T) domains. Furthermore, genes for a type II NRPS^[7]^ comprising freestanding adenylation (A, BolG) and peptidyl carrier protein (PCP, BolF) domains, and a fatty acid ligase (FAL, BolT) suggest that an activated *N*-acyl-amino acid would be merged with a tetrapeptide, likely yielding a lipopeptide with five amino acid residues. The detailed analysis of the A and C domain specificities,^[8]^ where possible, predicted that an *N*-acyl-L-Ala-derived building block would be linked to an l-Ser-d-Val-l-X-l-Ser chain (Figure 1C).

To create a null producer, we improved the current protocol for the genetic manipulation of *B. gladioli* pv. *cocovenenans.* We noted that this strain is naturally competent to take up DNA and achieved the targeted gene knockout of the NRPS gene by means of homologous recombination and integration of a kanamycin resistance cassette into *bolA,* using a double-selection plasmid containing the *pheS* gene.^[9]^ By antibiotic selection and PCR analysis the desired mutation was confirmed (Figure 1D). By comparison of the metabolic profiles of wild type and *ΔbolA* mutant grown under various conditions we detected two metabolites (**1** and **2**) in the wild-type culture that are absent in the mutant strain. HPLC-HRMS elucidated two major congeners with *m/z* 836.4322 [M–H]^-^ (**1**) and *m/z* 822.4168 [M–H^-^ (**2**) (Figure 1D). The deduced molecular formulas and the MS/MS fragmentation patterns suggested that **1** and **2** represent the predicted lipopentapeptides.

To determine their structures, we optimized culture conditions and subjected the up-scaled fermentation broth (3 L) to chromatographic separation by size-exclusion chromatography and preparative HPLC, yielding pure **1** (1.6 mg) and **2** (0.8 mg). The structures of **1** and **2** were fully elucidated by ESI-HR-MS, MS/MS, and 1D and 2D NMR experiments (Figure 2A). For **1** the molecular formula of C_40_H_63_N_5_O_14_ was deduced from ESI-HR-MS, and the number of carbon atoms was corroborated by ^13^C and HMBC NMR data. The presence of a fatty acid residue was confirmed by analysis of ^1^H NMR and DEPT-135 NMR data. From ^1^H,^1^H-COSY couplings the spin systems of five amino acids were deduced (Figure 2A). ^1^H,^13^C HMBC couplings of the NH protons of each amino acid with the corresponding carbonyl carbon atoms and the respective Cα atoms of the neighboring amino acid allowed the assignment of the peptide chain. An ^1^H,^13^C-HMBC coupling of the β-proton of the dehydroalanine unit with C-1 of the fatty acid revealed the *N*-acyl connection. Comparison of the deduced molecular composition (C39H61N5O14) and the MS/MS fragmentation of **2** with the ones determined for **1** indicated a Val to Ile replacement in **2**. 1D and 2D NMR analyses confirmed this assignment. The absolute configuration of the amino acids in **1** and **2** was unambiguously determined by Marfey’s analysis^[10]^ (see Supporting Information).

**Figure 2.**
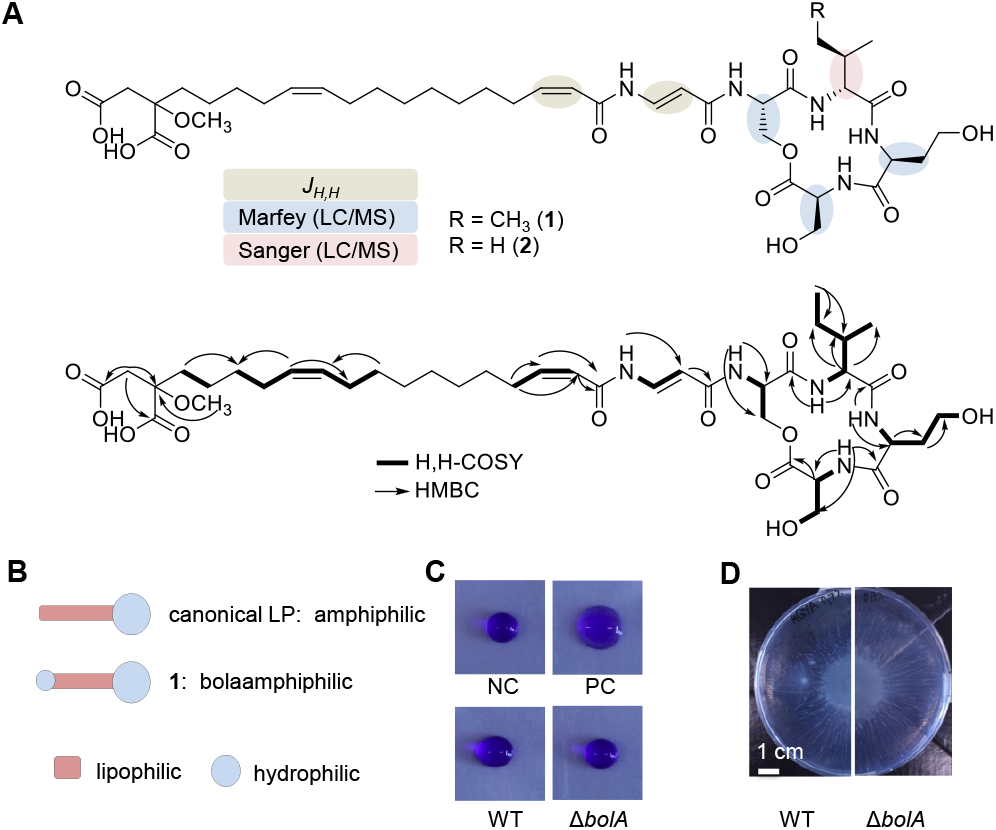
Structures and bioactivities of bolagladins. A) Structure elucidation by 2D NMR couplings and Marfey’s and Sanger’s reagents. B) Schematic representation of canonical lipopeptides (LPs) *vs.* bolagladin. C) Droplet collapse assay showing no tensioactive property of **1**, NC, negative control: water; PC, positive control, tenside producer; WT, wild type; *ΔbolA,* mutant. D) Swarming areas of WT mutant on semisolid agar.

The structure of the fatty acid moiety proved to be more complex than anticipated. MS/MS analyses revealed a fragment ion with *m/z* 450.2480, indicating an identical, yet highly substituted fatty acid residue (C24H36NO7) in **1** and **2**. ^1^H,^1^H-COSY couplings of the protons H-2 to H-4 and H-9 to H-14 and 1D NMR signals for additional methylene functions established the fatty acid chain. A coupling constant of *J_2,3_ =* 11 Hz indicated the *Z* configuration of the double bond. ^1^H,^13^C-HMBC couplings (Figure 2A) finally unveiled the presence of an unusual methoxy dicarboxylic acid terminus.

The structures of **1** and **2** are highly unusual. The rare dehydro-β-Ala linker is known from the *Streptomyces-derived* enamidonins, yet their biosynthesis has remained enigmatic.^[11]^ Furthermore, owing to the unprecedented tricarboxylic fatty acid tail, **1** and **2** constitute a novel family of lipopeptides. Regular lipopeptides consist of a hydrophilic head moiety (linear or cyclic peptide) and a hydrophobic fatty acid tail.^[12]^ The resulting amphiphilic nature of lipopeptides is usually reflected in their bioactivities, which encompass surfactant properties that may influence biofilm formation and swarming of the producer, and may disintegrate membranes of recipients (Figure 2B).^[12b, 13]^ Compounds **1** and **2** clearly deviate from the canonical lipopeptide architecture and polarity as they have two hydrophilic groups at both ends of a hydrophobic chain and thus are natural bolaamphiphiles.^[14]^ Therefore, we named them bolagladin A (**1**) and B (**2**).

The markedly poor solubility of the bolagladins hampered the routine profiling the biological activities of the isolated compounds. Thus, we first compared typical lipopeptide activities of *B. gladioli* pv. *cocovenenans* wild type and the *ΔbolA* mutant. We found that the mutant does not show altered tensioactive properties in a droplet-collapse assay, and when grown on low-density agar its swarming behavior is similar to the wild type (Figure 2 C and D). Notably, the strains do not produce any of the known tensioactive lipopeptides under these cultivation conditions. Next, we evaluated potential metal-binding properties of **1**, since *bolN* could code for a TonB-like receptor that is typically associated with siderophores. However, strong metallophore activities were not observed in a CAS agar assays. Furthermore, stable iron adducts of **1** were not detected by mass spectrometry. Finally, we subjected **1** to antimicrobial assays using a panel of bacterial and fungal strains. We observed moderate antifungal activities of **1** against *Candida albicans* and *Penicillium notatum*, moderate activities against St*aphylococcus aureus* (MRSA) and *Enterococcus faecalis* (VRE), and more pronounced activity against *Mycobacterium vaccae* (Supporting Information).

As to the most unusual feature of the bolagladins, their branched tricarboxylic fatty acid tail, we predicted that an encoded citrate synthase (BolW), an acyl-CoA ligase (BolT), an *O*-methyl transferase (BolX), and two encoded priming ketosynthases (KS III, BolR, BolU) play a key roles in the its formation. A phylogenetic analysis of BolW and a selection of citrate synthases (CS) and related biocatalysts showed that BolW falls into the group of CSs and related enzymes that promote an aldol reaction at the *Si-*face of 2-oxoglutarate (Figure 3A).^[15]^ Close relatives of BolW include the CSs from *B. gladioli* primary metabolism and CSs from the biosynthetic pathways of squalestatins^[16]^ and maleidrides.^[17]^

**Figure 3.**
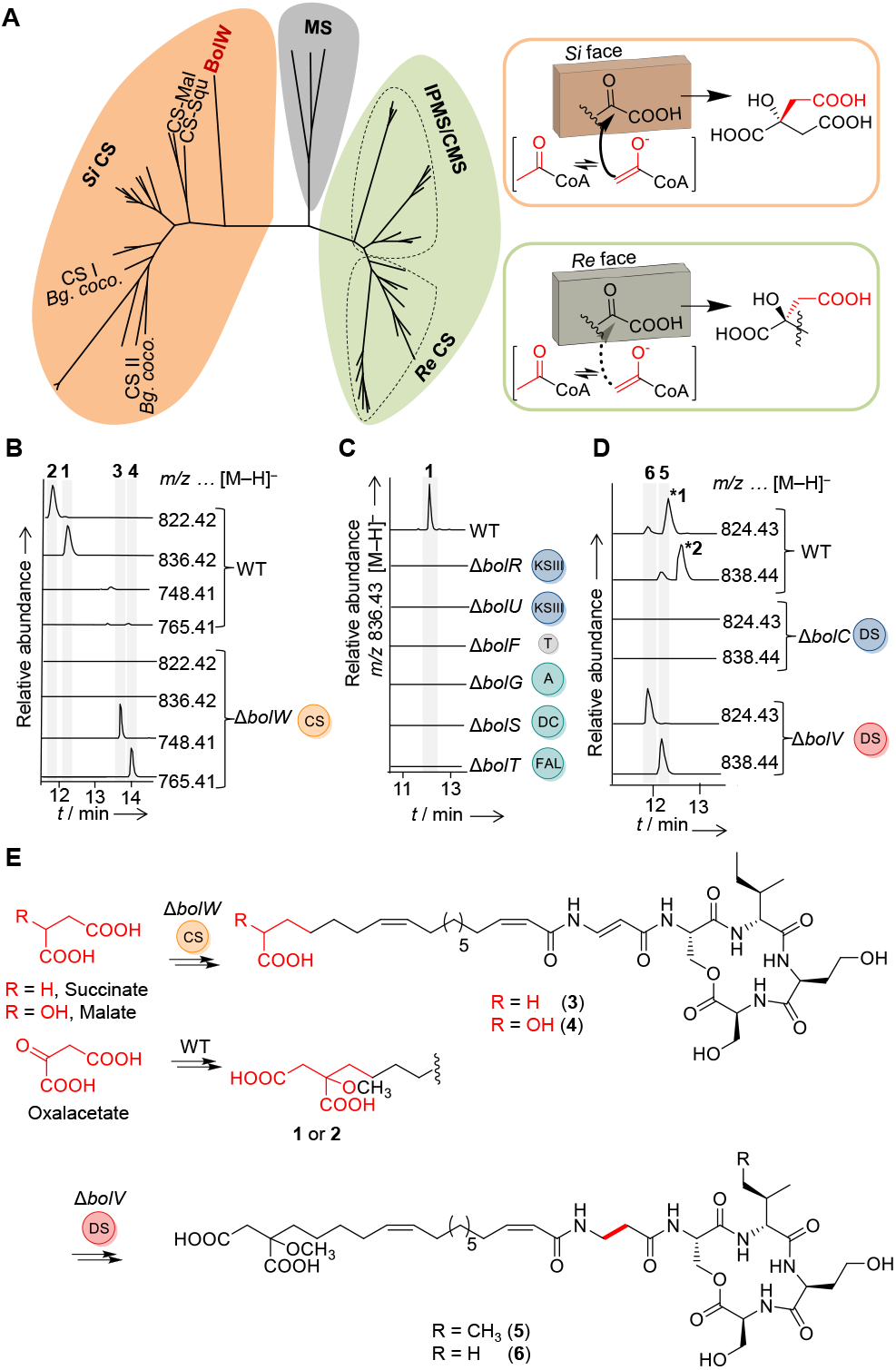
Phylogenetic and mutational analysis of key pathway enzymes. A) Cladogram of *Si- and Re*-face-specific citrate synthases *(Si/Re* CS, CSI/CSII) from primary metabolism, squalestatin CS, maleidrides CS, isopropylmalate and *Re*-citramalate synthase (IPMS and CMS). Outgroup: malate synthase (MS). B–D) Metabolic profiling of targeted mutants. \***1** and \***2**, Isotopes of **1** and **2.** E) Structures of **3** and **4** formed in Δ*bolW* mutant, likely resulting from incorporation of succinate and malate in lieu of citrate, and structures of **5** and **6** formed in Δ*bolV* mutant.

To verify the role of BolW, we created a null mutant by double crossover. Metabolic profiling of the resulting *ΔbolW* mutant culture by HPLC-HRMS revealed that the production of **1** and **2** was fully abrogated. Instead, we detected two new metabolites with *m/z* 748.4157 [M–H]^-^ (**3**) and 765.4120 [M–H]^-^ (**5**) that are absent in the wild-type metabolome (Figure 3B). Their deduced molecular formulas (**3**: C_36_H_56_N_5_O_11_, **4**: C_37_H_58_N_5_O_12_), MS/MS analyses and NMR data of **3** revealed that, in comparison to **1**, both compounds are devoid of the terminal carboxymethyl moiety. In addition, **3** lacks the methoxy group (Figure 3E). A plausible explanation for the production of **3** and **4** in the *ΔbolW* mutant would be the incorporation of succinate and malate building blocks in lieu of citrate (Figure 3E). Apparently, the *O*-methyl transferase BolX does not recognize the alternative substrate. Mutational analyses showed that both encoded KS III (BolR and BolU), and the ligases (BolG) are essential for *bol* biosynthesis, too (Figure 3C). Since none of the congeners could be detected in these mutants, we propose that the function of the ligases and KS III is the activation and elongation of citrate and the fatty acid, respectively. The tricarboxylic fatty acid would then be activated by the fatty acid ligase BolT and transferred onto the type II NRPS to form the acyl-β-dh-Ala subunit.

Initially, *N*-acyl-L-Asp, bound to the PCP (BolF), would be assembled by means of the freestanding A domain BolG. L-Asp-PCP would be transformed into β-Ala-PCP by means of a pyridoxal phosphate-dependent decarboxylase, BolS, in analogy to the destruxin and fluvirucin biosynthetic pathways.^[18]^ Finally, a desaturase would introduce the enamide double bond. We confirmed this model by targeted gene inactivation of *bolF, bolG, bolS* and *bolT* (Figure 3C). To pinpoint the desaturase in charge of the unusual peptide modification we generated and profiled a range of candidate mutant cultures *(ΔbolQ, ΔbolC,* and *ΔbolV).* While the *ΔbolQ* mutant showed the wild type profile, in the *ΔbolC* and *ΔbolV* mutants production of **1** and **2** was fully abrogated. In the *ΔbolV* mutant we identified congeners with *m/z* 838.4484 [M–H]^-^ (**5**) and 824.4327 [M–H]^-^ (**6**) lacking the enamide double bond (Figure 3E). Thus, BolV seems to be a novel *trans*-acting desaturase that generates the unusual enamide substructure online. Finally, the rare β-dh-Ala brick would be transferred onto the modular NRPS to assemble the full-length peptide (Figure 4A).

To gain insight into the evolution and distribution of similar biosynthetic pathways we created sequence similarity networks (SSNs) and genome neighborhood networks (GNNs) using CS I and BolW as the query sequences (Figure 4B).^[19]^ The CS I network forms a single cluster that places the CS gene exclusively among genes associated with primary metabolism in Proteobacteria. The BolW-based network, in contrast, shows several clusters and revealed a high synteny of orthologous genes for citrate synthase (BolW), KS III (BolR and BolU), the fatty acid CoA ligase (BolG) and PCP (BolF) all putatively involved in the biosynthesis, activation and transfer of a citrate building block onto the assembly line. Our BolW network analysis led to the discovery of cryptic NRPS gene clusters that could code for the biosynthesis of related tricarboxylic fatty acid-derived lipopeptides in *Burkholderia cepac*ia and *Burkholderia stabilis.* Furthermore, we found NRPS-independent biosynthetic pathways in genomes of *Burkholderia, Pseudomonas, Collimonas*, and *Chromobacterium* species.

**Figure 4.**
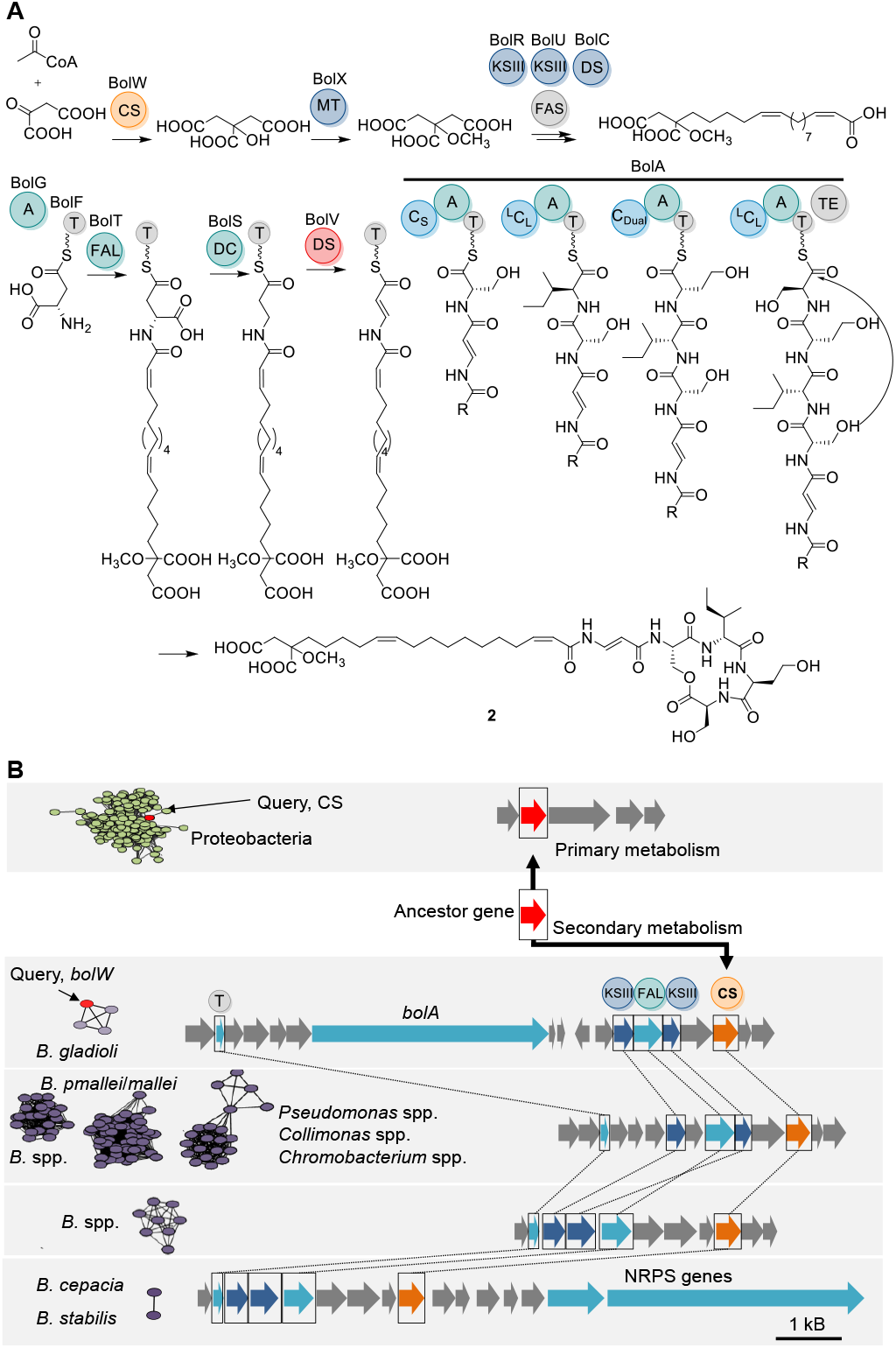
Model of bolagladin biosynthesis, and network analysis for genome mining. A) Biosynthesis of the tricarboxylic fatty acid involving citrate synthase (CS) BolW, MT, methyltransferase; KS III, priming ketosyntase; FAS, fatty acid synthase; DC, decarboxylase; Des, desaturase. B) Sequence similarity network and genome neighborhood networks identify the evolutionary origin of BolW and enable the discovery of cryptic CS/NRPS gene clusters.

Beyond genome mining for novel natural products, the SSNs and GNNs shed light on the origin of BolW. While regular CS and their surrounding genes are highly conserved in Proteobacteria, the *bolW* ancestor gene likely emerged from an ancient gene duplication event, followed by diverging evolution. In adapting to the secondary metabolism alongside with accessory enzymes, it warrants metabolic fluxes that allow the incorporation of tricarboxylic fatty acids into lipopeptides, thus increasing structural diversity.

In conclusion, by genome mining of infamous food-spoiling bacteria we discovered a new family of antibiotics that are an important addition to the metabolic inventory of diverse, ecology- and health-relevant *B. gladioli* strains. The bolagladins feature an unprecedented lipopeptide modification, a tricarboxylic fatty acid that creates a natural bolaamphiphile. We unveiled the key role of a citrate synthase in the *bol* pathway and provide an evolutionary model for the recruitment of a biocatalyst from the Krebs cycle. Using this designated citrate synthase as a bioinformatics handle led to the discovery of related cryptic pathways. Furthermore, the unusual lipopeptide assembly line, including the novel enamide-forming desaturase, may provide novel tools for bioengineering approaches.

## Acknowledgments

We thank H. Heinecke for NMR, A. Perner for MS/MS measurements, C. Weigel for bioactivity assays. This work was financially supported by the DFG (SFB 1127, ChemBioSys, and Leibniz Prize to C. H.).

## Note

The structure of bolagladin has been reported in the PhD thesis of a coauthor (Dr. Claudia Ross, Dissertation, Friedrich Schiller University Jena, 2015). This work and a related study by the Challis and Mahenthiralingam workgroups (https://www.biorxiv.org/content/10.1101/2020.06.16.153940v1.full.pdf+html) will be published back-to-back as companion papers.

## References

[1] M. Anwar, A. Kasper, A. R. Steck, J. G. Schier, J. Med. Toxicol. 2017, 13, 173–179.

[2] B.. Rohm, K. Scherlach, Hertweck C., Org. Biomol. Chem. 2010, 8, 1520–1522

b) N. Moebius, C. Ross, K. Scherlach, B. Rohm, M. Roth, C. Hertweck, Chem. Biol. 2012, 19, 1164–1174

c) C. Ross, V. Opel, K. Scherlach, C. Hertweck, Mycoses 2014, 57, 48–55.

[3] R. D. Süssmuth, A. Mainz, Angew. Chem. Int. Ed. 2017, 56, 3770–3821.

[4] N. Ziemert, M. Alanjary, T. Weber, Nat. Prod. Rep. 2016, 33, 988–1005.

[5] T. Thongkongkaew, W. Ding, E. Bratovanov, E. Oueis, M. A. Garcia-Altares, N. Zaburannyi, K. Harmrolfs, Y. Zhang, K. Scherlach, R. Müller, C. Hertweck, ACS Chem. Biol. 2018, 13, 1370–1379

b) B. Dose, S. P. Niehs, K. Scherlach, L. V. Flórez, M. Kaltenpoth, C. Hertweck, ACS Chem. Biol. 2018, 13, 2414–2420.

[6] R. Hermenau, K. Ishida, S. Gama, B. Hoffmann, M. Pfeifer-Leeg, W. Plass, J. F. Mohr, T. Wichard, H. P. Saluz, C. Hertweck, Nat. Chem. Biol. 2018, 14, 841–843

b) R. Hermenau, J. L. Mehl, K. Ishida, B. Dose, S. J. Pidot, T. P. Stinear, C. Hertweck, Angew. Chem. Int. Ed. 2019, 58, 13024–13029.

[7] M. J. Jaremko, T. D. Davis, J. C. Corpuz, M. D. Burkart, Nat. Prod. Rep. 2020, 37, 355–379.

[8] R. Finking, M. A. Marahiel, Annu. Rev. Microbiol. 2004, 58, 453–488

b) H. Jenke-Kodama, E. Dittmann, Nat. Prod. Rep. 2009, 26, 874–883.

[9] S. P. Niehs, B. Dose, K. Scherlach, S. J. Pidot, T. P. Stinear, C. Hertweck, ACS Chem. Biol. 2019, 14, 1811–1818.

[10] R. Bhushan, H. Bruckner, J. Chromatogr. B. Analyt. Technol. Biomed. Life. Sci. 2011, 879, 3148–3161.

[11] S. Koshino, H. Koshino, N. Matsuura, K. Kobinata, R. Onose, K. Isono, H. Osada, J. Antibiot. 1995, 48, 185–187

b) S. Son, S. K. Ko, S. M. Kim, E. Kim, G. S. Kim, B. Lee, I. J. Ryoo, W. G. Kim, J. S. Lee, Y. S. Hong, J. H. Jang, J. S. Ahn, J. Nat. Prod. 2018, 81, 2462–2469.

[12] M. Strieker, M. A. Marahiel, Med. Res. Rev. 2009, 10, 607–616

b) S. A. Cochrane, J. C. Vederas, Med. Res. Rev. 2016, 36, 4–31

c) N. Roongsawang, K. Washio, M. Morikawa, Int. J. Mol. Sci. 2011, 12, 141–172.

[13] I. Mnif, D. Ghribi, Biopolymers 2015, 104, 129–147.

[14] J. H. Fuhrhop, T. Wang, Chem. Rev. 2004, 104, 2901–2937

b) M. Fariya, A. Jain, V. Dhawan, S. Shah, M. S. Nagarsenker, Adv. Pharm. Bull. 2014, 4, 483–491.

[15] F. E. Cole, M. G. Kalyanpur, C. M. Stevens, Biochemistry 2002, 12, 3346–3350

b) S. L. Bulfer, E. M. Scott, J. F. Couture, L. Pillus, R. C. Trievel, J. Biol. Chem. 2009, 284, 35769–35780.

[16] B. Bonsch, V. Belt, C. Bartel, N. Duensing, M. Koziol, C. M. Lazarus, A. M. Bailey, T. J. Simpson, R. J. Cox, Chem. Commun. 2016, 52, 6777–6780.

[17] K. Williams, A. J. Szwalbe, N. P. Mulholland, J. L. Vincent, A. M. Bailey, C. L. Willis, T. J. Simpson, R. J. Cox, Angew. Chem. Int. Ed. 2016, 55, 6784–6788.

[18] B. Wang, Q. Kang, Y. Lu, L. Bai, C. Wang, Proc. Natl. Acad. Sci. U. S. A. 2012, 109, 1287–1292

b) T. Y. Lin, L. S. Borketey, G. Prasad, S. A. Waters, N. A. Schnarr, ACS Chem. Biol. 2013, 2, 635–642

c) F. Kudo, A. Miyanaga, T. Eguchi, Nat. Prod. Rep. 2014, 31, 1056–1073.

[19] R. Zallot, N. O. Oberg, J. A. Gerlt, Curr. Opin. Chem. Biol. 2018, 47, 77–85

b) J. A. Gerlt, Biochemistry 2017, 56, 4293–4308.

