## Supporting Material for "Food-Poisoning Bacteria Employ a Citrate Synthase and a Type II NRPS to Synthesize Bolaamphiphilic Lipopeptide Antibiotics"

### Table of contents

List of figures 3

List of tables 4

Experimental procedures 5

Bacterial strains and media 5

General analytical methods 6

Identification, isolation and purification of bolagladin 6

Absolute configuration of amino acids 7

Sequencing and annotation of bolagladin gene cluster 8

Phylogenetic analysis of C domains from the bolagladin gene cluster 12

Strains containing the bolagladin gene cluster 15

Gene cluster comparison using EasyFig2.3 16

Genetic manipulation of *Burkholderia gladioli* pv. *cocovenenans* HKI0521 17

Construction of the knockout plasmids 17

Gene knockout of *Burkholderia gladioli* pv. *cocovenenans* HKI0521 19

Bioactivity assays 21

Antibiotic assays 21

Swarming assays 22

References 24

NMR data and spectra 25

### List of figures


Figure S1: Bolagladin biosynthetic gene cluster (*bol*). 9
Figure S2. Phylogenetic analysis of C domains from BolA and from the NaPDos website. 12
Figure S3. Phylogenetic analysis of *Si*- and *Re*-Citrate synthase (CS) proteins. 13
Figure S4. Extract of the multiple sequence alignment of citrate synthase amino acid sequences confirms conservation of catalytically active residues in BolW (marked with red boxes). 14
Figure S5. Comparison of bolagladin gene clusters from *Burkholderia gladioli* (*Bg*.) spp. isolated from different sources. 16
Figure S6. Droplet collapsing assay to test for surface activity of wild type *versus* mutant strains. 22
Figure 7. *B. gladioli* pv. *cocovenenans* or bolagladin incubated on CAS agar plates at 30 °C, respectively, 20 °C for 24 h. 23
Figure S8. (-)ESI MS/MS spectrum of bolagladin A (1). 25
Figure S9. (+)ESI MS/MS spectrum of bolagladin A (1). 26
Figure S10. (+)ESI MS/MS spectrum of bolagladin A (1), mass range m/z 300-500. 26
Figure S11. (+)ESI MS/MS spectrum of the fragment *m/z* 450 amu derived from bolagladin A (1). 27
Figure S12. $^{1}$H NMR spectrum of bolagladin A (1). 30
Figure S13. $^{13}$C NMR spectrum of bolagladin A (1). 31
Figure S14. DEPT-135 NMR spectrum of bolagladin A (1). 32
Figure S15. H,H-COSY spectrum of bolagladin A (1). 33
Figure S16. HSQC spectrum of bolagladin A (1). 34
Figure S17. HMBC spectrum of bolagladin A (1). 35
Figure S18. (+)ESI MS/MS spectrum of bolagladin B (2). 36
Figure S19. $^{1}$H NMR spectrum of bolagladin B (2). 39
Figure S20. $^{13}$C NMR spectrum of bolagladin B (2). 40
Figure S21. H,H-COSY spectrum of bolagladin B (2). 41
Figure S22. HSQC spectrum of bolagladin B (2). 42
Figure S23. HMBC spectrum of bolagladin B (2). 43
Figure 24. Selected diagnostic $^{13}$C NMR signals of bolagladin A (1)and bolagladin M749 (3). 44
Figure S25. (-)ESI MS/MS spectrum of bolagladin M749 (3). 47
Figure S26. (+)ESI MS/MS spectrum of bolagladin M749 (3). 47
Figure S27. (+)ESI MS/MS spectrum of bolagladin M749 (3) mass range m/z 300-500. 48
Figure S28. (+)ESI MS/MS spectrum of the fragment *m/z* 362 amu derived from bolagladin M749 (3). 48
Figure S29. (+)ESI MS/MS spectrum of bolagladin M765 (4). 49
Figure S30. (+)ESI MS/MS spectrum of bolagladin M765 (4) mass range m/z 300-500. 50
Figure S31. (+)ESI MS/MS spectrum of the fragment *m/z* 360 amu derived from bolagladin M765 (4). 50
Figure 32. (+)ESI MS/MS spectrum of bolagladin M839 (5). 51
Figure 33. (+)ESI MS/MS spectrum of bolagladin M839 (5) mass range m/z 300-500. 52
Figure 34. (+)ESI MS/MS spectrum of bolagladin M824 (6). 53
Figure 35. (+)ESI MS/MS spectrum of bolagladin M824 (6) mass range m/z 300-500. 54

### List of tables

**Table S1. Bacterial and fungal strains used in this study. 5**
**Table S2. Composition of media used in this study. 5**
**Table S3. Gradient used for Marfey method. 7**
**Table S4. Analysis of the absolute configuration of amin acids bolagladin A (1) and B (2) using Marfey method. 7**
**Table S5. Analysis of the configuration of Isoleucine in bolagladin A (1) using Sanger reagent. 7**
**Table S6. Prediction of the NRPS modules of BolA according to antiSMASH 5.0; M – Module; C – condensation, A – adenylation, T – peptidylcarrier domain, TE – thioesterase, X – no prediction available, Ser – serine, Ile – isoleucine, Hse – homoserine, Val - valine. 8**
**Table S7. Proteins encoded up- and downstream of the NRPS *bolA* gene. 10**
**Table S8. Proteins encoded up- and downstream of the NRPS bolA (continued). 11**
**Table S9. Genomes of Burkholderia spp. that contain the bolagladin NRPS gene cluster. 15**
**Table S10. Detailed information on plasmids constructed using cloning strategy I. 17**
**Table S11. Detailed information plasmids constructed using cloning strategy II. 18**
**Table S12. Detailed information on the plasmids constructed using cloning strategy III; hr, homologous region; KanR, Kanamycin resistance cassette. 19**
**Table S13. Primer pairs and expected product sizes for colony PCRs to test for homologous recombination. 20**
**Table S14. Primer pairs and expected product sizes for colony PCRs to test for homologous recombination. (continued) 21**
**Table S15. Inhibitory effects of bolagladin (1) against selected strains. 21**
**Table S16. NMR data of bolagladin A (1). 28**
**Table S17. NMR data of bolagladin B (2). 37**
**Table 18. NMR data of bolagladin M749 (3). 45**

### Experimental procedures

#### Bacterial strains and media

Bacterial and fungal strains used in this study are listed in table S1. *Burkholderia gladioli* pv. *cocovenenans* HKI0521 was continuously cultured in MGY+M9, PDB or CYMG medium (Table S2). For long-term storage, the bacteria were frozen with 1:1 volume of 50% glycerol at –80 °C. *Burkholderia* gladioli pv. *cocovenenans* HKI0521 mutant strains were either cultured in MGY+M9 or PDB medium/agar with addition of kanamycin (50 µg $mL^{-1}$). *E. coli* strains were cultured in LB medium or on LB agar plates at 37 °C with appropriate antibiotic concentrations (kanamycin 50 µg $mL^{-1}$).

**Table S1.** Bacterial and fungal strains used in this study.

| Bacterial strain | Additional information | Reference |
|---|---|---|
| *B. gladioli* pv. *cocovenenans* | HKI0521 | - |
| *B. gladioli* DSM4285 | ATCC 10248[1] | |
| *Escherichia coli* | One Shot™ TOP10 Electrocomp™ *E. coli* | ThermoFisher Scientific |

**Table S2.** Composition of media used in this study.

| Medium or *medium additive* | Composition ($L^{-1}$) |
|---|---|
| MGY+M9 medium | 10 g Glycerol, 1.25 g yeast extract (technical yeast extract, BD, Bacto®), 960 mL distilled water, sterilization at 120 °C for 20 min, then add: 20 mL M9 salt A, 20 mL M9 salt B |
| CYMG | 10 g Glycerol, 8 g casein peptone, 4 g yeast extract (technical yeast extract, BD, Bacto®), 6.29 g $MgCl_2$*6 $H_2O$, sterilization at 120 °C for 20 min |
| PDB | Potato dextrose broth (BD, Bacto®) |
| PDA | PDB with addition of 1.5% agar (BD, Bacto®) |
| LB medium/agar | 10 g Tryptone (BD, Bacto®), 5 g yeast extract (BD, Bacto®), 10 g NaCl, sterilization at 120 °C for 20 min; for agar: addition of 1.5% agar |
| NAG agar | 34 g Standard I nutrient agar (Merk KGaA), 10 g glycerol, sterilization at 120 °C for 20 min |
| NA | Standard I nutrient agar (Merk KGaA) |
| *M9 salt solution A* | 350 g $K_2HPO_4$, 100 g $KH_2PO_4$, sterilization at 120 °C for 20 min |
| *M9 salt solution B* | 29.4 g Sodium citrate, 50 g $(NH_4)_2SO_4$, 5 g $MgSO_4$, sterilization at 120 °C for 20 min |

**General analytical methods**

LC-MS analysis was performed using an Exactive Orbitrap High Performance Benchtop LC-MS (Thermo Fisher Scientific) with an electron spray ion source and an Accela HPLC System, C18 column (Betasil C18, 150 x 2.1 mm, Thermo Fisher Scientific, Germany), solvents: acetonitrile and water (both supplemented with 0.1% formic acid), flow rate: 0.2 mL $min^{-1}$; program: hold 1 min at 5% acetonitrile, 1–16 min 5–99% acetonitrile, hold 16–31 min 99% acetonitrile, 31-32 min 99–5%, 32–43 min to 5% acetonitrile. For MS-MS measurements a QExactive Orbitrap High Performance Benchtop LC-MS (Thermo Fisher Scientific) with an electron spray ion source and an Accela HPLC System was used (column: Accucore C18 2.6 µm, 100 x 2.1 mm, Thermo Fisher Scientific solvents: acetonitrile and water (both supplemented with 0.1% formic acid), flow rate: 0.2 mL $min^{-1}$; program: hold 1 min at 5% acetonitrile, 1–16 min 5–99% acetonitrile, hold 16-31 min 99% acetonitrile, 31–32 min 99–5 %, 32–43 min to 5% acetonitrile. NMR spectra were recorded with a Bruker 500 or 600 MHz Avance III Ultra Shield (Bruker BioSpin GmbH) in DMSO-$d_6$ $^1H$ 600 MHz; $^{13}C$ 150 MHz.

**Identification, isolation and purification of bolagladin**

For isolation of bolagladins, bacteria were cultured in PDB media. 50 mL of PDB medium was inoculated ($OD_{600}$ ~ 0.4) and incubated at 30 °C overnight, at 120 rpm. Fourteen 1 L flasks with 250 mL of PDB each were inoculated with 4 mL of this bacterial preculture and incubated without shaking at 30 °C for 4 d. The cultures were thoroughly extracted twice with 1:1 volume of ethyl acetate. The organic extract was dried with sodium sulfate and concentrated under reduced pressure. The residue was dissolved in methanol and separated by size exclusion chromatography using Sephadex LH-20 (Pharmacia Fine Chemicals) with MeOH as eluent. Bolagladin-containing fractions were concentrated (minimum: 5 mL methanol). Final purification was achieved by preparative reversed-phase HPLC (Shimadzu LC-8A HPLC system) using either a Phenomenex Synergi 4 µm Fusion-RP 80 Å, 250 × 21.2 mm column with gradient MeCN/$H_2O$ supplemented with 0.01% trifluoroacetic acid (*v/v*) (58/42 in 20 min to 80/20, 80/20 in 2 min to 100/0. MeCN 100% for 10 min, flow rate 12 mL $min^{-1}$).

### Absolute configuration of amino acids

Bolagladins were hydrolyzed with 6 M HCl for 12 h at 105 °C and subsequently dried under reduced pressure. After the addition of 100 µL 1 M $NaHCO_3$ and 50 µL 1-fluoro-2,4-dinitrophenyl-5-L-alaninamide solution (L-FDAA, Marfey reagent, 10 mg $mL^{-1}$ acetone) the samples were heated for 1 h at 50 °C. 50 µl 2 M HCl was added as well as 200 µL 50% (*v/v*) MeCN. The samples were analyzed with an analytical HPLC and a Kinetex 5 µm XB-C18 100 Å 250 × 4.6 µm (Phenomenex) column and a flow rate of 0.5 mL $min^{-1}$. MeCN and $H_2O$ both supplemented with 0.1% trifluoroacetic acid were used. The gradient can be found in Table S3.

**Table S3**. Gradient used for Marfey method.

| Zeit (min) | $dH_2O$ (%) | ACN (%) |
|---|---|---|
| 5 | 70 | 30 |
| 30 | 50 | 50 |
| 35 | 0 | 100 |
| 37 | 70 | 30 |
| 42 | 70 | 30 |

The retention times of derivatized amino acids can be found in Table S4 and Table S5. To distinguish isoleucine and *allo*-isoleucine the samples were treated with L-FDAA 50 µl 1-fluoro-2,4-dinitrobenzol (Sanger reagent) (10 mg $mL^{-1}$ acetone). A Eurospher C18 5 µm 100 Å 250 × 4.6 µm column and an isocratic flow of 40% MeCN in $H_2O$ supplemented with 0.1% trifluoroacetic acid was used.

**Table S4**. Analysis of the absolute configuration of amin acids bolagladin A (**1**) and B (**2**) using Marfey method.

| Amino acid | Configuration | Retention time of derivatized standard amino acid [min] | Retention time of derivatized amino acid in bolagladin A (**1**) [min] | Retention time of derivatized amino acid in bolagladin B (**2**) [min] |
|---|---|---|---|---|
| Serine | D | 11.6 | 11.4 (L) | 11.4 (L) |
| | L | 11.4 | | |
| Valine | D | 30.4 | - | 30.4 (D) |
| | L | 25.3 | | |
| Homoserine | D L | 12.0 and 12.4 | 12.0 (L) | 12.0 (L) |
| | L | 12.0 | | |
| Isoleucine | D | 35.1 | 35.1 (D) | - |
| | L | 29.8 | | |

**Table S5.** Analysis of the configuration of Isoleucine in bolagladin A (1) using Sanger reagent.

| Amino acid | Configuration | Retention time of standard amino acid [min] | Retention time of amino acid in bolagladin A (**1**)[min] |
|---|---|---|---|
| Isoleucine | D | 25.5 | 25.5 (D) |
| | D-*allo* | 26.5 | |

Comparison of the retention times of the standard amino acid and the samples allowed for identification of the amino acid building blocks of bolagladin A (L-serine, D-*allo*-isoleucine, L-homoserine and L-serine) and bolagladin B (L-serine, D-valine, L-homoserine and L-serine). This is in accordance with the predictions based on the Stachelhaus code.

### Sequencing and annotation of bolagladin gene cluster

The bolagladin gene cluster (*bol*) was uploaded to the NCBI database, accession number: (xxx). The webtool antiSMASH 5.0[2] was used to for the identification, annotation and analysis of secondary metabolite biosynthesis gene clusters in the genome of *Burkholderia gladioli* pv. *cocovenenans* HKI0521. The bolagladin gene cluster consists of a 13.6 kb NRPS gene that contains four modules (Table S6). The predicted amino acid and the Stachelhaus code were taken from the antiSMASH analysis.

**Table S6.** Prediction of the NRPS modules of BolA according to antiSMASH 5.0; M – Module; C – condensation, A – adenylation, T – peptidylcarrier domain, TE – thioesterase, X – no prediction available, Ser – serine, Ile – isoleucine, Hse – homoserine, Val - valine.

| **Modules of BolA** | **Specificity code (antiSMASH)** | **Predicted amino acid (Stachelhaus code match)** | **Amino acid sequence of bolagladin A** |
| --- | --- | --- | --- |
| M1 (C-A-T) | DVWHMSLVDK | Ser (100%) | Ser |
| M2 (C-A-T) | DALWMGGVFK | Val (80%) | Ile |
| M3 (C-A-T) | DLKNVGSDVK | X | Hse |
| M4 (C-A-TE) | DVWHVSLIDK | Ser (100%) | Ser |

NCBI Blast and HHpred[3] were used for sequence database searching to find homologous genes or proteins that allow for speculations on their functionalities. A promotor search was conducted using PromotorHunter (http://www.phisite.org/promoterhunter). Based on the calculated Gibbs energy and a position-specific weight matrix, a scoring system was used to predict putative promotor sites.

| Matrix (–35 region): | Matrix (–10 region): |
|---|---|
| A: 10 6 9 56 21 54 | A: 5 76 15 61 56 6 |
| C: 10 7 12 17 54 13 | C: 10 6 11 13 20 7 |
| G: 10 8 61 11 9 16 | G: 8 6 14 14 8 5 |
| T: 70 79 18 16 16 1782 | T: 77 12 60 12 15 |

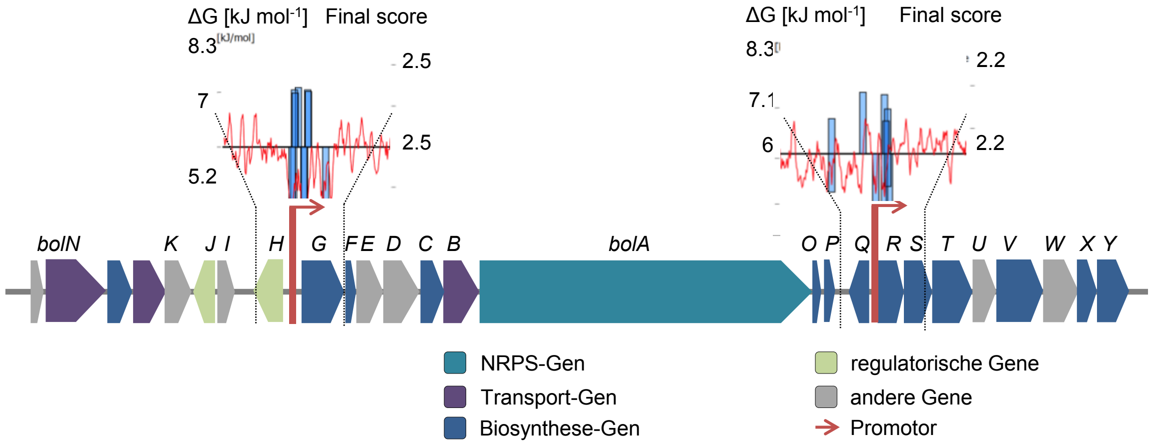


**Figure S1**: Bolagladin biosynthetic gene cluster (*bol*).

Further information on the genes respectively encoded proteins can be found in Table S7.

**Table S7.** Proteins encoded up- and downstream of the NRPS *bolA* gene.

| Gene | Length [AA] | Putatively encoded protein | Closest characterized orthologous protein and (organism) | Accession number | Identity/ similarity |
|---|---|---|---|---|---|
| *bolO* (-1) | 69 | Domain binding protein | MbtH-Homolog (*Geobacillus* sp. Y4.1MC1) | 5U89_B | 40%/90% |
| *bol*P (–2) | 140 | / | / | / | / |
| *bolQ* (–4) | 281 | Reductase/ desaturase | Possible acyl desaturase (*Mycobacterium tuberculosis*) | 1ZA0_A | 19%/20% |
| *bolR* (–5) | 364 | Transferase | Alkyldiketide–CoA synthase; polyketidesynthase (*Tetradium ruticarpum*) | 5WX6_A | |
| *bolS* (–6) | 392 | Decarboxylase | Diaminopimelate decarboxylase (*Actinosynnema pretiosum* subsp. *auranticum*) | O69203.1 | 28%/36% |
| *bolT* (–7) | 527 | Long-chain-fatty-acid-CoA ligase | Long chain fatty acid-CoA ligase (*Bacillus subtilis*) | O07610.2 | 29%/44% |
| *bolU* (–8) | 260 | Fatty acid biosynthesis | 3-Oxoacyl-acyl-carrier-protein synthase III (*Haemophilus influenzae*) | 3IL3_A | 17%/11% |
| *bolV* (–9) | 615 | Acyl-CoA dehydrogenase | Dehydrogenation (*Roseovarius nubinhibens* ISM) | 6IJC_B | 29%/48% |
| *bolW* (–10) | 459 | Citrate synthase | Citrate synthase (*Burkholderia thailandensis*) | 4XGH_A | 20%/18% |
| *bolX* (–11) | 220 | Methyltransferase | SAM–dependent methyltransferase (*Thiobacillus denitrificans*) | 5EPE_A | 16%/16% |
| *bolY* (–12) | 357 | Dioxygenase/ Decarboxylase | Clavaminate synthase- Protein ScoE Protein (*Arabidopsis thaliana*) | 2Q4A_A/ | 30%/54% |

**Table S8.** Proteins encoded up- and downstream of the NRPS gene *bolA* (continued).

| Gene | Length [AA] | Putatively encoded protein | Closest characterized orthologous protein and (Organism) | Accession number | Identity/ similarity |
|---|---|---|---|---|---|
| *bol*A (0) | 4,547 | NRPS | Tyrocidine synthase 3 (*Brevibacillus parabrevis*) | O30409.1 | 35%/51% |
| *bolB* (+1) | 441 | Major facilitator superfamily (MFS) | MFS-type transporter MT0042 (*Mycobacterium tuberculosis*) | P9WJY0.1 | 28%/28% |
| *bolC* (+2) | 343 | Desaturase | Sphingolipid Δ(4)-desaturase/C4-monooxygenase DES2 (*Homo sapiens*) | Q6QHC5.2 | 28%/41% |
| *bolD* (+3) | 480 | Dehydrogenase, putative oxidoreductase | D-2-hydroxyglutarate dehydrogenase (*Oryza sativa*) | Q7XI14.1 | 33%/50% |
| *bolE* (+4) | 352 | Oxygenase/ reductase | p-Aminobenzoate N-oxygenase (*Streptomyces thioluteus*) | 3CHH_B | 15%/4% |
| *bolF* (+5) | 82 | ACP | Acyl carrier protein (*Nitrosomonas europaea*) | 2LKI_A | 15%/18% |
| *bolG* (+6) | 583 | Ligase/A-domain | Long-chain-fatty-acid ligase (*Mycobacterium tuberculosis*) | P0A4X9.1/ | 37%/48% |
| *bolH* (+7) | 54 | / | / | / | / |
| *bolI* (+8) | 339 | HTH-type transcriptional regulator | Transcriptional activator (*Escherichia coli* O157) | P0A9F8.1 | 33%/52% |
| *bolJ* (+9) | 140 | Putative plasmid stability protein | VapC ribonuclease (*Sinorhizobium fredii*) | P55511.1 | 32%/48% |
| *bolK* (+10) | 226 | / | Deoxyribose-phosphate aldolase (*Burkholderia lata*) | Q39NL8.1 | 85%/90% |
| *bolL* (+11) | 296 | Transcriptional regulator | Deoxyribose operon repressor (*Escherichia coli* O157) | P0ACK7.1 | 43%/57% |
| *bolM* (+12) | 333 | Ribokinase | Ribokinase (*Homo sapiens*) | Q9H477.1 | 38%/54% |
| *bolN* (+13) | 804 | TonB-dependent Rezeptor | Enterobactin-iron-transporter (*Pseudomonas aeruginosa*) | Q05098.1 | 22%/37% |

#### Phylogenetic analysis of C domains from the bolagladin gene cluster

C domain amino acid sequences of the bolagladin biosynthesis gene cluster as well as sequences acquired from the NaPDos website[4] were aligned using MAFFT[5] with default settings. Maximum likelihood phylogeny was constructed using IQ-tree[6]. Ultrafast bootstrapping (1000 iterations) analysis was performed. The neighbor-joining tree construction method showed similar tree topology. The phylogeny indicates the functionalities of the subjected C domains and allows a proposal for the biosynthetic logic of bolagladin.

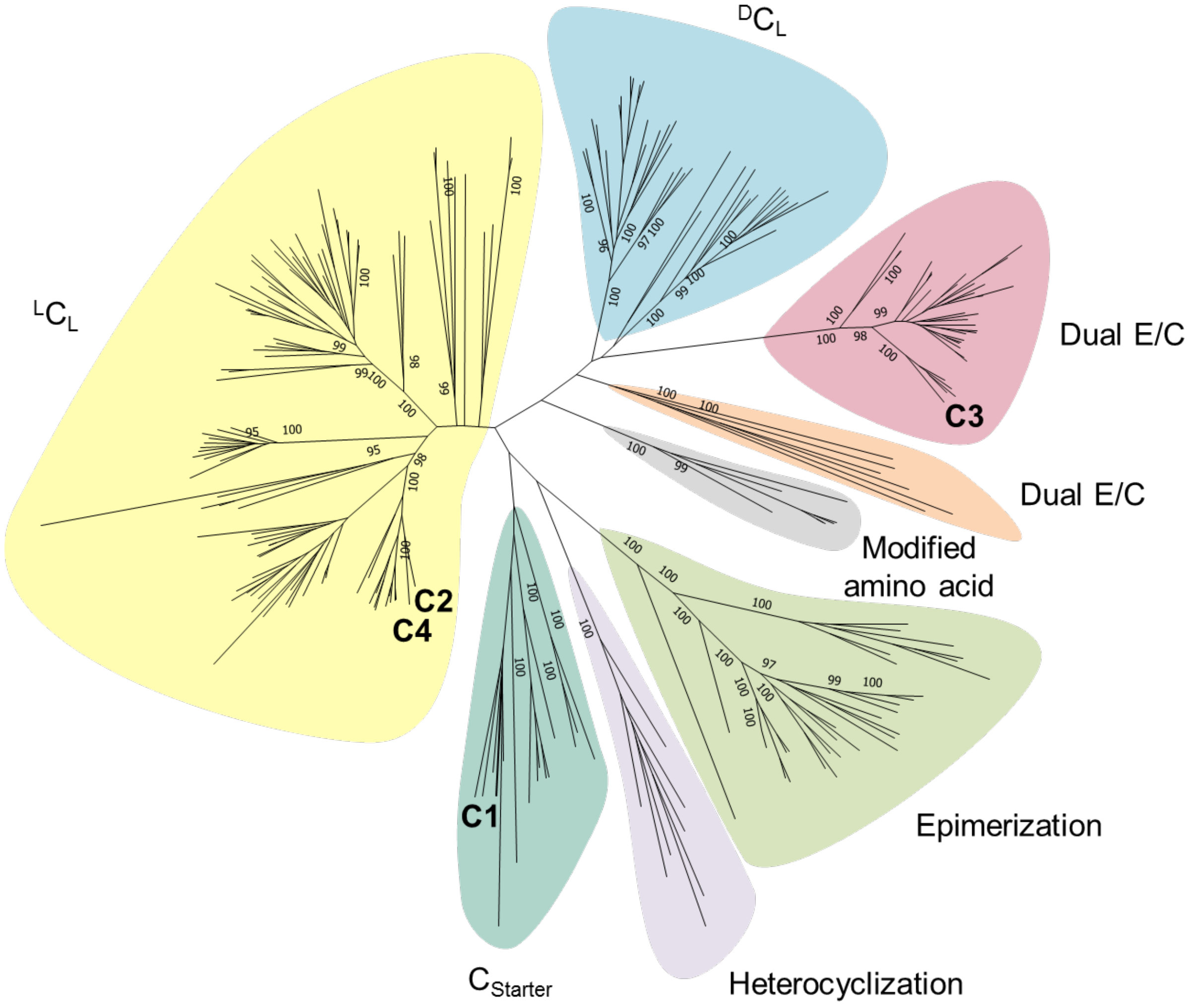


**Figure S2**. Phylogenetic analysis of C domains from BolA and from the NaPDos website.
C domains cluster into the canonical subfamilies ($^{L}C_L$, $^{D}C_L$, starter, dual E/C, epimerization, heterocyclization domains and modified amino acid domain).

#### Phylogenetic analysis of the citrate synthase BolW

**Characterized (bold) and non-characterized *Re*- and *Si*-citrate synthases amino acid sequences were downloaded from Uniprot database (Figure S3Figure S2**. Phylogenetic analysis of C domains from BolA and from the NaPDos website.

C domains cluster into the canonical subfamilies ($^{L}C_L$, $^{D}C_L$, starter, dual E/C, epimerization, heterocyclization domains and modified amino acid domain).Figure S2). Additionally, three proteins annotated as citrate synthases encoded in the genome of *B. gladioli* pv. *cocovenenans* were added, one of which was encoded in the bolagladin biosynthesis gene cluster. Citrate synthase proteins from the phomoidride[7], maleidrides[8] and squalestatin[9] biosynthesis pathways were added. Malate synthesis amino acid sequences were added as outgroup. The sequence alignment was performed using MAFFT 7 with L-INS-I, an iterative refinement method. Maximum likelihood phylogeny was constructed using IQ-tree. Ultrafast bootstrapping (1000 iterations) analysis was performed. The bolagladin citrate synthase (BolW), as well as the CS-phomoidride, CS-maleidrides and CS-squalestatin clusters with *Si*-Citrate synthases. *Re*-CS cluster next to 2-isopropylmalate (EC 2.3.3.13), citramalate synthases (EC 2.3.3.14) and are sometimes wrongly annotated such because they have a high sequence similarity. Notable, citramalate synthase or 2-isopropylmalate synthases are *Re*-face stereospecific with respect to C-2 of their substrates, 2-oxoglutarate and 2-oxo-3-methylbutyrate.[10] Inspection of the multiple sequence alignment of BolW shows conservation of reported, catalytically active residues His302, His350 and Asp356 (Figure S4).

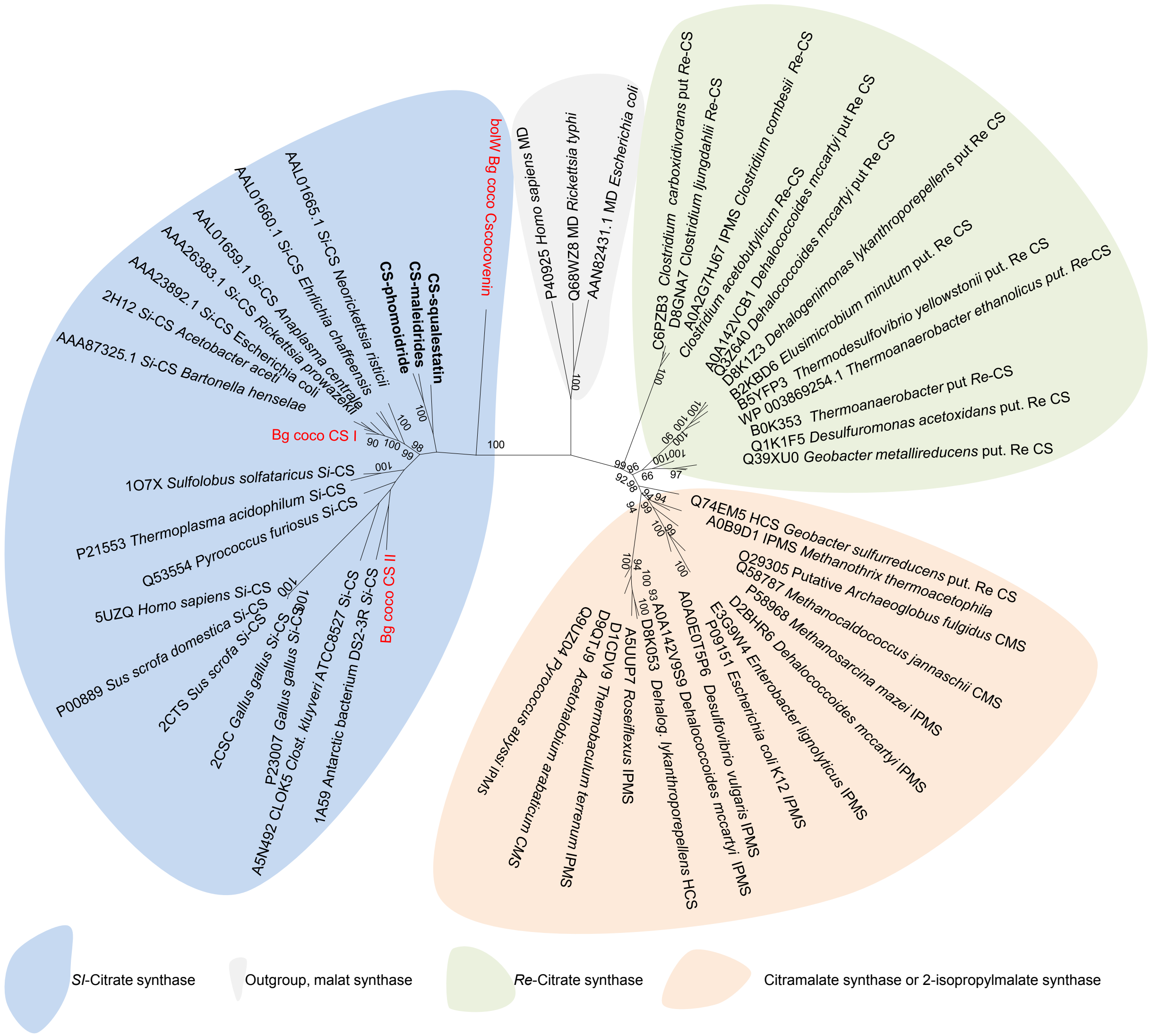


**Figure S3**. Phylogenetic analysis of *Si*- and *Re*-Citrate synthase (CS) proteins.
Citramalate synthase (CMS), homocitrate synthase (HCS) and 2-siopropylmalate synthase (IPMS) proteins are included. Malate syntases are used as outgroup. Alignment and construction of the phylogenetic tree was performed using MAFFT 7 respectively IQ-Tree. Malate synthases were chosen as an out group. Ultrafast bootstrapping (1000 iterations) analysis was performed. Red characters, B. gladioli pv. cocovenenans CS; green characters CS from other secondary metabolite pathways, bold characters in vitro characterized proteins.

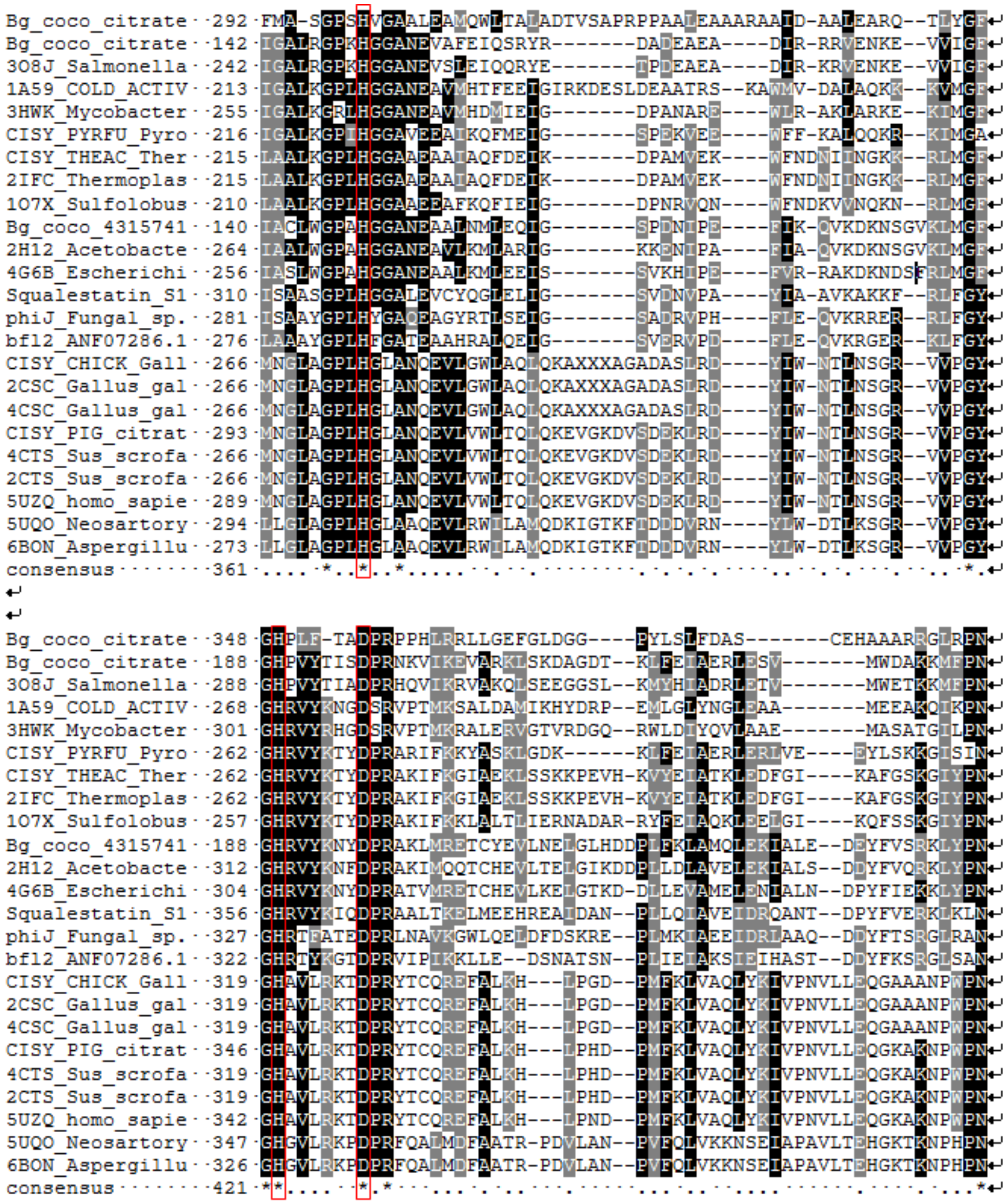


**Figure S4.** Extract of the multiple sequence alignment of citrate synthase amino acid sequences confirms conservation of catalytically active residues in BolW (marked with red boxes).

#### Strains containing the bolagladin gene cluster

**Table S9**. Genomes of Burkholderia spp. that contain the bolagladin NRPS gene cluster.

| Organism | Origin | Accession number |
|---|---|---|
| *B. gladioli* pv. *cocovenenans* HKI0521 | Contaminant in fungal culture in the fermentation of coconut patties | DSM11318 |
| *Burkholderia gladioli* AU26456 | Sputum, *Homo sapiens* | NZ_PVGZ01000021.1 |
| *Burkholderia gladioli* AU30473 | Sputum, *Homo sapiens* | NZ_PVHI01000143.1 |
| *Burkholderia gladioli* AU29541 | Sputum, *Homo sapiens* | NZ_PVHE01000070.1 |
| *Burkholderia gladioli 3723STDY6437372* | *Homo sapiens*, cystic fibrosis, USA | NZ_UWYX01000001.1 |
| *Burkholderia gladioli* BSR3 | Diseased rice sheath (South Korea) | NC_015376.1 |
| *Burkholderia gladioli* 3723STDY6437373 | Clinical; Pathogen (United Kingdom) | NZ_UWYW01000003.1 |
| *Burkholderia gladioli* Co14 | Fermented corn meal (Heilongjiang Province) | NZ_CP033431.1 |
| *Burkholderia gladioli* AU0032 | Sputum, *Homo sapiens* | NZ_PVFC01000046.1 |
| *Burkholderia gladioli* MSMB1756 | Soil | NZ_LOZK01000085.1 |
| *Burkholderia* BCC238 | Clinical | NZ_UZVS01000001.1 |
| *Burkholderia gladioli* pv. *gladioli* FDAARGOS_188 | - | NZ_CP022210.1 |
| *Burkholderia gladioli* NCTC12378 | Plant (*Gladiolus* species) | NZ_UARB01000021.1 |
| *Burkholderia gladioli* pv. *gladioli* KACC 11889 | *Gladiolus*, (South Korea) | NZ_CP022006.1 |
| *Burkholderia gladioli* ATCC 10248 | Plant | NZ_CP009322.1 |
| *Burkholderia gladioli* NBRC 13700 | Free living | NZ_BBJG01000123.1 |
| Burkholderia gladioli ATCC 25417 | Leaves, *Gladiolus* | NZ_KN150850.1 |

**Gene cluster comparison using EasyFig2.3**

The sequence similarity of *bol* gene clusters is visualized using the tool Easyfig 2.3.[11] Color code represents the sequence similarity values as indicated (**Figure S5**).

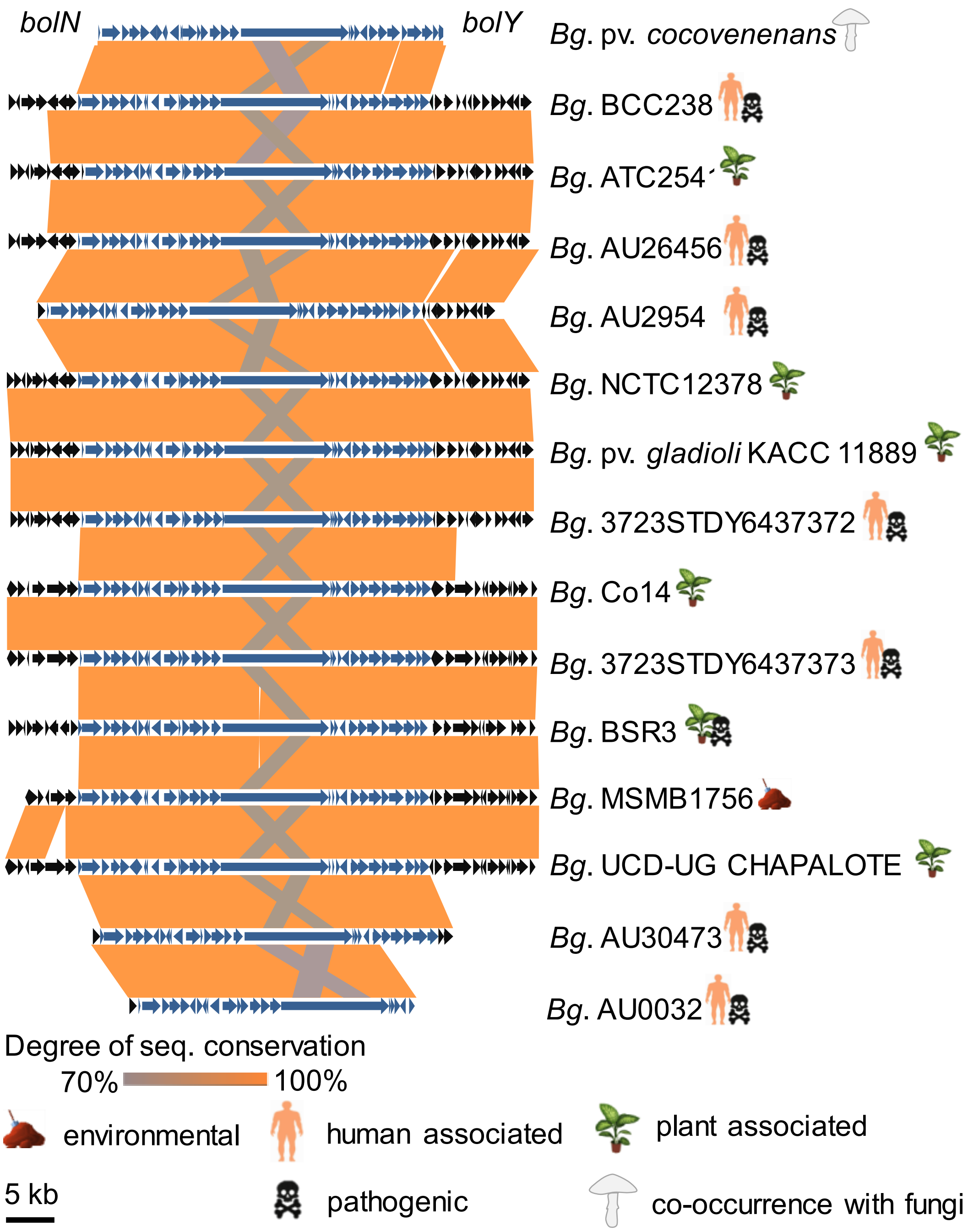


**Figure S5**. Comparison of *bol* gene clusters from *Burkholderia gladioli* (*Bg.*) spp. isolated from different sources.
Genes shared by all listed *bol* gene clusters are represented by blue arrows, others are depicted in black.

**Genetic manipulation of *Burkholderia gladioli* pv. *cocovenenans* HKI0521**

*Burkholderia gladioli* was genetically manipulated by homologous recombination to inactivate genes putatively involved in the biosynthesis of bolagladin. Homologous regions encoded on a suicide plasmid were recombined with the genome of the bacteria leading to the integration of a kanamycin resistance cassette and thereby rendering the target gene inactive.

**Construction of the knockout plasmids**

All experiments have been performed according to manufacturer's recommendations if not stated otherwise. Genomic DNA isolation of bacteria was performed using PureTM DNA-Isolation kit (Epicentre Biotechnologies). PCRs have been performed as follows: *Burkholderia gladioli* HKI0521genomic DNA, 2x master mix of KAPA2G Robust HotStart ReadyMix PCR kit (Merck KGaA) and 35 cycles of 95 °C for 30 s, 60 °C for 30 s $kb^{-1}$, and a final extension time at 72 °C for 30 s $kb^{-1}$. The following cloning strategies have been used to construct the knockout plasmid:

I: Amplification of homologous regions and subsequently restriction to introduce a kanamycin resistance cassette, construction of pJB02, pJB04, pJB06and pJB14. Primers listed in table S10 have been used in a PCR to amplify homologous regions from *Burkholderia gladioli* genomic DNA. The PCR product was blunt-end ligated into pJET1.2 linear vector (CloneJET PCR Cloning Kit, Thermo Fisher Scientific). The resulting plasmids as well as the kanamycin resistance cassettes amplified from pGEM-Kan[12] with the listed primers have been digested with the indicated restriction enzymes (Table S10).

**Table S10**. Detailed information on plasmids constructed using cloning strategy I.

| Plasmid (target gene) | Name and sequence 5' → 3' | Product size and name | Digestion enzyme |
|---|---|---|---|
| pJB02 (NRPS gene, *bolA*) | JB01, ccagttcgtcctcgctgtag<br>JB02, cctctacctgctggccttc | 2,299 bp, homologous region | *Eco*RI |
| | JB03, cgtggaattcgtaagcttaggctgctgcc<br>JB04, cgtggaattctcagaagaactcgtcaag | 1,182 bp, KanR | - |
| pJB04 (A domain gene, *bolG)* | JB07, cttctcgtcgagcttgttcc<br>JB08, gcctcgactacgaggtgttc | 2,371 bp, homologous region | *Sac*I |
| | BD247, cggacgagctcgtaagcttaggctg<br>BD248, cgcgcgagctctcagaagaactcgtcaag | 1,162 bp, KanR | - |
| pJB06 (desaturase gene, *bolC*) | JB11, cgcacatcgacatgaagc<br>JB12, tctcaagcagctgttcatcg | 2,071 bp homologous region | *Bam*HI |
| | BD357, ctcgggatccgtaagcttaggctgctgcc<br>BD358, tcaaggatcctcagaagaactcgtcaag | 1,162 bp KanR | - |
| pJB14 (fatty acid ligase gene, *bolT*) | BD406, gaacttgtggcggtgcag<br>BD407, gcgatgttcgacggctatc | 2,032 bp homologous region | *Sph*I |
| | BD408, gccggcatgctcagaagaactcgtcaag<br>BD409, aagcgcatgctcagccaatcggatgaatg | 1,088 bp, KanR | - |

II: NEBuilder mediated fusion of two homologous regions with a resistance cassette and subsequent pJet1.2 cloning, construction of pJB10, pJB12, and pBD87.

Primers listed in table S11 were used to amplify homologous regions from genomic DNA of *Burkholderia gladioli* pv. *cocovenenans* respectively the kanamycin resistance cassette from pGEM-Kan. In a subsequent step a NEBuilder three-fragments ligation have been performed (New England Biolabs, Frankfurt am Main) according to the manufacture's recommendations. Finally, the ligated DNA fragments have been blunt-end ligated into pJET1.2 linear vector yielding the final knockout plasmids.

**Table S11.** Detailed information plasmids constructed using cloning strategy II.

| Plasmid (target gene) | Name and sequence of primers used 5' → 3' | Product size and name |
|---|---|---|
| pJB10 (Thiolation domain gene, *bolF*) | JB27, ctcctttggattcctgatcg<br>JB28, ctaagcttacagatcaacctcacgcagg | 1,241 bp, homologous region |
| | JB29, aggttgatctgtaagcttaggctgctgc<br>JB30, tcgtgttctgtcagaagaactcgtcaagaag | 1,182 bp, KanR |
| | JB31, gttcttctgacagaacacgatcagctcg<br>JB32, acgtgctcgacaacagctac | 1,471 bp, homologous region |
| pJB12 (Desaturase gene, *bolQ*) | JB43, gtcgaagccggtctcgtag<br>JB44, gatggacgagctcatgaaggcggagaagtc | 1,296 bp, homologous region |
| | JB45, ccttcatgagctcgtccatcgtaagcttag<br>JB46, atagcgattcccgatcgatgtcagaagaac | 1,182 bp, KanR |
| | JB47, catcgatcgggaatcgctatcgcaaggaag<br>JB48, aggacatccacctcgacaag | 1,079 bp, homologous region |
| pBD87 (Ketosynthase type III gene, *bolR*) | BD424, gcgatagccgtcgaacatc<br>BD425, ctaagcttacctacgagaccggcttcgac | 1,206 bp, homologous region |
| | BD426, ggtctcgtaggtaagcttaggctgctgc<br>BD427, ccagccgttctcagaagaactcgtcaagaag | 1,162 bp, KanR |
| | BD428, gttcttctgagaacggctgggtatgcac<br>BD429, gacttctccgccttcatgag | 946 bp, homologous region |

III: Four fragment NEBuilder fusion of two homologous regions, a resistance cassette and a vector backbone. Primers listed in table ## were used to amplify two homologous regions from genomic DNA of *Burkholderia gladioli* pv. *cocovenenans* respectively the kanamycin resistance cassette from pJK354. The plasmid pJK354 was digested with *Xho*I and *Xba*I. In a subsequent step a NEBuilder four-fragment ligation has been performed according to the manufacture's recommendations yielding the final knockout plasmids (Table S12).

**Table S12.** Detailed information on the plasmids constructed using cloning strategy III; hr, homologous region; KanR, Kanamycin resistance cassette.

| Plasmid (target gene) | Name and sequence of primers used 5' → 3' | Product size and name |
|---|---|---|
| pBD92 (Decarboxylase gene, *bolS*) | BD459, tatagggagagcggccgccagatcttccggatggcagccggaggtgtagagcag<br>BD460, ctgcggactggctttctacgtgttcaatcactagtgatggacgaggacttcatgc | 1,498 bp, hr |
| | JK582, gaattcgattctggtcggtc<br>JK583, actagtgattgaacacgtag | 1,151 bp, KanR |
| | BD461, tctggggttcgaaatgaccgaccagaatcgaattccgggtcgtacacgtagcag<br>BD462, tttccatggcagctgagaatattgtaggagatcttgcggattcgacaagatgattg | 1,371 bp, hr |
| pBD93 (Ketosynthase type III gene, *bolU*) | BD476, tatagggagagcggccgccagatcttccggatggctcggaagtgccgaaatac<br>BD477, ctgcggactggctttctacgtgttcaatcactagtaagggatgcagccatttatc | 1,272 bp, hr |
| | JK582, gaattcgattctggtcggtc<br>JK583, actagtgattgaacacgtag | 1,151 bp, KanR |
| | BD478, tctggggttcgaaatgaccgaccagaatcgaattccgacacgctcgactggtg<br>BD479, tttccatggcagctgagaatattgtaggagatcttggctcttcatgtcgatcagc | 1,421 bp, hr |
| pBD95 (Citrat synthase gene, *bolW*) | BD459, tatagggagagcggccgccagatcttccggatggctctgggaaggcaccaactac<br>BD460, ctgcggactggctttctacgtgttcaatcactagtcacggtcggcgtgatatag | 1,387 bp, hr |
| | JK582, gaattcgattctggtcggtc<br>JK583, actagtgattgaacacgtag | 1,151 bp, KanR |
| | BD461, tctggggttcgaaatgaccgaccagaatcgaattccctacctgagcctgttcg<br>BD462, tttccatggcagctgagaatattgtaggagatcttatgtaggccatctcctgg | 1,434 bp, hr |
| pBD107 (acyl-CoA-dehydrogenase) | BD555, tatagggagagcggccgccagatcttccggatggccgcagggctacctctacttc<br>BD556, ctgcggactggctttctacgtgttcaatcactagtgacatccgattgctggtag | 1, 505 bp, hr |
| | JK582, gaattcgattctggtcggtc<br>JK583, actagtgattgaacacgtag | 1,151 bp, KanR |
| | BD557, tctggggttcgaaatgaccgaccagaatcgaattctcctgacccaggtgctctac<br>BD558, tttccatggcagctgagaatattgtaggagatcttcgaggaagatctccagttcg | 1,201 bp, hr |

### Gene knockout of *Burkholderia gladioli* pv. *cocovenenans* HKI0521

*Burkholderia gladioli* pv. *cocovenenans* HKI0521 cells were washed three times with one volume of 300 mM sucrose in $dH_2O$. Cells have been spun down at 6,000 x g for 5 min. Afterwards the cells have been suspended in 100 µL aliquots. 2 µL of a plasmid have been added to the cell solution and afterwards a pulse of 2.5 kV has been applied. Cells have been suspended in MGY media and recovered for 3–4 h at 30 °C and 120 rpm. Afterwards, cell solution was plated onto NAG agar containing 50 µg $mL^{-1}$ kanamycin and incubated at 30 °C until colonies appeared. Colonies grown after transformation with knockout plasmids have been subjected to a colony PCR to detect mutations. The templates were colony material that have been picked with a pipette tip and transferred into 100 µL $dH_2O$, incubated for 5 min at 99 °C and centrifuged at 18,000 × g for 1 min. 5 µL of the template have been added to 5 µL of 2x master mix of KAPA2G Robust HotStart ReadyMix PCR kit (Merck KGaA) with the appropriate primers.

Three separate PCRs have been performed to confirm homologous recombination in each mutant that has been created. The PCR were named front arm (FA), back arm (BA) and wild type control (WT). The primer pairs and expected products are listed (Table S13 and Table S14). The PCRs FA and BA only yield a product if the recombination of the front and respectively back homologous region with the genome took place. The WT PCR was used to exclude that wild type cells were present in the tested colony. As an additional control all PCRs were run in parallel with wild type cell material as template.

**Table S13**. Primer pairs and expected product sizes for colony PCRs to test for homologous recombination.

| Mutant strain | PCR | Name and sequence of used primers 5' → 3' | Product size |
|---|---|---|---|
| Bg_coco_pJB02 (*ΔbolA*) | FA | JB05, tatcgttccattcgctcaag<br>TIISSD_B_fwd, cgttggctacccgtgatatt | 1,360 bp |
| | BA | JB06, ctacaacctgccgaagatgc<br>TIISS_A_rv, agtgacaacgtcgagcacag | 2,224 bp |
| | WT | JB05, tatcgttccattcgctcaag<br>JB06, ctacaacctgccgaagatgc | 2,919 bp |
| Bg_coco_pJB04 (*ΔbolG*) | FA | JB09, gcgttaccaggtccttgatg<br>TIISS_A_rv, agtgacaacgtcgagcacag | 2,134 bp |
| | BA | JB10, tgtcacgtcatgggagattc<br>TIISSD_B_fwd, cgttggctacccgtgatatt | 1,505 bp |
| | WT | JB09, gcgttaccaggtccttgatg<br>JB10, tgtcacgtcatgggagattc | 2,880 bp |
| Bg_coco_pJB06 (*ΔbolC*) | FA | JB13, gaaggccagcaggtagagg<br>TIISSD_B_fwd, cgttggctacccgtgatatt | 1,038 bp |
| | BA | JB14, ctgctcaagtacggcgacac<br>TIISS_A_rv, agtgacaacgtcgagcacag | 2,180 bp |
| | WT | JB13, gaaggccagcaggtagagg<br>JB14, ctgctcaagtacggcgacac | 2,459 bp |
| Bg_coco_pJB10 (*ΔbolF*) | FA | JB33, gtgtcgccgtacttgagcag<br>TIISS_A_rv, agtgacaacgtcgagcacag | 2,363 bp |
| | BA | JB34, gcctcgactacgaggtgttc<br>TIISSD_B_fwd, cgttggctacccgtgatatt | 2,045 bp |
| | WT | JB33, gtgtcgccgtacttgagcag<br>JB34, gcctcgactacgaggtgttc | 4,818 bp |
| Bg_coco_BD87 (*ΔbolR*) | FA | BD430, gagtaggtgcggtgataggc<br>TIISS_A_rv, agtgacaacgtcgagcacag | 2,198 bp |
| | BA | BD431, tctgctcctcgaagaaccac<br>TIISSD_B_fwd, cgttggctacccgtgatatt | 1,090 bp |
| | WT | BD430, gagtaggtgcggtgataggc<br>BD431, tctgctcctcgaagaaccac | 2,876 bp |
| Bg_coco_pJB14 (*ΔbolT*) | FA | BD410, aggctcatgcgtgtgtctc<br>TIISS_A_rv, agtgacaacgtcgagcacag | 1,773 bp |
| | BA | BD411, gatggacgaggacttcatgc<br>TIISSD_B_fwd, cgttggctacccgtgatatt | 1,403 bp |
| | WT | BD410, aggctcatgcgtgtgtctc<br>BD411, gatggacgaggacttcatgc | 2,511 bp |
| Bg_coco_pBD92 (*ΔbolS*) | FA | BD463, tcagcccgtacatcaggtag<br>TIISS_A_rv, agtgacaacgtcgagcacag | 1,968 bp |
| | BA | BD462, ccctgacaaacgacttttcc<br>TIISSD_B_fwd, cgttggctacccgtgatatt | 1,558 bp |
| | WT | BD463, tcagcccgtacatcaggtag<br>BD462, ccctgacaaacgacttttcc | 3,483 bp |
| Bg_coco_pBD93 (*ΔbolU*) | FA | BD480, gatcaatccgcagttgaagg<br>TIISS_A_rv, agtgacaacgtcgagcacag | 2,225 bp |
| | BA | BD481, cgaggaagatctccagttcg<br>TIISSD_B_fwd, cgttggctacccgtgatatt | 1,902 bp |
| | WT | BD480, gatcaatccgcagttgaagg<br>BD481, cgaggaagatctccagttcg | 3,646 bp |

**Table S14**. Primer pairs and expected product sizes for colony PCRs to test for homologous recombination. (continued)

| | | | |
|---|---|---|---|
| Bg_coco_ pBD107 (*ΔbolV*) | FA | BD559, tcggaagtgccgaaatactg<br>TIISS_A_rv, agtgacaacgtcgagcacag | 2,560 bp |
| | BA | BD560, tcgtctccatgtcattccac<br>TIISSD_B_fwd, cgttggctacccgtgatatt | 1,722 bp |
| | WT | BD559, tcggaagtgccgaaatactg<br>BD560, tcgtctccatgtcattccac | 3,610 bp |
| Bg_coco_pBD95 (*ΔbolW*) | FA | BD492, gcatgctgatcgacatgaag<br>TIISS_A_rv, agtgacaacgtcgagcacag | 2,174 bp |
| | BA | BD493, aagttgcgcaggtacatcac<br>TIISSD_B_fwd, cgttggctacccgtgatatt | 1,693 bp |
| | WT | BD492, gcatgctgatcgacatgaag<br>BD493, aagttgcgcaggtacatcac | 3,432 bp |

All mutant strains were cultivated on 20 mL PDA agar overnight at 30 °C, extracted, analyzed using LC-MS and their metabolic profiles compared to the wild type.

### Bioactivity assays

#### Antibiotic assays

Bolagladin A (1 mg mL $^{-1}$, in DMSO) was tested against *Bacillus subtilis*, *Escherichia co*li, *Pseudomonas aeruginosa*, methicillin-resistant *Staphylococcus aureus* (MRSA), *Mycobacterium vaccae*, *Candida albicans* and P*enicillium notatum* as described previously.[14] Ciprofloxacin (5 µg mL$^{-1}$, in dH$_2$O) or amphotericin B (10 µg mL$^{-1}$, in DMSO and MeOH) was used as a positive control for test against bacteria respectively fungi (Table S15).

**Table S15.** Inhibitory effects of bolagladin (**1**) against selected strains.

| **Bacterial strains tested** | **Zone of inhibition in mm** | | |
|---|---|---|---|
| | **Bolagladin A (1) (1 mg mL$^{-1}$)** | **Ciprofloxacin (5 µg mL$^{-1}$)** | **DMSO** |
| *Bacillus subtilis* 6633 B1 | 13 p | 28 | 11 p |
| *Escherichia coli* 458 B4 | 15 p | 24/31p | 13 |
| *Pseudomonas aeruginosa SG 137 B9* | 0 | 27/34 p | 13 p |
| *Staphylococcus aureus* 134/94 R9 (MRSA) | 14 p | 0 | 11 p |
| *Enterococcus faecalis* 1528 R10 (VRE) | 17 | 16 | 12 p |
| *Mycobacterium vaccae* 10670 M4 | 18 | 22 p | 13 p |
| **Fungal strains tested** | **Bolagladin A (1) (1 mg mL$^{-1}$)** | **Amphotericin B (10 mg mL$^{-1}$)** | **DMSO** |
| *Candida albicans* H8 | 14 p | 20 | 0 |
| *Penicillium notatum* JP36 P1 | 15 p | 18 p | 12 p |

p – partial inhibition

### Swarming assays

MGY+M9 medium containing 1.5% agar was prepared, sterilized, and diluted with 1:1 water (0.7% MGY agar plates). 10 µL of culture with an $OD_{600}$ of 0.4, that was inoculated from an overnight *B. gladioli* pv. *cocovenenans* wild type and *ΔbolA* culture was pipetted in the middle of a 0.7% MGY agar plate and incubated at 30 °C. Similar swarming behavior was observed within 14 days of incubation.

### Drop-collapsing assay[15]

Droplets of water (20 µL) were placed on a hydrophobic surface (Parafilm 'M'). Bacterial cells of a single colony were transferred with a toothpick and resuspended in one droplet (Figure S6). The positive control, water *B. gladioli* HKI0739 (Bg739) droplets, resulted in a collapsed droplet. The control water, no additive and neither the *B. gladioli* pv. *cocovenenans* (*Bg*. pv. *coco*) nor the *B. gladioli* pv. *cocovenenans ΔbolG* lead to a collapse of the droplet. For visualization purposes 0.0025% crystal violet were added to the droplet. It had no influence on the shape of the droplet.

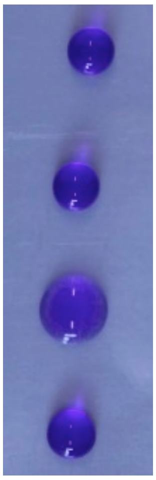


**Figure S6.** Droplet collapsing assay to test for surface activity of wild type *versus* mutant strains. pJB04, *B. gladioli* pv. *cocovenenans* pJB04; Bg739*, B. gladioli* HKI0739; *Bg*. pv. *coco*, *B. gladioli* pv. *cocovenenans*).

### CAS agar test

To test for siderophore production of *B. gladioli* pv. *cocovenenans* we used CAS agar plates.[13] 100 µL of an overnight culture was pipetted onto the agar and incubated for a varying periods at 30 °C. A discoloration and halo formation colonies indicates siderophore activity. No or minor differences were observed when comparing the halo formation of *B. gladioli* pv. *cocovenenans* wild type, *B. gladioli* pv. *cocovenenans ΔbolA* or *B. gladioli* pv. *cocovenenans ΔbolW* (producer of **3** and **4**) colonies (**Fehler! Verweisquelle konnte nicht gefunden werden.**). The discoloration likely occurs due to the production of gladiobactin. Spotting of pure compound did not result in discoloration of the agar.

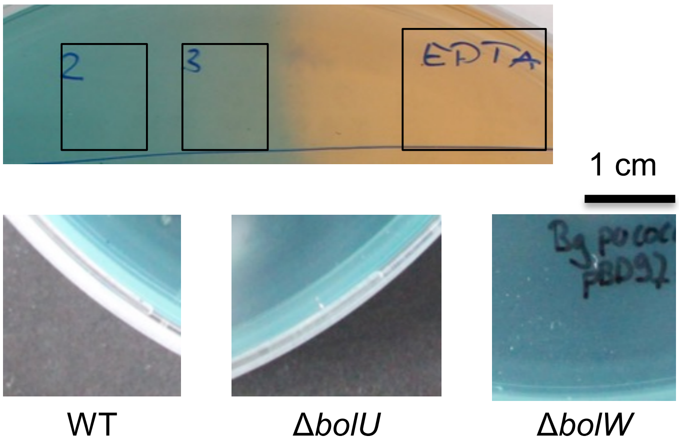


**Figure 7.** *B. gladioli* pv. *cocovenenans* or bolagladin incubated on CAS agar plates at 30 °C, respectively, 20 °C for 24 h.
Spotting of 2 = bolagladin A (1) 0.1 mg $ml^{-1}$; 3 = bolagladin A (1) 0.5 mg $ml^{-1}$. EDTA was used as positive control. No noteworthy differences have been observed between *B. gladioli* pv. *cocovenenans* wild type (WT), *B. gladioli* pv. *cocovenenans ΔbolU* (no bolagladin production), *B. gladioli* pv. *cocovenenans ΔbolW* (producer of 3 and 4).

### References


[1] E. Yabuuchi, Y. Kosako, H. Oyaizu, I. Yano, H. Hotta, Y. Hashimoto, T. Ezaki, M. Arakawa, *Microbiol. Immunol.* **1992**, *36*, 1251-1275.
[2] K. Blin, S. Shaw, K. Steinke, R. Villebro, N. Ziemert, S. Y. Lee, M. H. Medema, T. Weber, *Nucleic Acids Res.* **2019**, *47*, W81-W87.
[3] L. Zimmermann, A. Stephens, S.-Z. Nam, D. Rau, J. Kübler, M. Lozajic, F. Gabler, J. Söding, A. N. Lupas, V. Alva, *J. Mol. Biol.* **2018**, *430*, 2237-2243.
[4] N. Ziemert, S. Podell, K. Penn, J. H. Badger, E. Allen, P. R. Jensen, *PLoS One* **2012**, *7*.
[5] K. Katoh, D. M. Standley, in *Methods Mol. Biol.*, Springer, **2014**, pp. 131-146.
[6] J. Trifinopoulos, L.-T. Nguyen, A. von Haeseler, B. Q. Minh, *Nucleic Acids Res.* **2016**, *44*, W232-W235.
[7] R. Fujii, Y. Matsu, A. Minami, S. Nagamine, I. Takeuchi, K. Gomi, H. Oikawa, *Org. Lett.* **2015**, *17*, 5658-5661.
[8] K. Williams, A. J. Szwalbe, N. P. Mulholland, J. L. Vincent, A. M. Bailey, C. L. Willis, T. J. Simpson, R. J. Cox, *Angew. Chem. Int. Ed.* **2016**, *55*, 6784-6788.
[9] B. Bonsch, V. Belt, C. Bartel, N. Duensing, M. Koziol, C. M. Lazarus, A. M. Bailey, T. J. Simpson, R. J. Cox, *Chem. Commun.* **2016**, *52*, 6777-6780.
[10] aF. E. Cole, M. G. Kalyanpur, C. M. Stevens, *Biochemistry* **2002**, *12*, 3346-3350; bU. Thomas, M. G. Kalyanpur, C. M. Stevens, *Biochemistry* **1966**, *5*, 2513-2516.
[11] M. J. Sullivan, N. K. Petty, S. A. Beatson, *Bioinformatics* **2011**, *27*, 1009-1010.
[12] K. Ishida, T. Lincke, C. Hertweck, *Angew. Chem. Int. Ed.* **2012**, *51*, 5470-5474.
[13] B. Schwyn, J. B. Neilands, *Anal. Biochem.* **1987**, *160*, 47-56.
[14] a) R. Abdou, K. Scherlach, H.-M. Dahse, I. Sattler, C. Hertweck, *Phytochemistry* **2010**, *71*, 110-116; b) K. Scherlach, L. P. Partida-Martinez, H.-M. Dahse, C. Hertweck, *J. Am. Chem. Soc.* **2006**, *128*, 11529-11536.
[15] D. K. Jain, D. L. Collins-Thompson, H. Lee, J. T. Trevors, *J. Microbiol. Methods* **1991**, *13*, 271-279.
[16] E. Pauwelyn, C.-J. Huang, M. Ongena, V. Leclère, P. Jacques, P. Bleyaert, H. Budzikiewicz, M. Schäfer, M. Höfte, *Mol. Plant-Microbe Interact.* **2013**, *26*, 585-598.

### NMR data and spectra

### Physicochemical data

#### Bolagladin A (1)

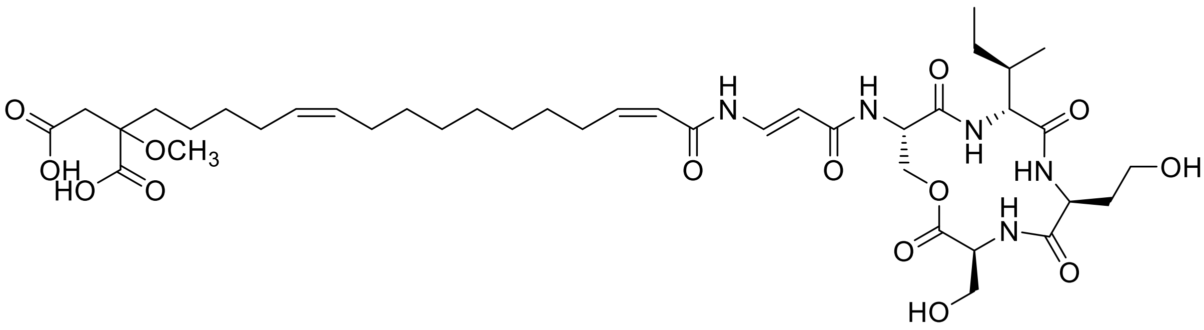

(–)ESI HR-MS: *m/z* 836.4315 [M–H]$^-$ (calcd. $C_{40}H_{62}N_5O_{14}$ 836.4299)

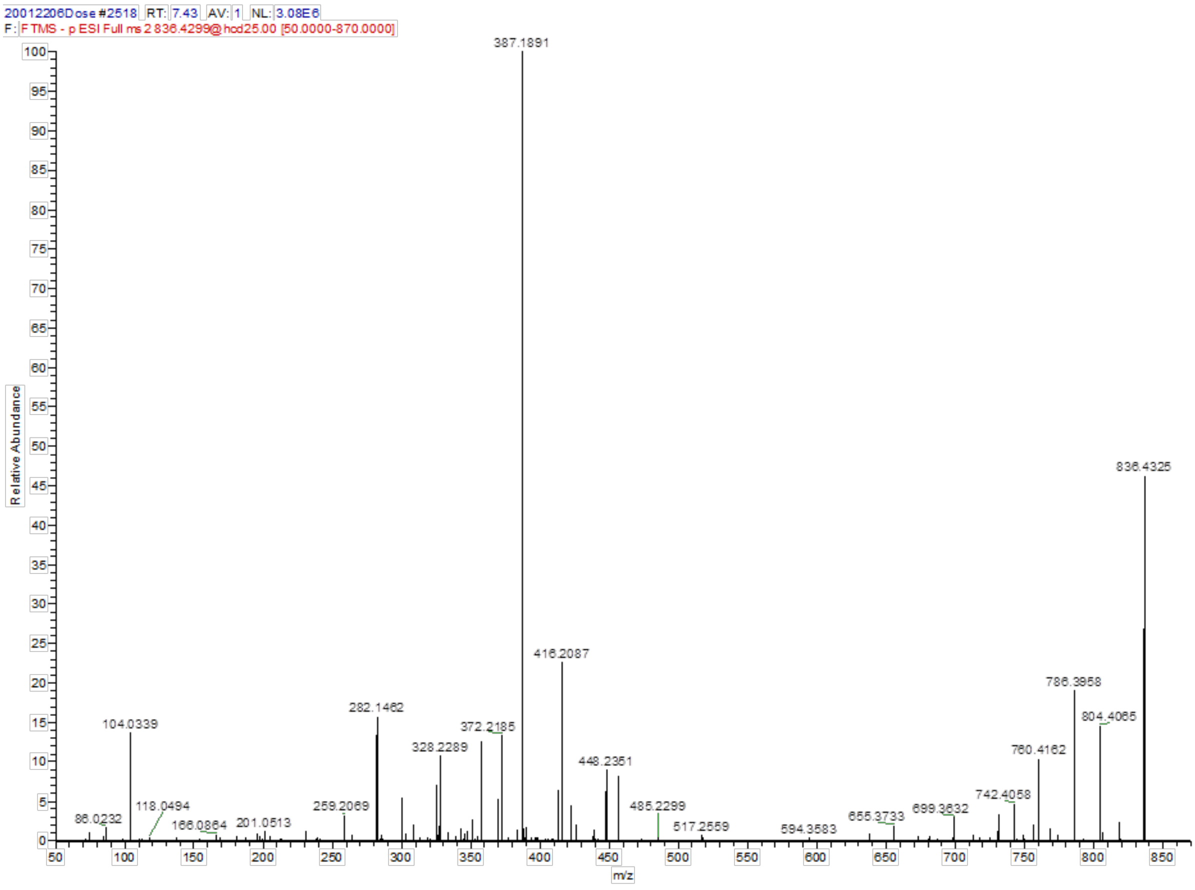


**Figure S8.** (–)ESI MS/MS spectrum of bolagladin A (**1**).

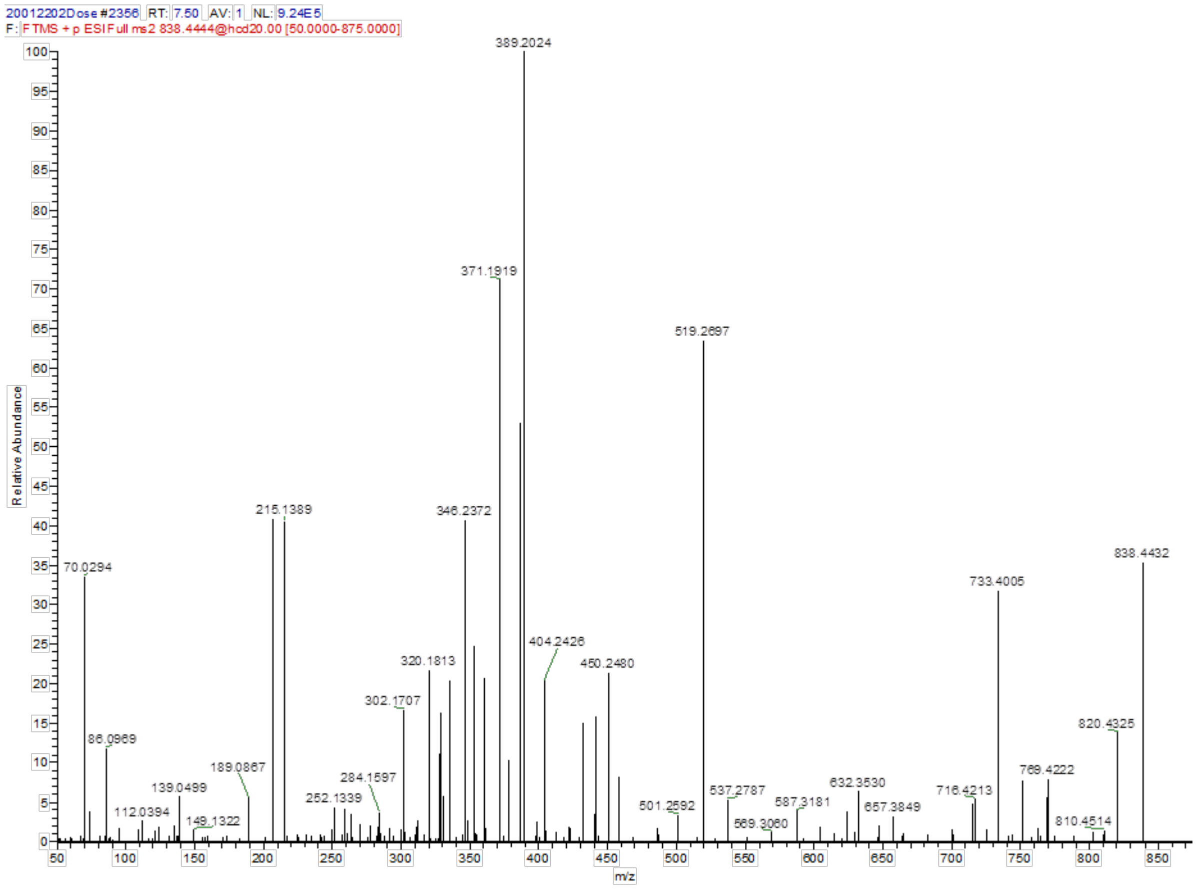


**Figure S9.** (+)ESI MS/MS spectrum of bolagladin A (**1**).

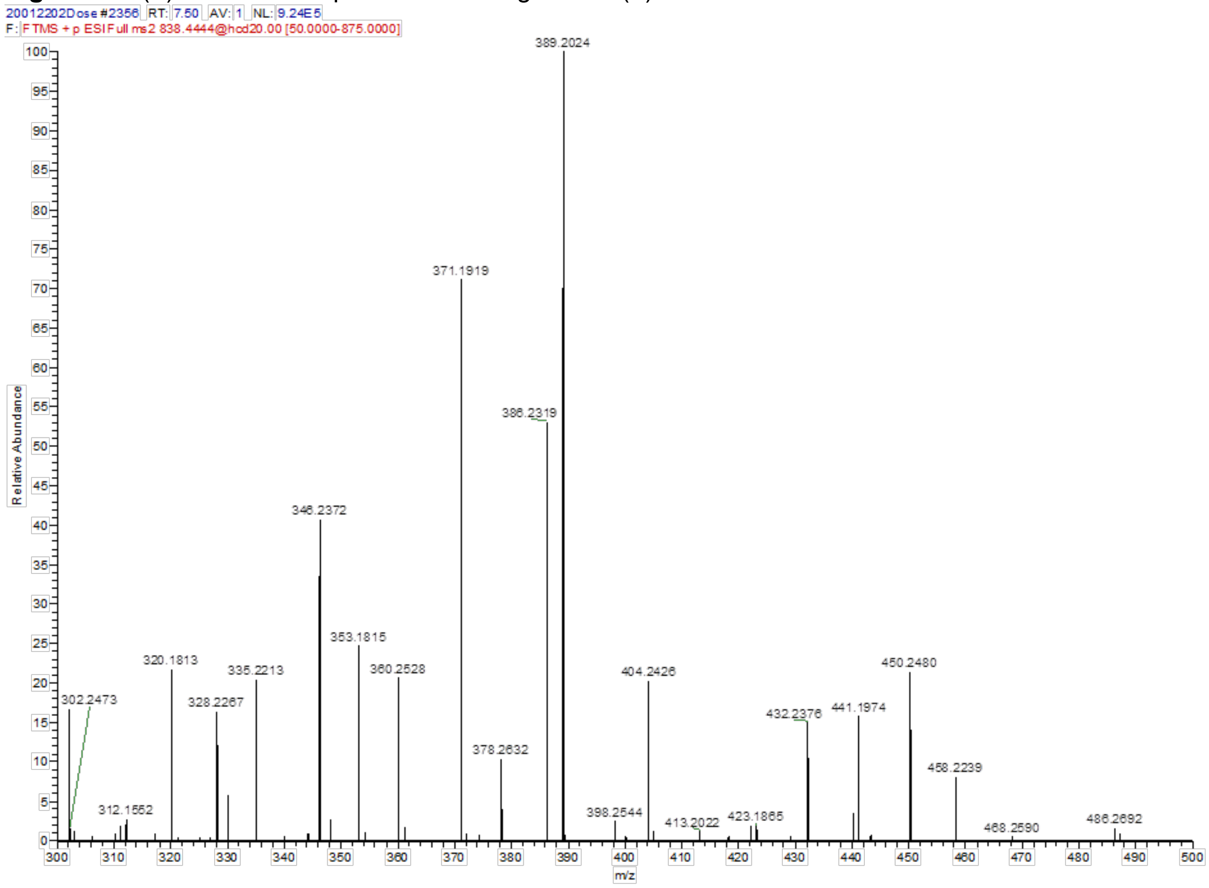


**Figure S10.** (+)ESI MS/MS spectrum of bolagladin A (**1**), mass range *m/z* 300–500.

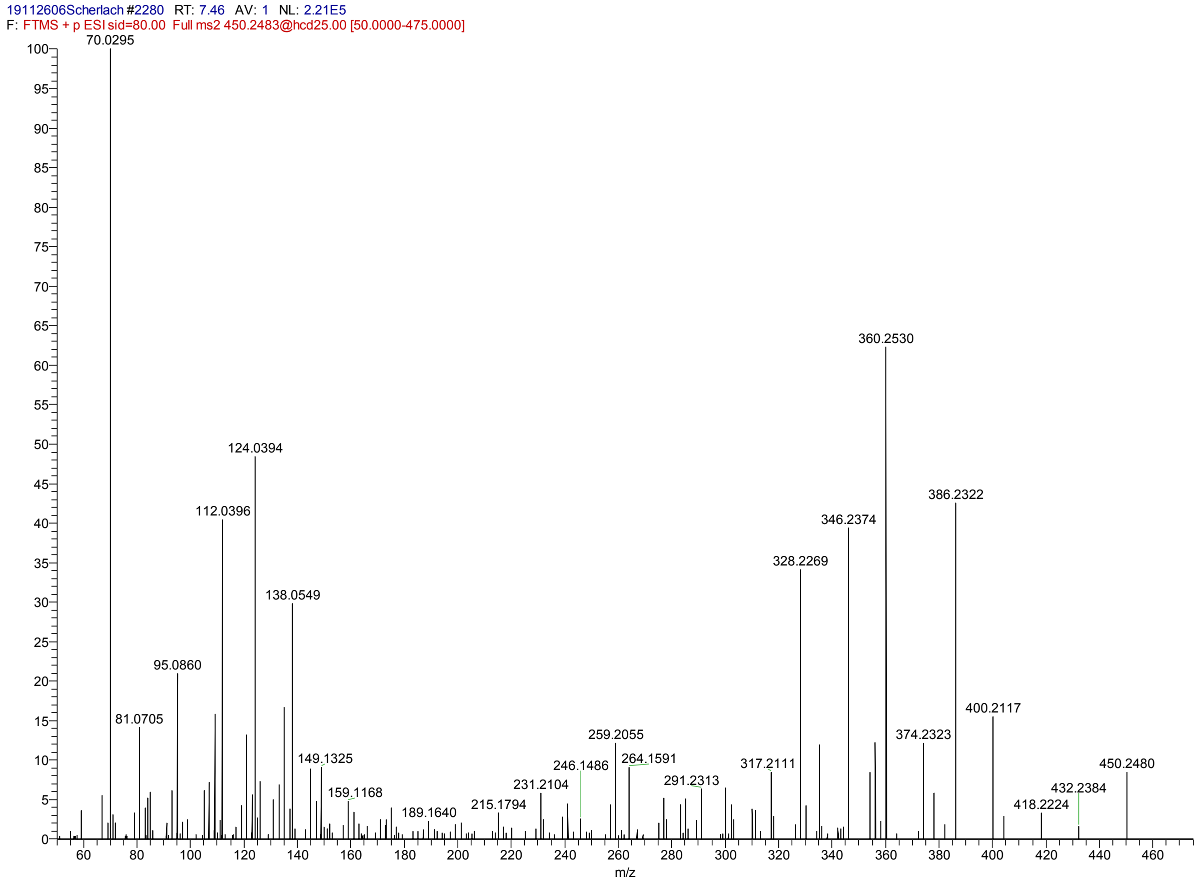


**Figure S11.** (+)ESI MS/MS spectrum of the fragment *m/z* 450 amu derived from bolagladin A (**1**).

**Table S16.** NMR data of bolagladin A (**1**).

| | | **Bolagladin A (1)** | |
|---|---|---|---|
| | **Position** | **$^{13}C$** | **$^{1}H$ (*J* in Hz)** |
| L-Serine 1 | 1 | 168.8 | - |
| | 2 | 54.4 | 4.33 m |
| | 3 | 60.7 | 3.67 m |
| | | | 3.57 m |
| | NH | - | 7.20 d (*7.8*) |
| L-Homoserine | 1 | 171.0 | - |
| | 2 | 51.2 | 4.22 m |
| | 3 | 33.5 | 1.78 m |
| | 4 | 57.3 | 3.45 m |
| | | | 3.37 m |
| | NH | - | 8.62 d (*7.7*) |
| D-Isoleucine | 1 | 171.8 | - |
| | 2 | 57.5 | 4.11 t (*10.1*) |
| | 3 | 34.1 | 1.73 m |
| | 4 | 24.2 | 1.42 m |
| | | | 1.04 m |
| | 5 | 10.2 | 0.80 t (*7.4*) |
| | 6 | 15.0 | 0.77 d (*6.7*) |
| | NH | - | 7.38 brd |
| L-Serine 2 | 1 | 169.5 | - |
| | 2 | 52.5 | 4.56 m |
| | 3 | 64.8 | 4.22 m |
| | NH | - | 8.45 brd |
| Dehydro-*β*-alanine | 1 | 166.2 | - |
| | 2 | 103.1 | 5.76 d (*14.0*) |
| | 3 | 134.7 | 7.72 dd (*14.0; 11.2*) |
| | NH | - | 10.73 d (*10.8*) |

| | | | |
|---|---|---|---|
| Fatty acid | 1 | 163.9 | - |
| | 2 | 121.3 | 5.92 d (*11.5*) |
| | 3 | 149.1 | 6.20 dt (*11.5;7.5*) |
| | 4 | 28.3 | 2.63 m |
| | | | 1.35 m |
| | 5-9 | Not assigned | Not assigned |
| | 10 | 26.6 | 1.92 m |
| | 11 | 129.8 | 5.30 m |
| | 12 | 129.3 | 5.30 m |
| | 13 | 26.5 | 1.92 m |
| | 14 | 29.2 | 1.25 m |
| | 15 | 22.4 | 1.23 m |
| | | | 1.17 m |
| | 16 | 33.9 | 1.73 m |
| | 17 | 79.8 | - |
| | 18 | 39.0 | 2.63 |
| | 19 | 171.2 | - |
| | 20 | 173.2 | - |
| | 17-$OCH_3$ | 50.9 | 3.16 s |

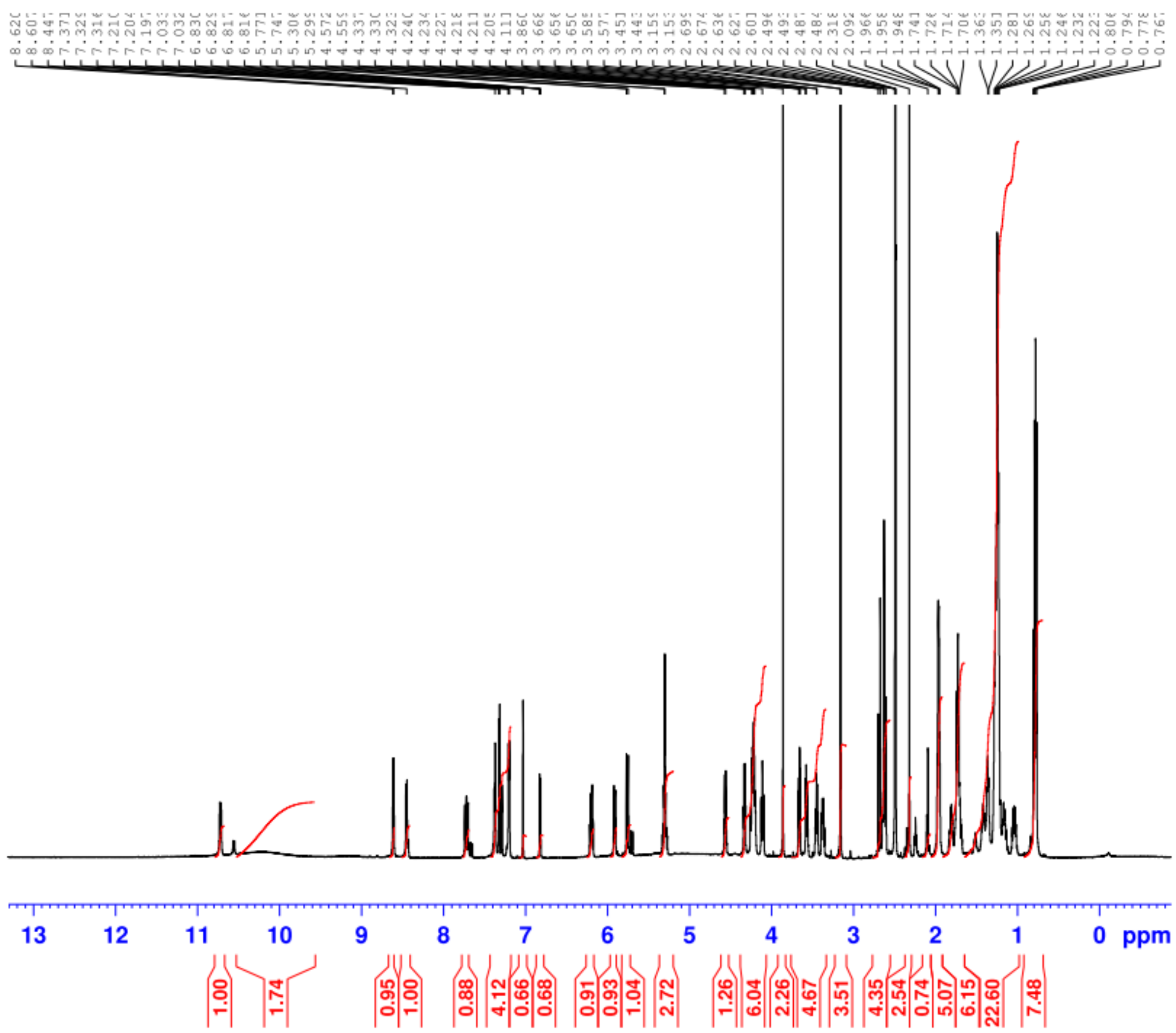

**Figure S12.** $^{1}$H NMR spectrum of bolagladin A (**1**).

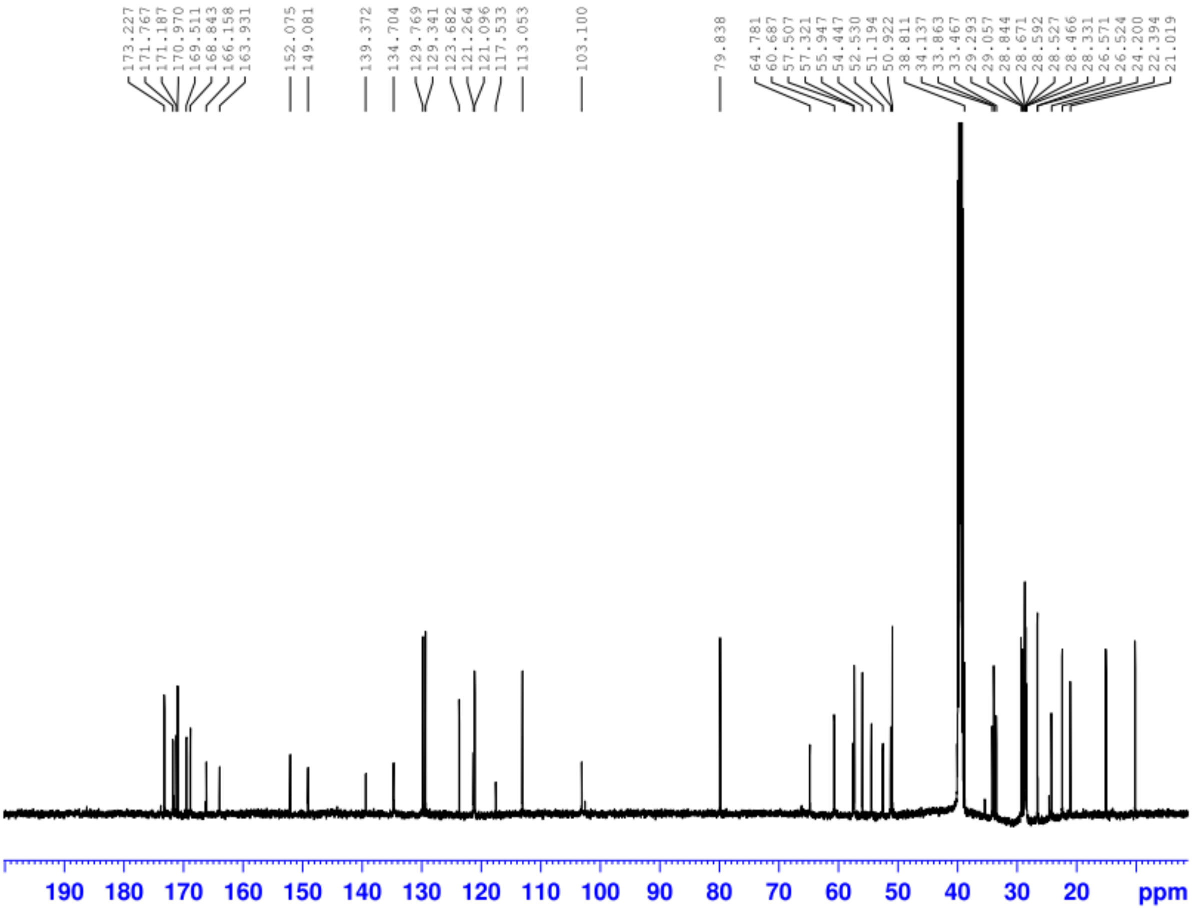


**Figure S13.** $^{13}C$ NMR spectrum of bolagladin A (**1**).

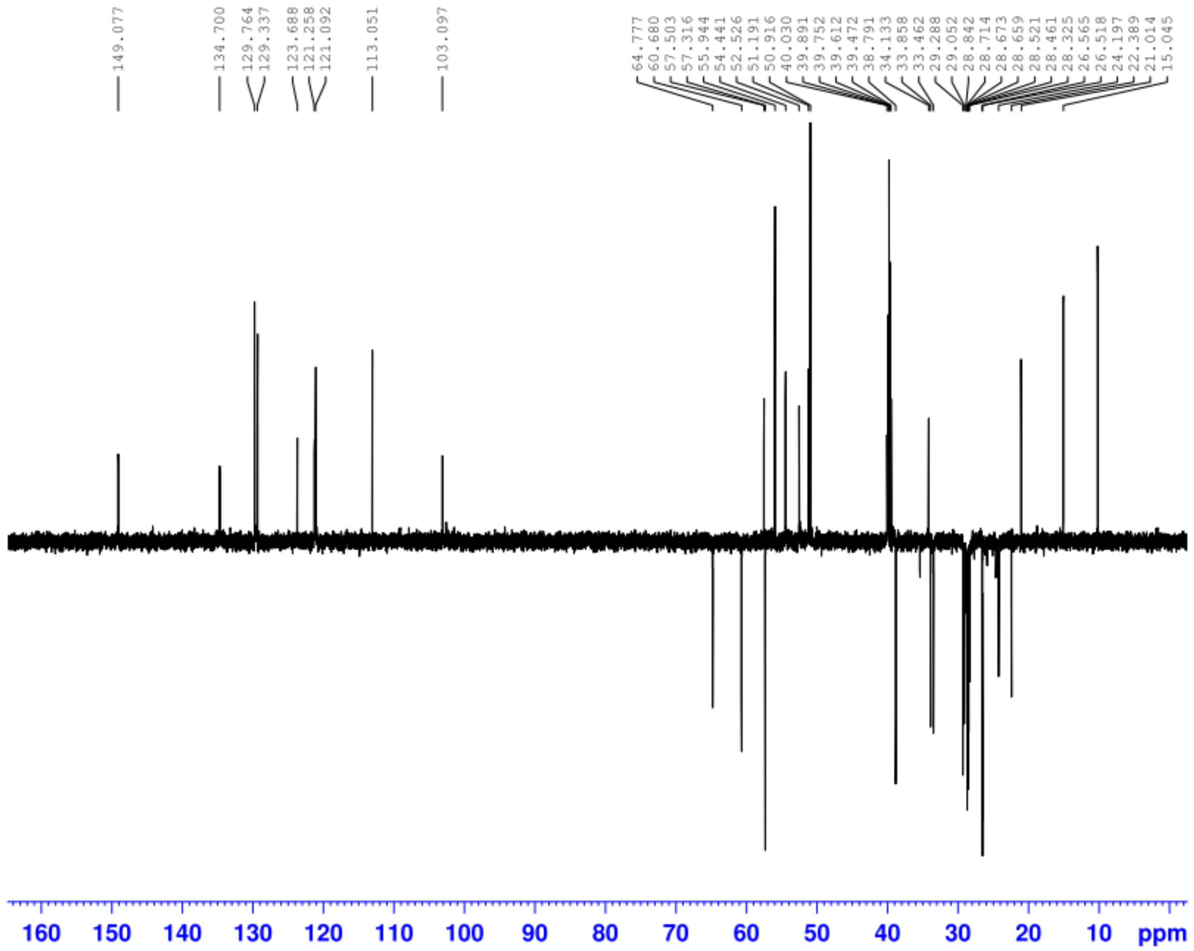

**Figure S14.** DEPT-135 NMR spectrum of bolagladin A (**1**).

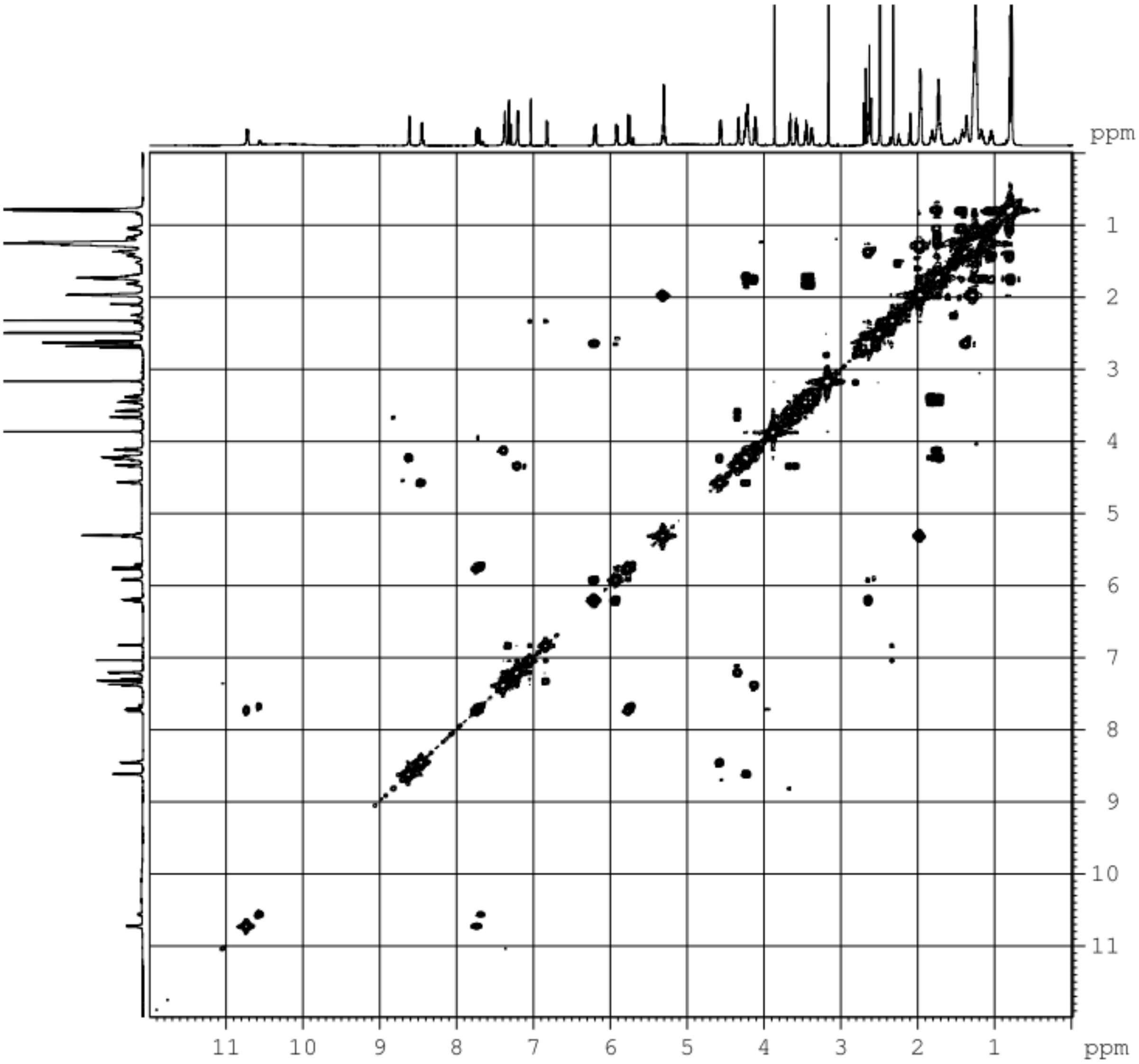


**Figure S15**. H,H-COSY spectrum of bolagladin A (**1**).

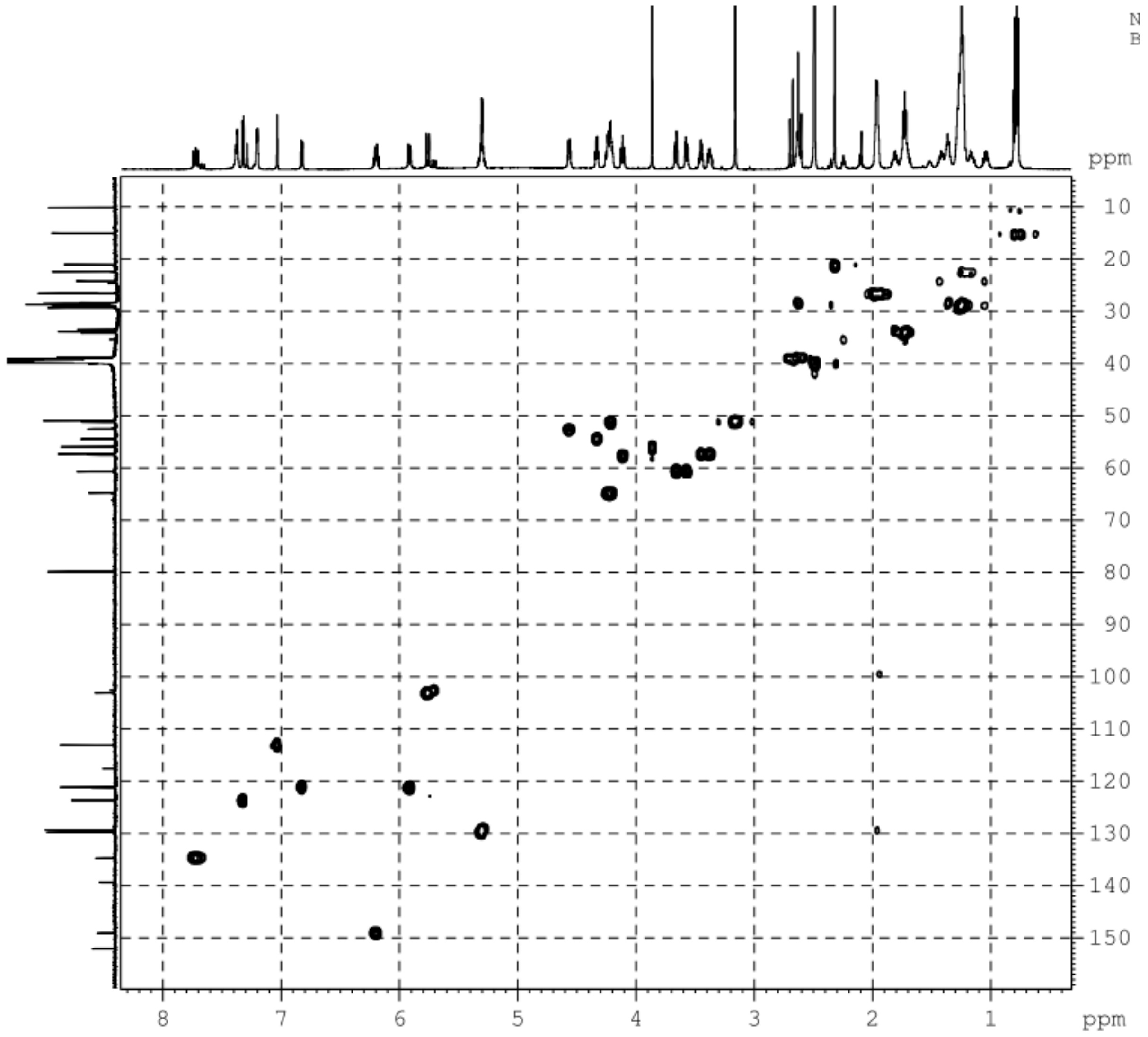


**Figure S16**. HSQC spectrum of bolagladin A (**1**).

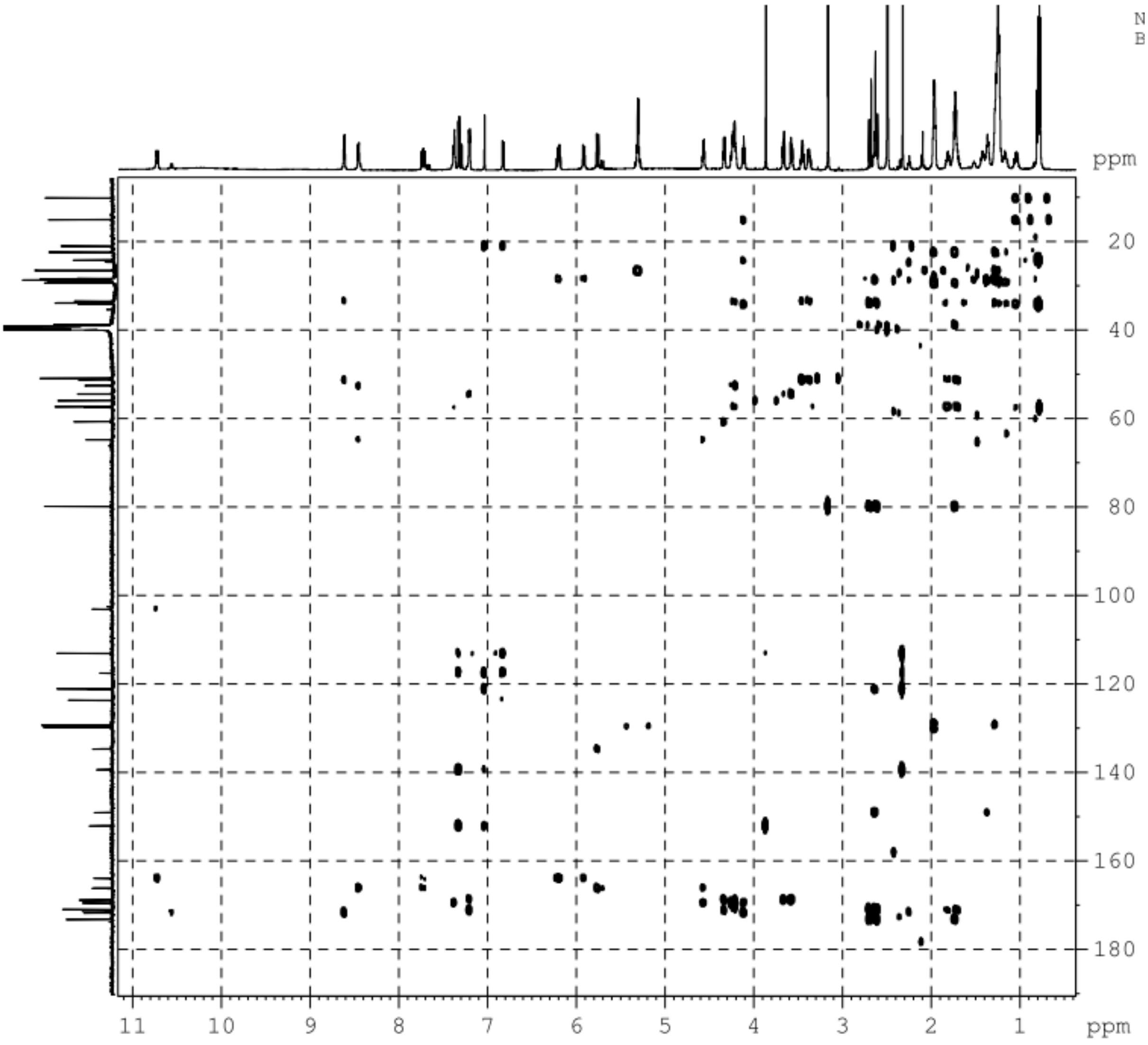


**Figure S17**. HMBC spectrum of bolagladin A (**1**).

**Bolagladin B (2)**

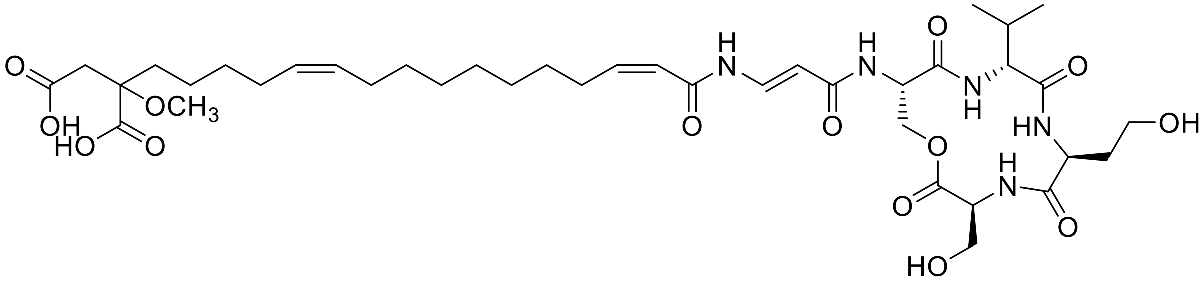

(–)ESI HR-MS: *m/z* 822.4163 [M–H]$^{-}$ (calcd. $C_{39}H_{60}N_5O_{14}$ 822.4142)

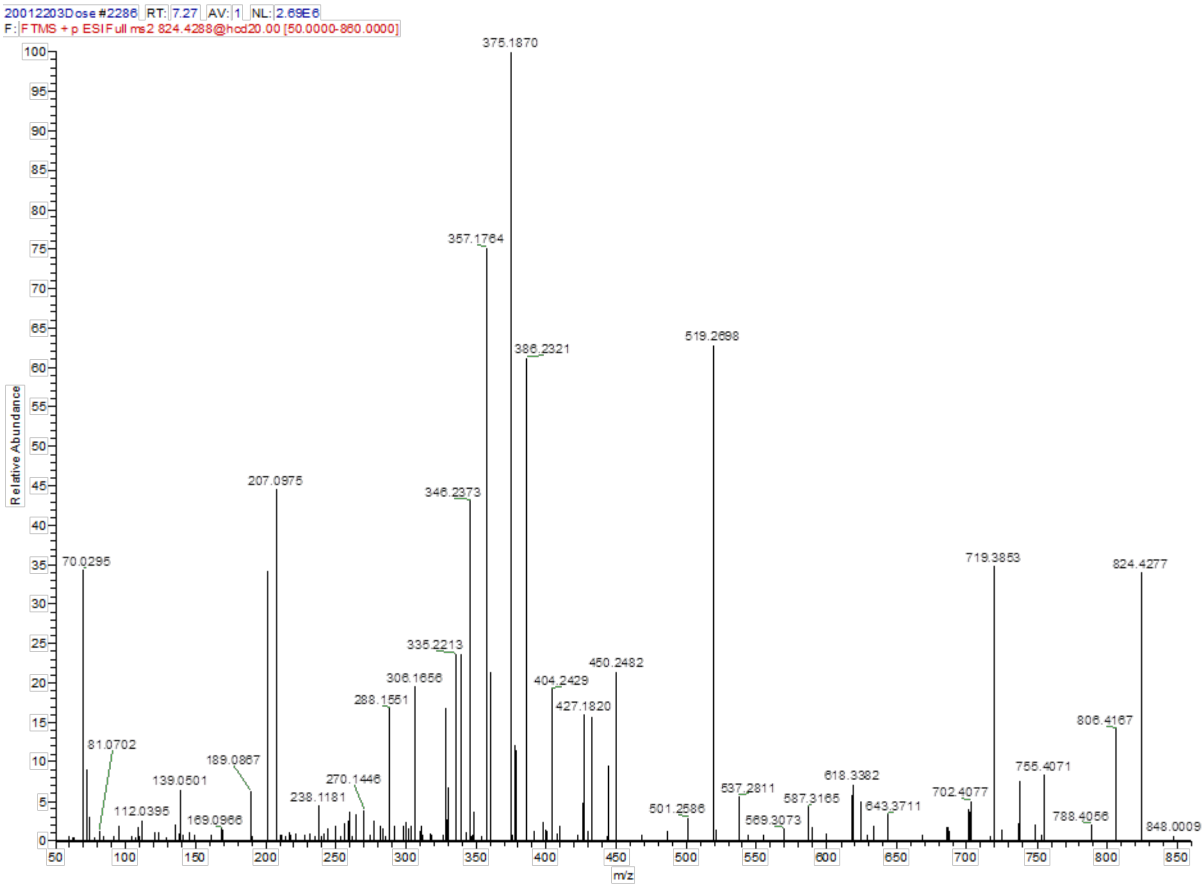


**Figure S18**. (+)ESI MS/MS spectrum of bolagladin B (**2**).

**Table S17**. NMR data of bolagladin B (**2**).

| | | **Bolagladin B (2)** | |
|---|---|---|---|
| | **Position** | **$^{13}$C** | **$^{1}$H (*J* in Hz)** |
| L-Serine 1 | 1 | 169.4 | - |
| | 2 | 55.0 | 4.32 m |
| | 3 | 61.0 | 3.68 m |
| | | | 3.57 m |
| | NH | - | 7.22 d (*8.0*) |
| L-Homoserine | 1 | 1 | - |
| | 2 | 51.5 | 4.22 m |
| | 3 | n. d. | 1.80 m |
| | 4 | 57.8 | 3.45 m |
| | | | 3.37 m |
| | NH | - | 8.56 d (*7.7*) |
| D-Valine | 1 | 171.7 | - |
| | 2 | 59.6 | 4.07 t (*10.0*) |
| | 3 | 29.0 | 1.23 m |
| | 4 | 19.0 | 0.81 d (*6.5*) |
| | 5 | 19.5 | 0.81 d (*6.5*) |
| | NH | - | 7.29 d (*8.9*) |
| L-Serine 2 | 1 | 169.9 | - |
| | 2 | 53.1 | 4.54 m |
| | 3 | 65.3 | 4.23 m |
| | NH | - | 8.33 d (*7.3*) |
| Dehydro-*β*-alanine | 1 | 166.7 | - |
| | 2 | 103.6 | 5.72 d (*14.0*) |
| | 3 | 135.2 | 7.72 dd (*13.8; 11.3*) |
| | NH | - | 10.53 d (*11.1*) |
| Fatty acid | 1 | 164.4 | - |
| | 2 | 121.7 | 5.87 d (*11.4*) |
| | 3 | 149.8 | 6.21 dt (*11.3; 7.5*) |

| | | |
|---|---|---|
| 4 | 28.7 | 2.63 m |
| | | 1.35 m |
| 5-9 | Not assigned | Not assigned |
| 10 | 27.1 | 1.96 m |
| 11 | 130.2 | 5.30 m |
| 12 | 129.9 | 5.30 m |
| 13 | 27.1 | 1.96 m |
| 14 | Not assigned | Not assigned |
| 15 | Not assigned | Not assigned |
| 16 | 33.9 | 1.70 m |
| 17 | 80.2 | - |
| 18 | 39.0 | 2.63 |
| 19 | 172.3 | - |
| 20 | 174.0 | - |
| 17-$OCH_3$ | 51.3 | 3.13 s |

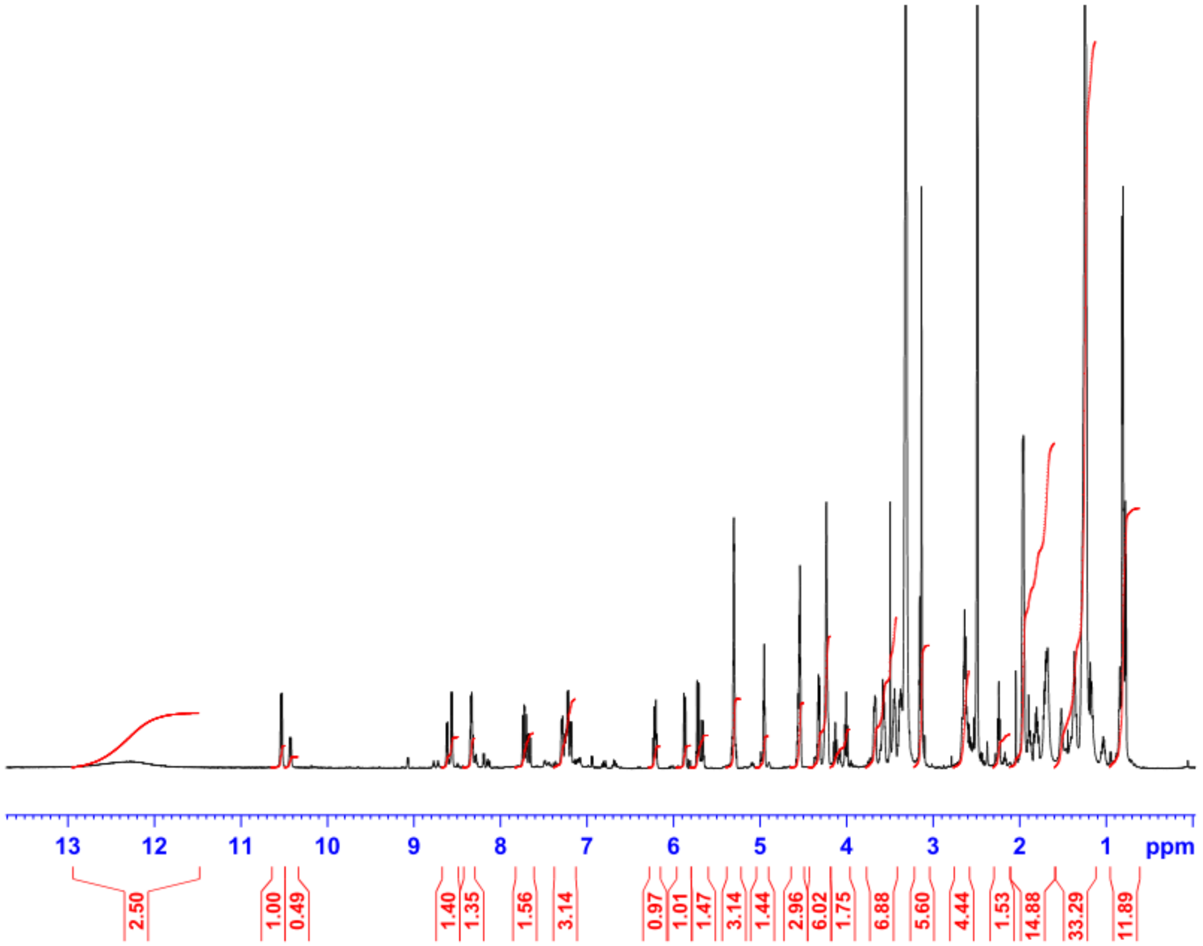


**Figure S19**. $^{1}$H NMR spectrum of bolagladin B (**2**).

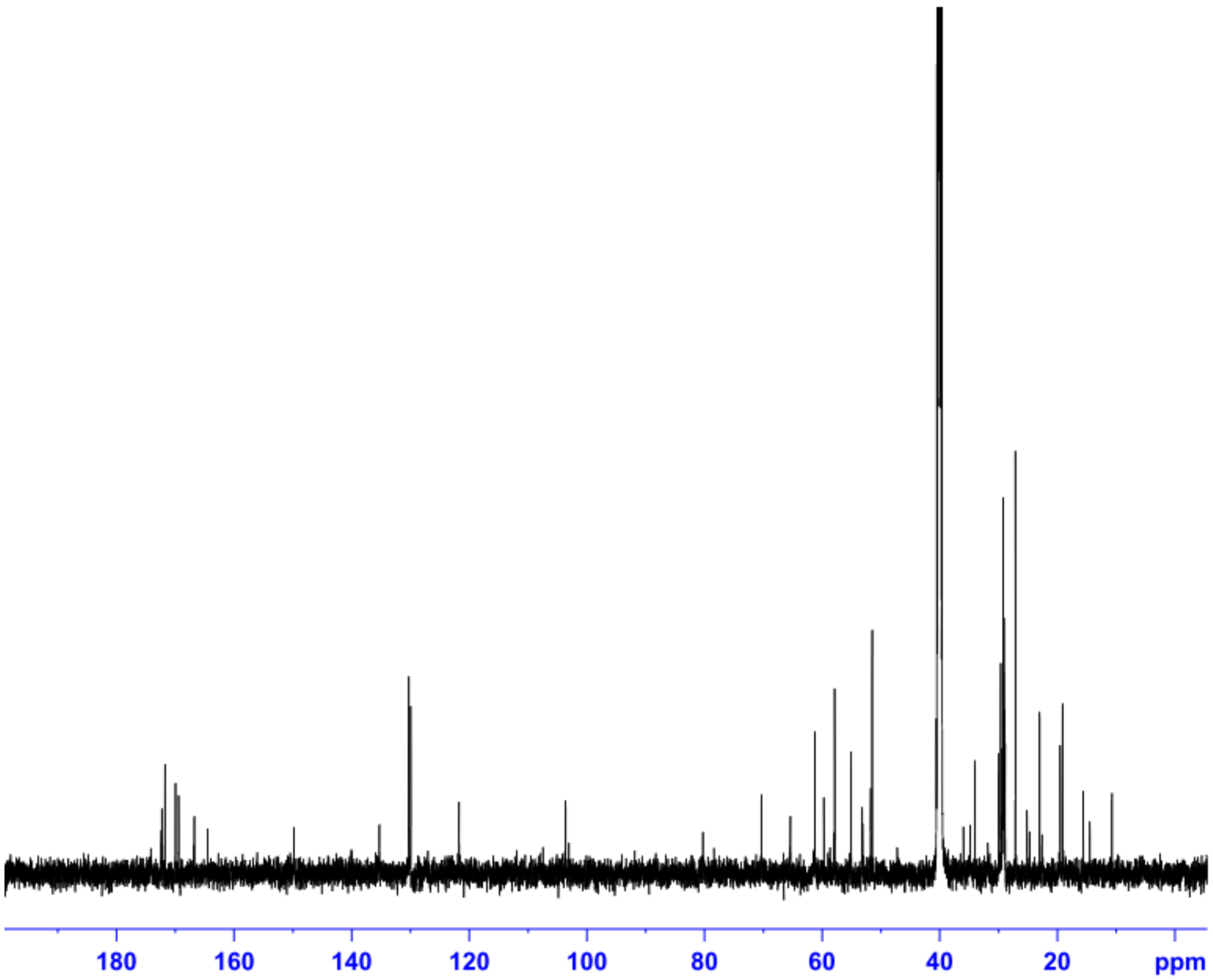


**Figure S20.** $^{13}$C NMR spectrum of bolagladin B (**2**).

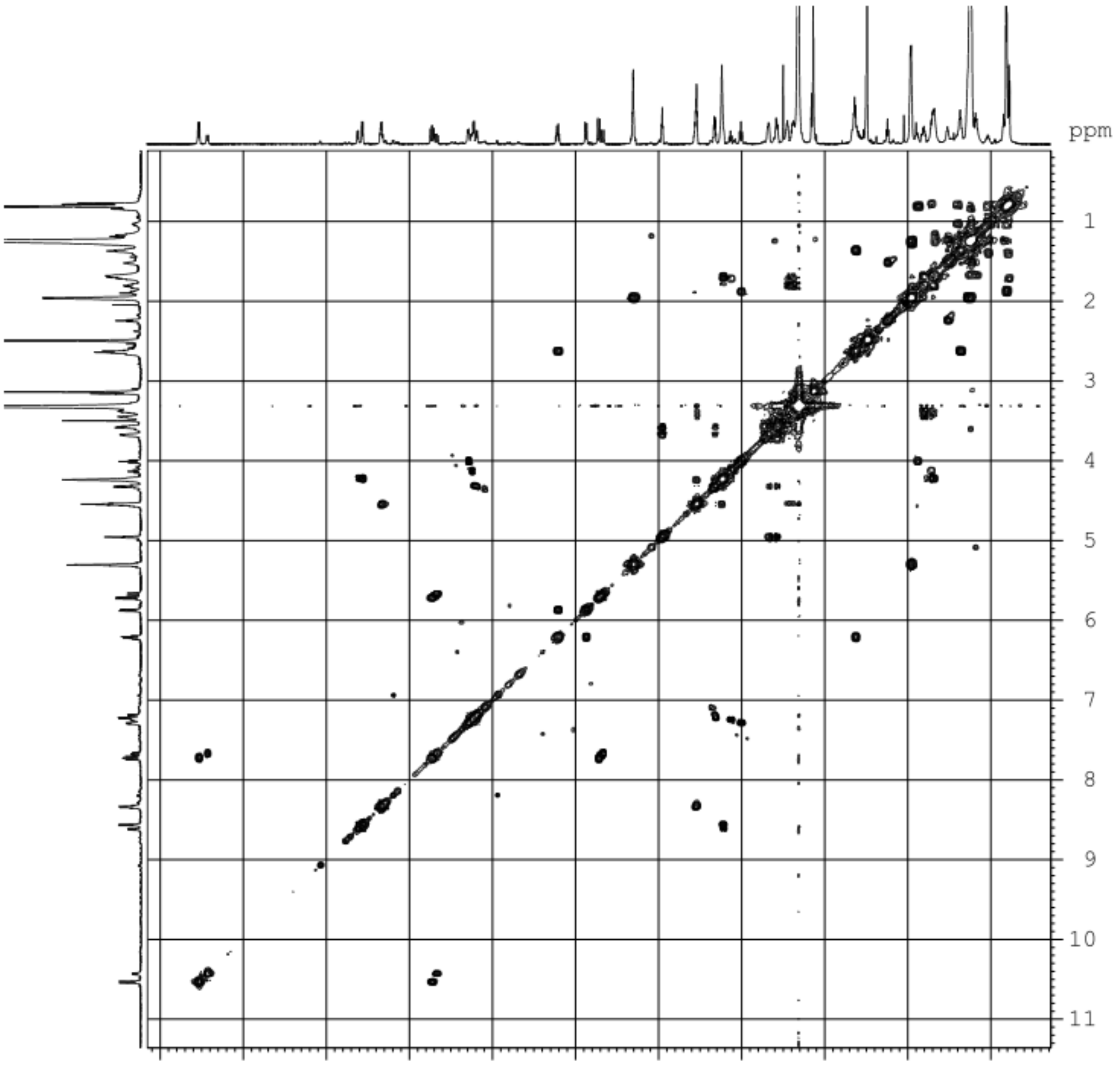


**Figure S21**. H,H-COSY spectrum of bolagladin B (**2**).

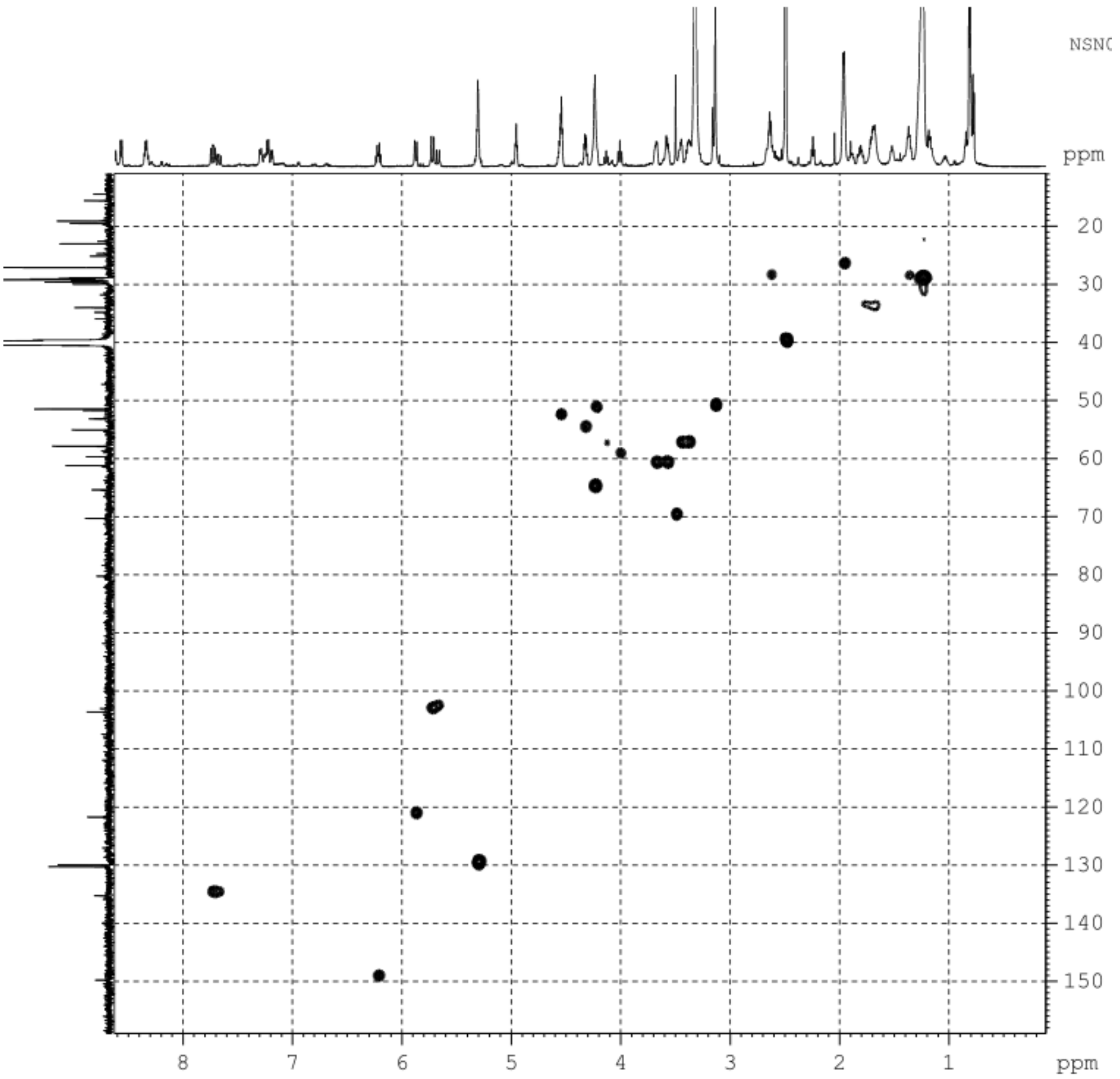


**Figure S22.** HSQC spectrum of bolagladin B (**2**).

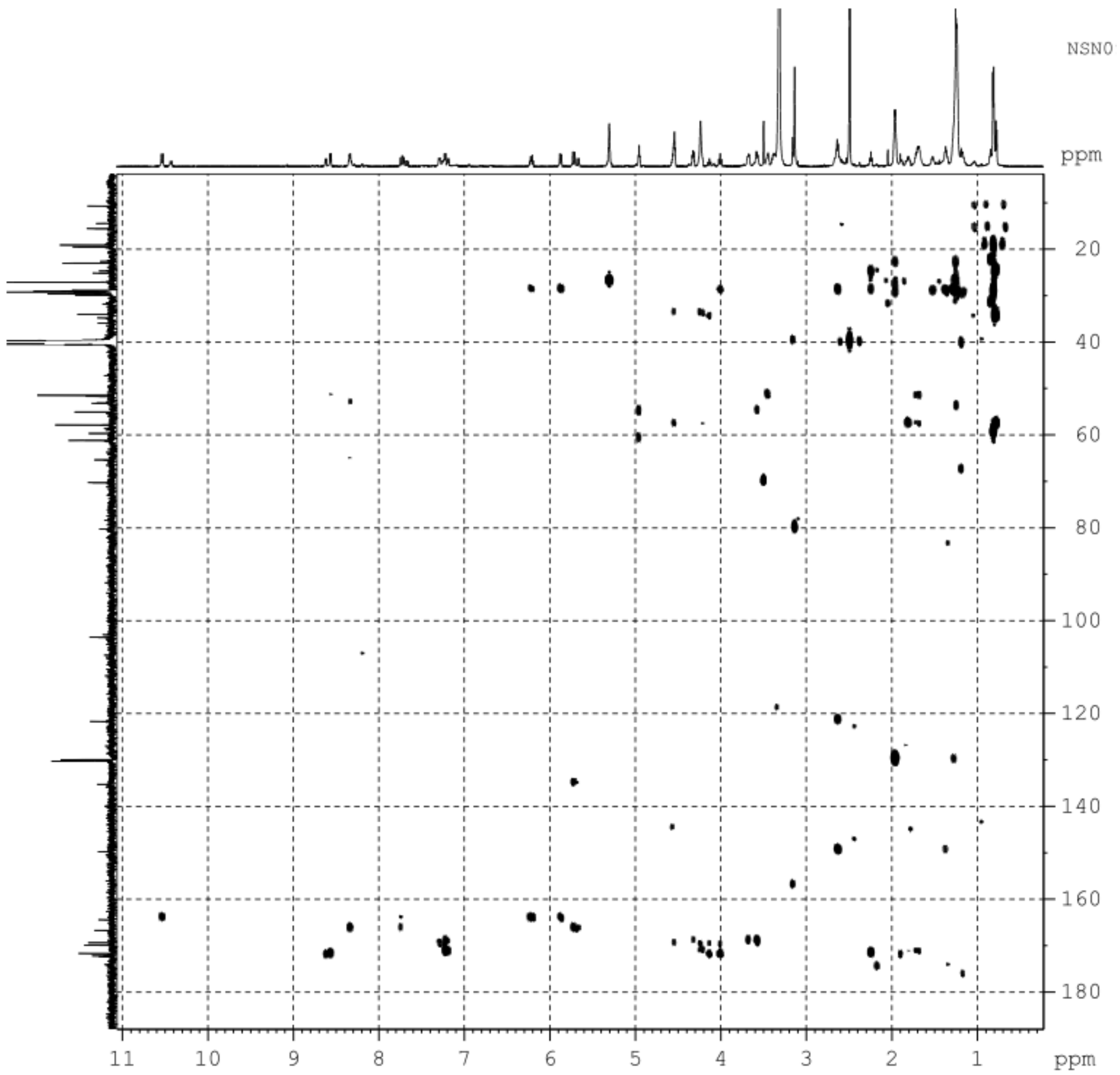


**Figure S23.** HMBC spectrum of bolagladin B (**2**).

**Bolagladin M749 (3)**

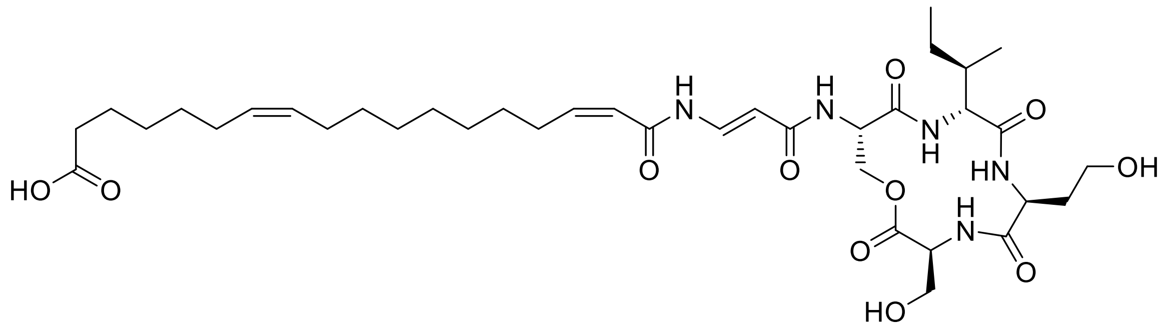


(–)ESI HR-MS: *m/z* 748.4157 $[M–H]^-$ (calcd. $C_{37}H_{58}N_5O_{11}$ 748.4138)

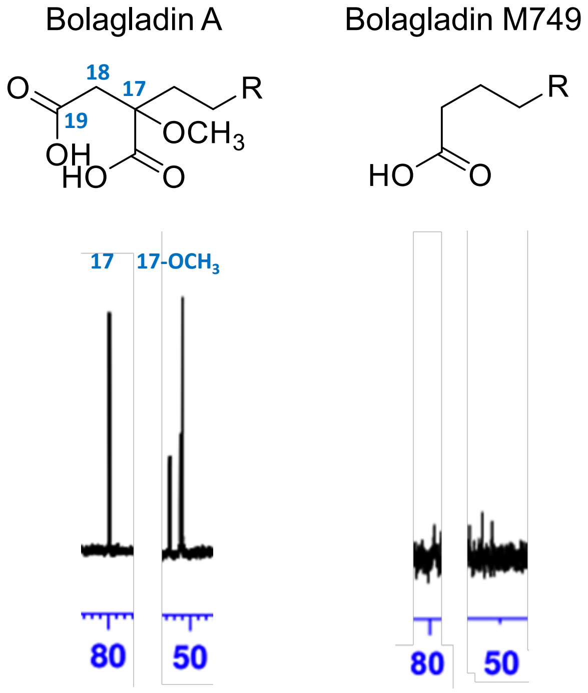


**Figure 24.** Selected diagnostic $^{13}C$ NMR signals of bolagladin A (1)and bolagladin M749 (**3**).

It should be noted that the signals for C18 and C19 in bolagladin A are overlapping with either the solvent signal (C18) or amide carbon signals (C19).

**Table 18.** NMR data of bolagladin M749 (**3**).

| | | Bolagladin M749 (3) | |
|---|---|---|---|
| | **Position** | **$^{13}C$** | **$^{1}H$ (*J* in Hz)** |
| L-Serine 1 | 1 | 169.3 | - |
| | 2 | 54.4 | 4.33 m |
| | 3 | 60.7 | 3.67 m |
| | | | 3.57 m |
| | NH | - | 7.19 |
| L-Homoserine | 1 | 171.2 | - |
| | 2 | 51.2 | 4.24 m |
| | 3 | 33.5 | 1.73 m |
| | 4 | 57.3 | 3.45 m |
| | | | 3.37 m |
| | NH | - | 8.62 d (*7.7*) |
| D-Isoleucine | 1 | 171.8 | - |
| | 2 | 57.3 | 4.12 t (*10.0*) |
| | 3 | 33.9 | 1.73 m |
| | 4 | 24.2 | 1.46 m |
| | | | 1.04 m |
| | 5 | 10.2 | 0.80 t |
| | 6 | 15.0 | 0.77 d |
| | NH | - | 7.23 brd |
| L-Serine 2 | 1 | 169.4 | - |
| | 2 | 52.5 | 4.57 m |
| | 3 | 64.9 | 4.23 m |
| | NH | - | 8.34 d |
| Dehydro-*β*-alanine | 1 | 166.2 | -. |
| | 2 | 103.2 | 5.73 d |
| | 3 | 134.7 | 7.72 dd (*14.0*; *11.2*) |
| | NH | - | 10.53 d (*10.8*) |
| Fatty acid | 1 | 163.9 | - |

| | | |
|---|---|---|
| 2 | 121.2 | 5.87 d (*11.5*) |
| 3 | 149.1 | 6.20 dt (*11.5;7.5*) |
| 4 | 28.4 | 2.63 m |
| | | 1.35 m |
| 5-9 | Not assigned | Not assigned |
| 10 | 26.6 | 1.95 m |
| 11 | 129.7 | 5.30 m |
| 12 | 129.5 | 5.30 m |
| 13 | 26.6 | 1.95 m |
| 14 | 29.2 | 1.25 m |
| 15 | 22.0 | 1.12-1.25 m |
| 16 | 24.2[a] | 1.46 m[a] |
| 17 | 31.6[a] | 2.10-2.30 m[a] |
| 18 | - | - |
| 19 | - | - |
| 20 | 174.4[a] | - |
| 17-$OCH_3$ | - | - |

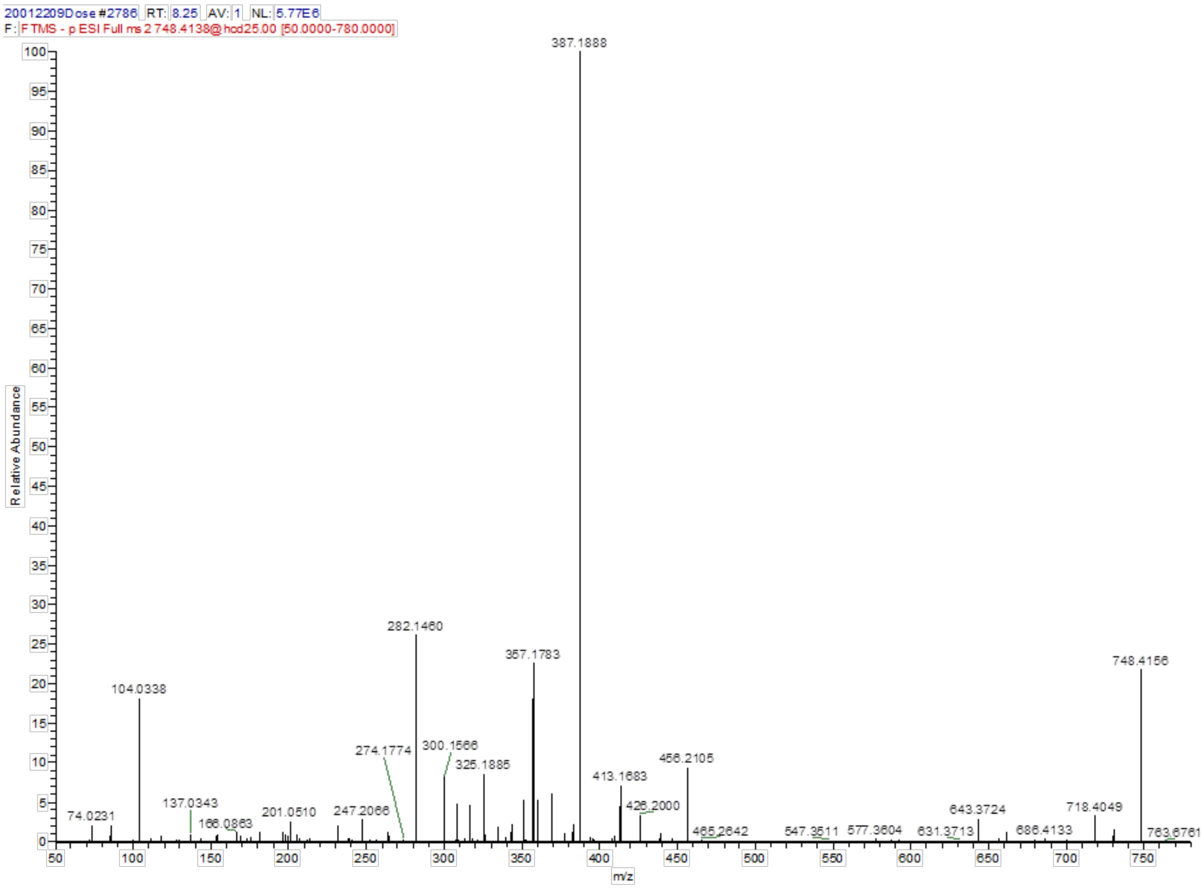


**Figure S25.** (–)ESI MS/MS spectrum of bolagladin M749 (**3**).

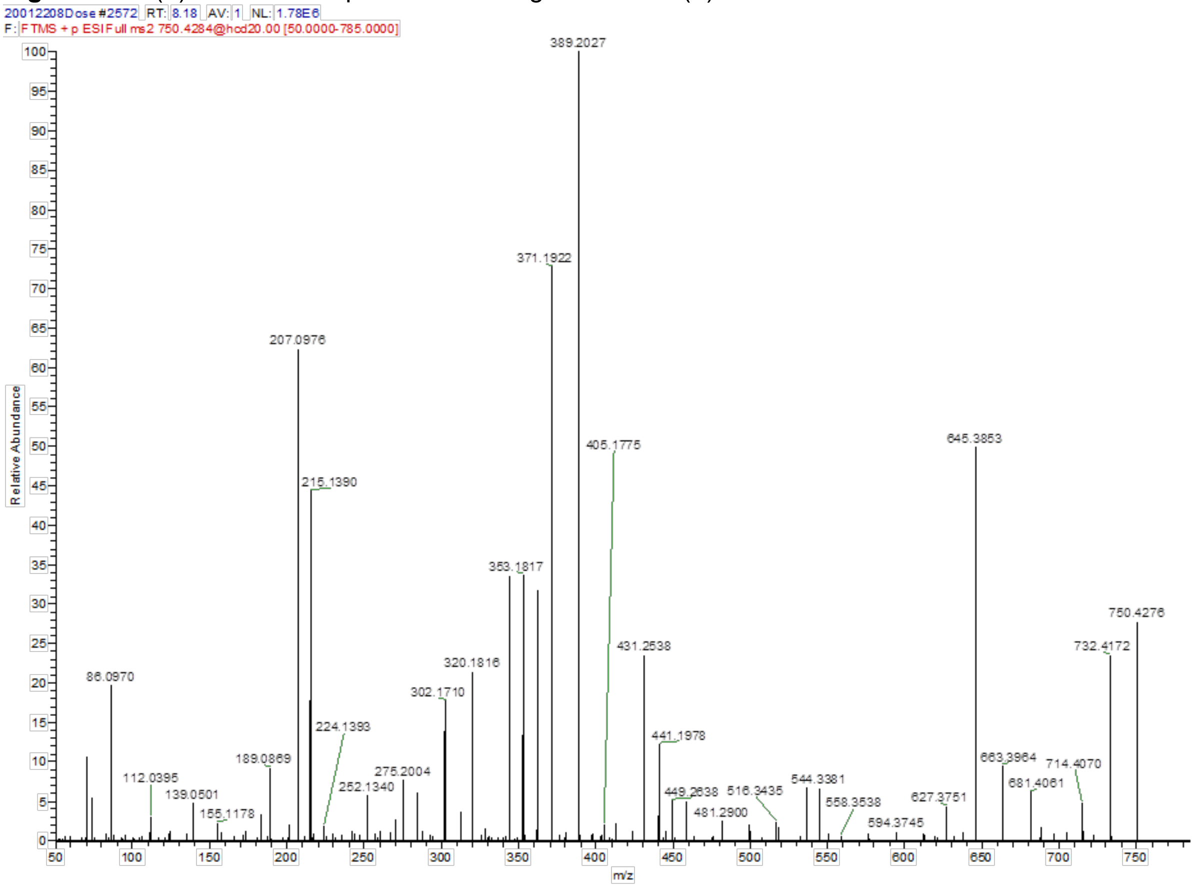


**Figure S26.** (+)ESI MS/MS spectrum of bolagladin M749 (**3**).

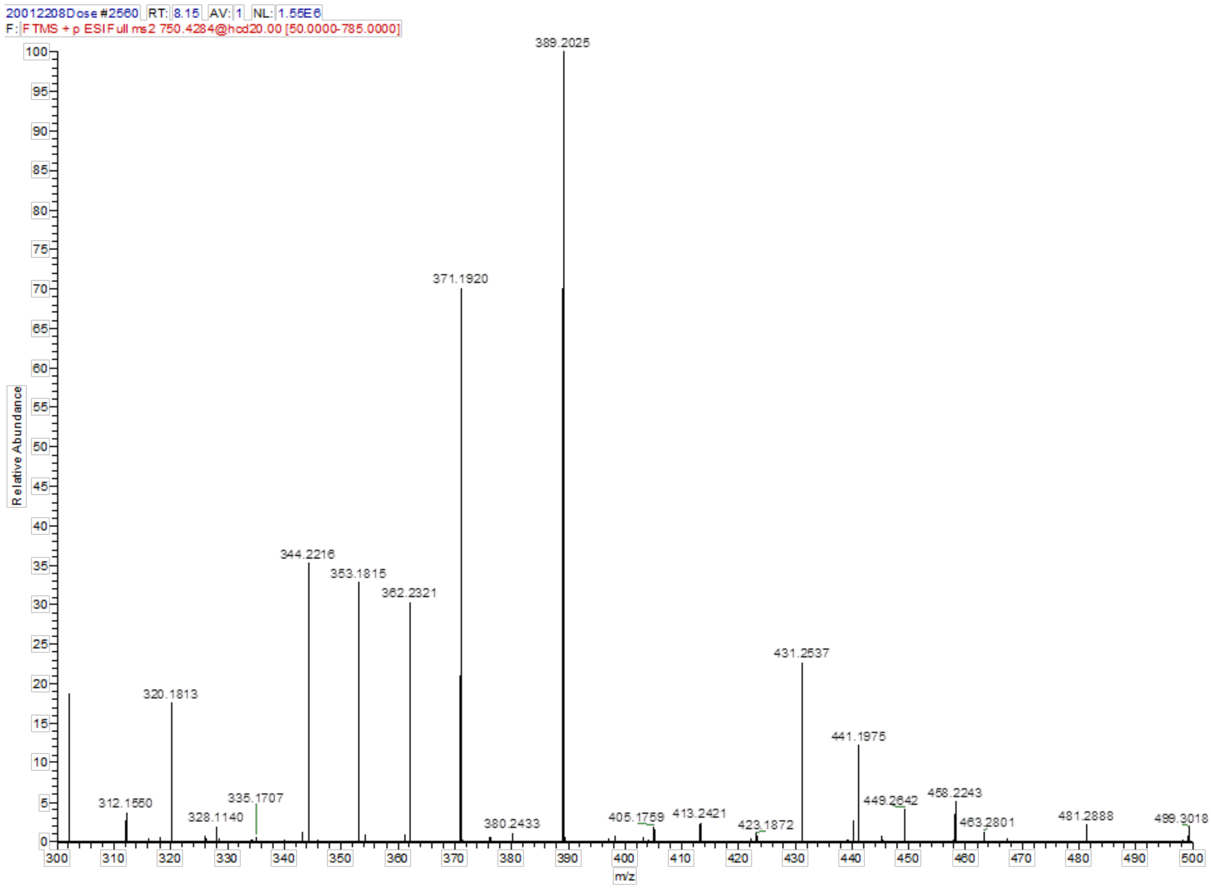


**Figure S27**. (+)ESI MS/MS spectrum of bolagladin M749 (**3**) mass range m/z 300–500.

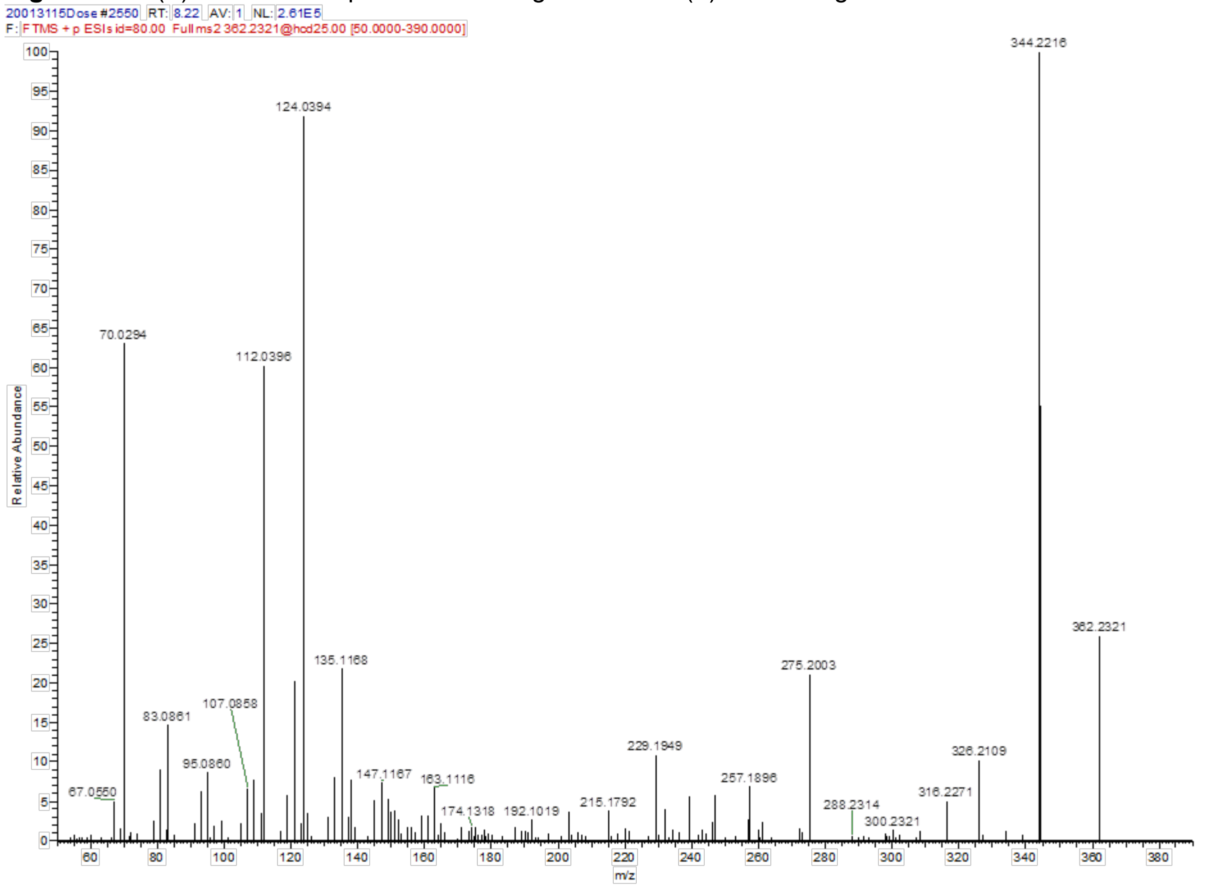


**Figure S28.** (+)ESI MS/MS spectrum of the fragment *m/z* 362 amu derived from bolagladin M749 (**3**).

**Bolagladin M765 (4)**

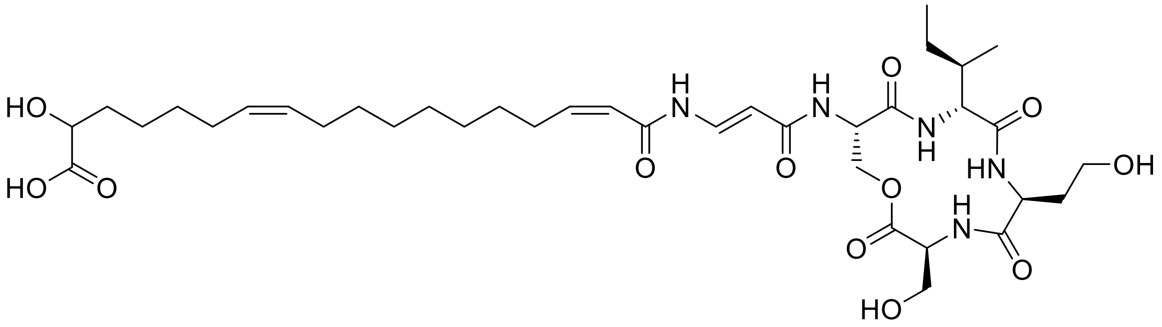

(–)ESI HR-MS: *m/z* 764.4099 [M–H]$^-$ (calcd. $C_{37}H_{58}N_5O_{12}$ 764.4087)

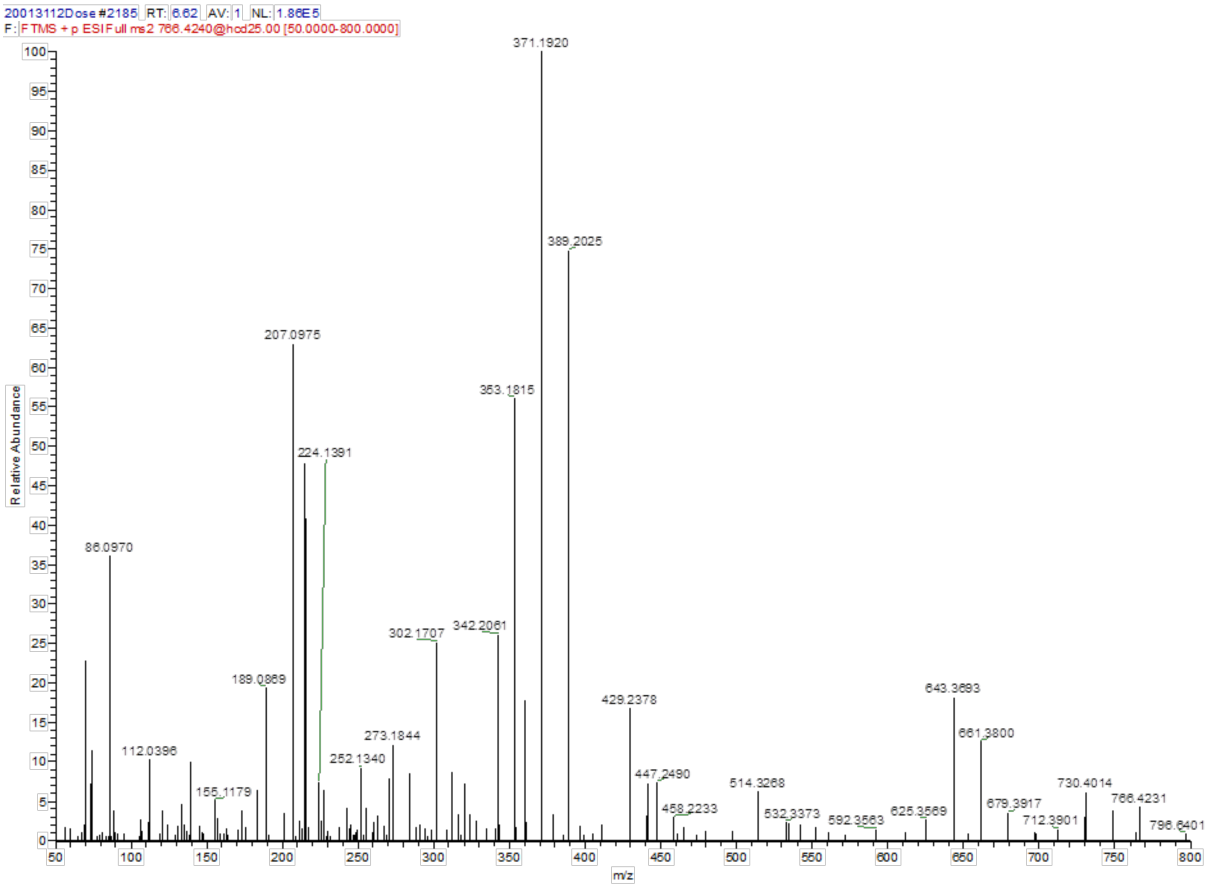


**Figure S29**. (+)ESI MS/MS spectrum of bolagladin M765 (**4**).

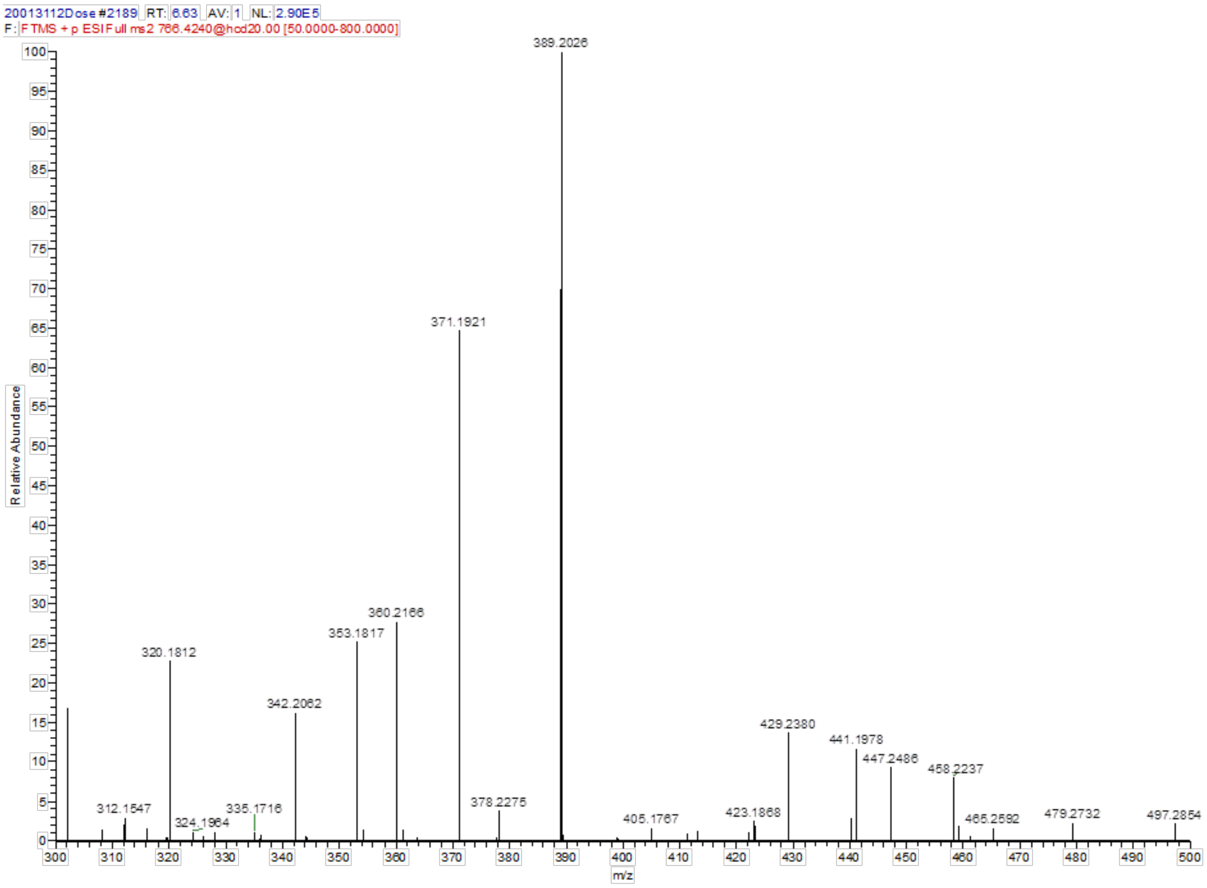


**Figure S30.** (+)ESI MS/MS spectrum of bolagladin M765 (**4**) mass range m/z 300–500.

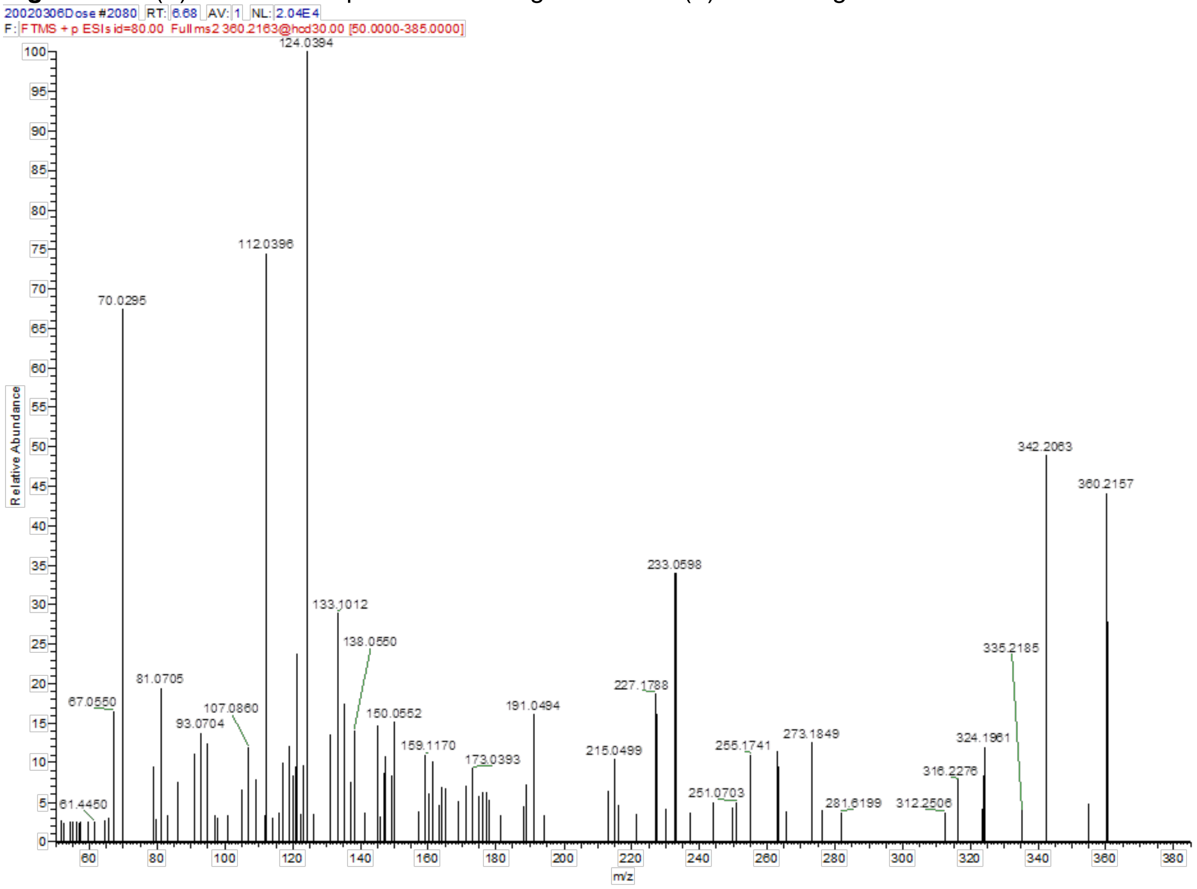


**Figure S31.** (+)ESI MS/MS spectrum of the fragment *m/z* 360 amu derived from bolagladin M765 (**4**).

### Bolagladin M839 (5)

(–)ESI HR-MS: *m/z* 838.4479 $[M–H]^-$ (calcd. $C_{40}H_{64}N_5O_{14}$, 838.4455)

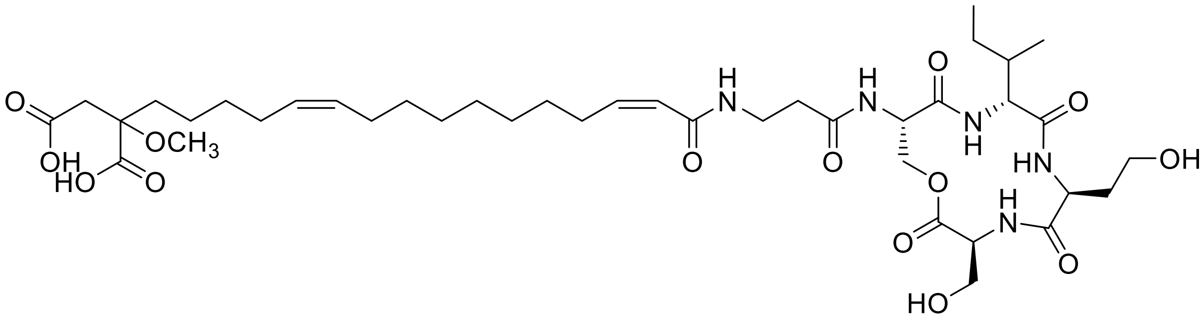


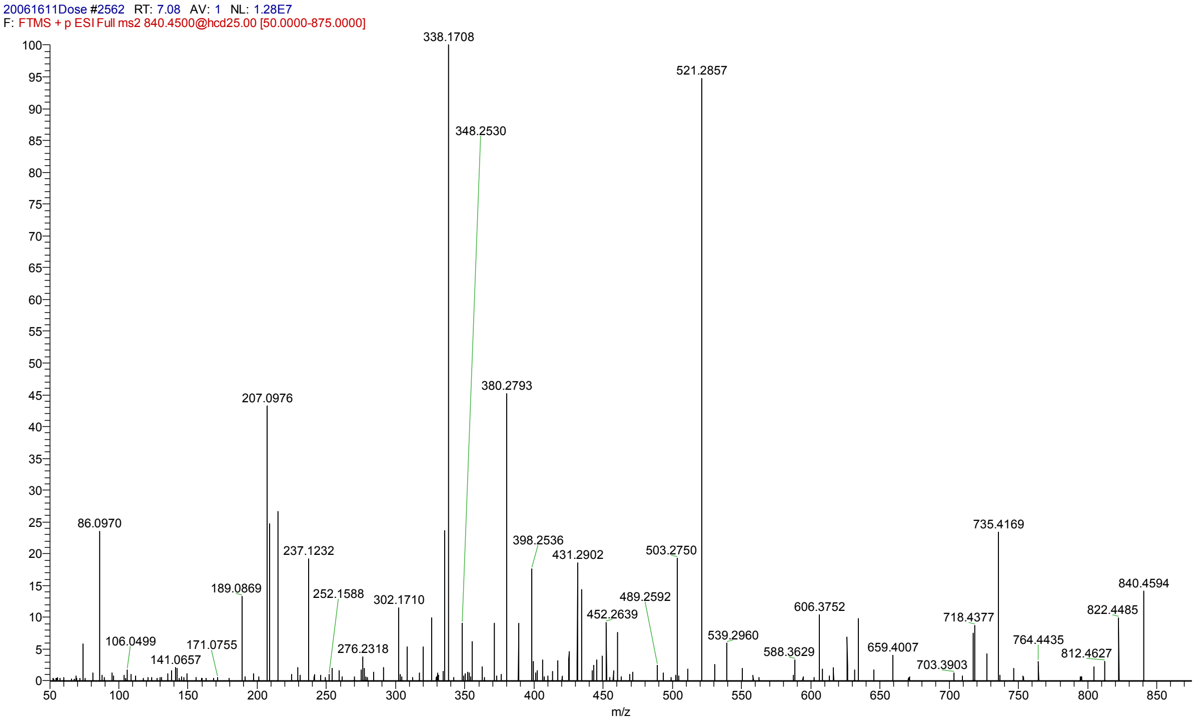


**Figure 32.** (+)ESI MS/MS spectrum of bolagladin M839 (**5**).

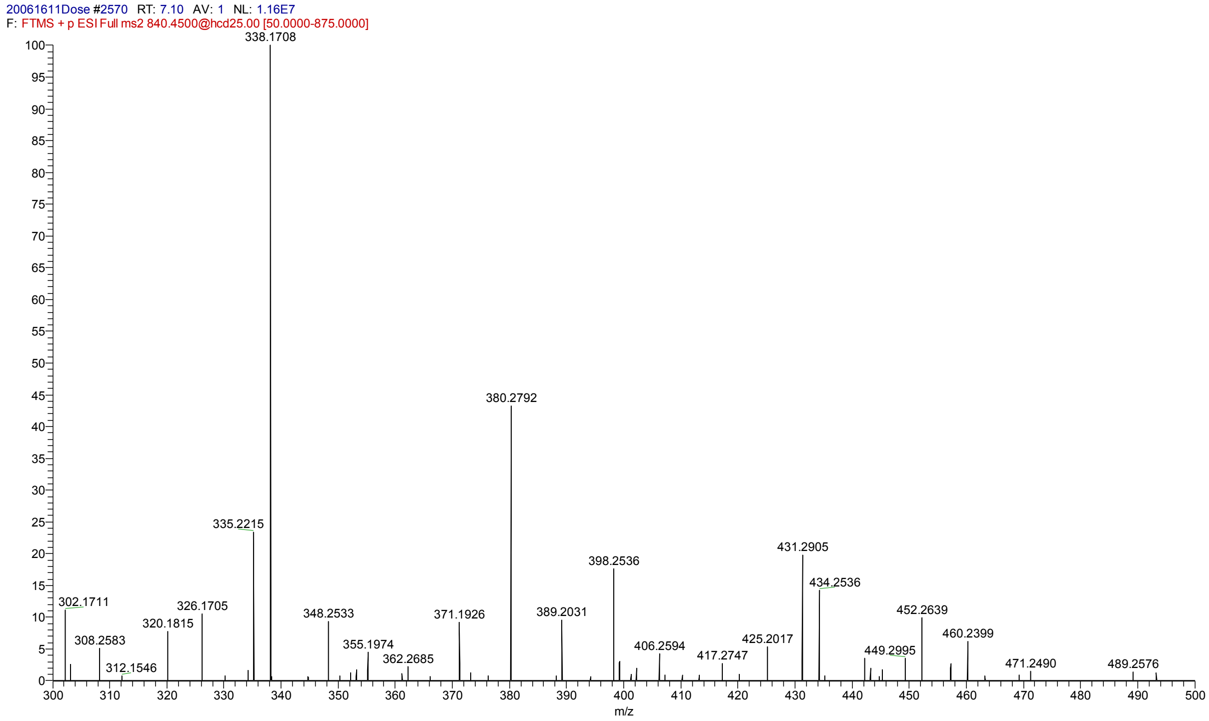


**Figure 33.** (+)ESI MS/MS spectrum of bolagladin M839 (**5**) mass range *m/z* 300–500.

### Bolagladin M824 (6)

(–)ESI HR-MS: *m/z* 824.4322 [M–H]⁻ (calcd. $C_{39}H_{62}N_5O_{14}$, 824.4299)

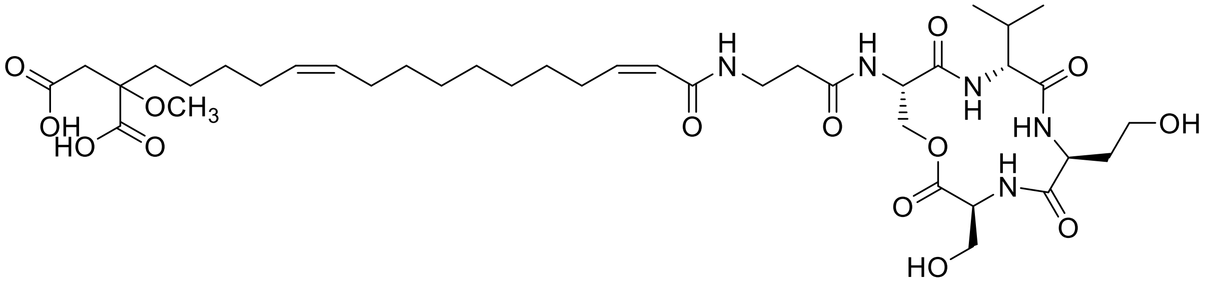

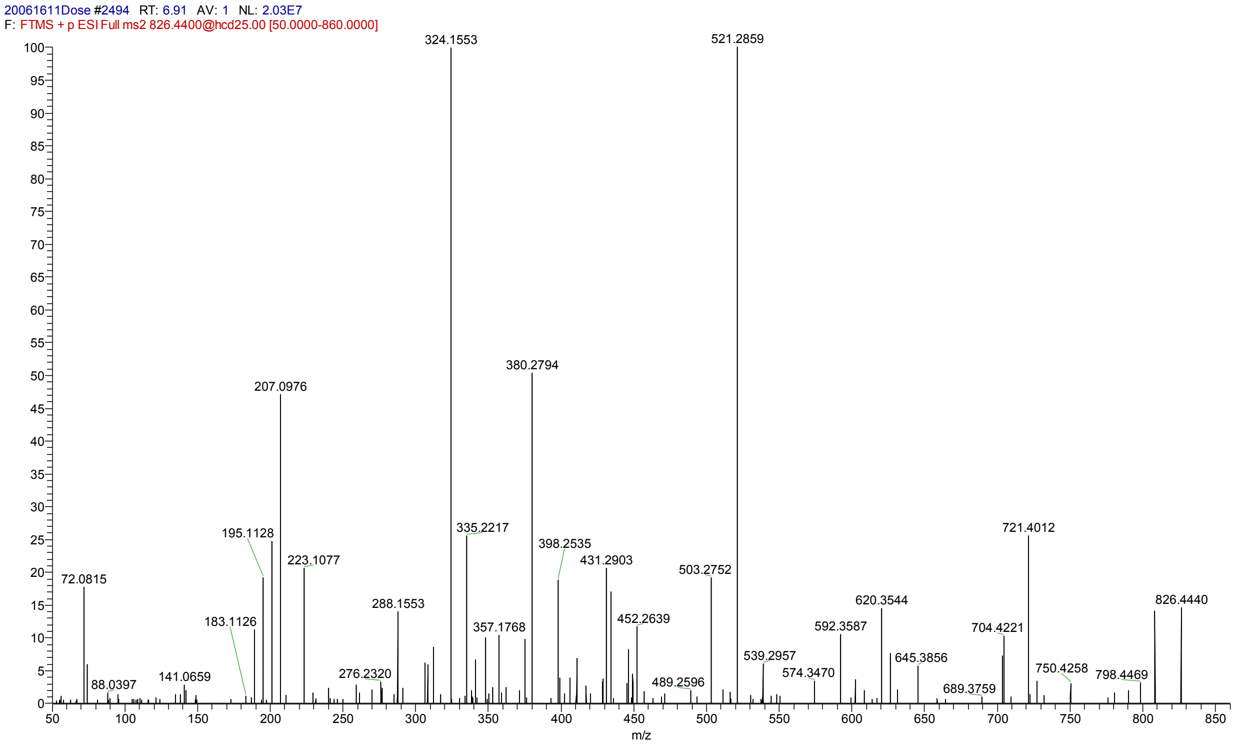


**Figure 34.** (+)ESI MS/MS spectrum of bolagladin M824 (**6**).

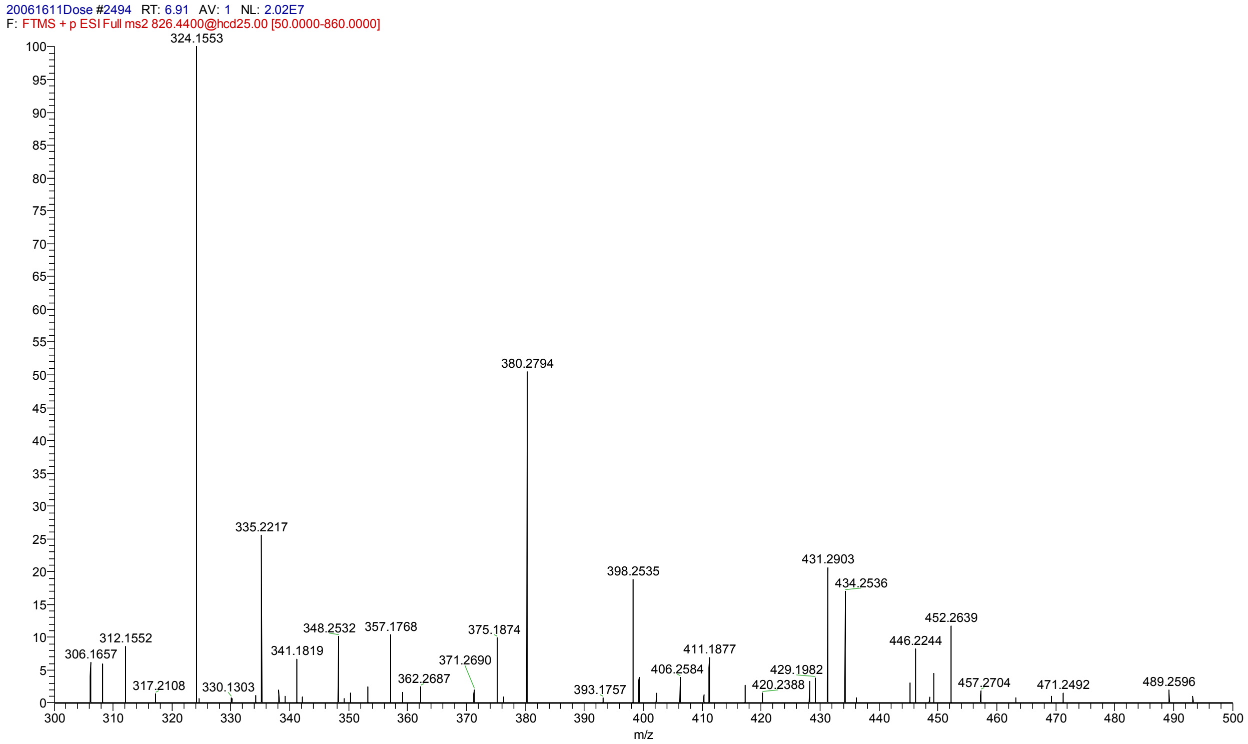


**Figure 35.** (+)ESI MS/MS spectrum of bolagladin M824 (**6**) mass range *m/z* 300–500.
